# Identification of Cell-Type-Specific Spatially Variable Genes Accounting for Excess Zeros

**DOI:** 10.1101/2021.12.27.474316

**Authors:** Jinge Yu, Xiangyu Luo

**Affiliations:** Institute of Statistics and Big Data, Renmin University of China

**Keywords:** cell-type-specific, multiple testing, spatial transcriptomics, zero-inflated negative binomial regression

## Abstract

Spatial transcriptomic techniques can profile gene expressions while retaining the spatial information, thus offering unprecedented opportunities to explore the relationship between gene expression and spatial locations. The spatial relationship may vary across cell types, but there is a lack of statistical methods to identify cell-type-specific spatially variable (SV) genes by simultaneously modeling excess zeros and cell-type proportions. We develop a statistical approach CTSV to detect cell-type-specific SV genes. CTSV directly models spatial raw count data and considers zero-inflation as well as overdispersion using a zero-inflated negative binomial distribution. It then incorporates cell-type proportions and spatial effect functions in the zero-inflated negative binomial regression framework. The R **package pscl** (Zeileis et al., 2008) is employed to fit the model. For robustness, a Cauchy combination rule is applied to integrate p-values from multiple choices of spatial effect functions. Simulation studies show that CTSV not only outperforms competing methods at the aggregated level but also achieves more power at the cell-type level. By analyzing pancreatic ductal adenocarcinoma spatial transcriptomic data, SV genes identified by CTSV reveal biological insights at the cell-type level. The R package of CTSV is available on https://github.com/jingeyu/CTSV.

## 1 Introduction

The development of spatial transcriptomic techniques has enabled the measurement of gene expression with accompanied spatial context information (Larsson et al., 2021; Zhuang, 2021; Close et al., 2021), providing unprecedented opportunities to investigate the interaction between expression and spatial locations. One crucial challenge in the spatial expression data analysis is to identify genes whose expression levels vary with spatial coordinates in a tissue section, which are termed as spatially variable (SV) genes. In recent years, the task of SV gene detection draws much attention from bioinformaticians, and several statistical methods (Edsgärd et al., 2018; Svensson et al., 2018; Sun et al., 2020; Zhu et al., 2021; Hao et al., 2021; Li et al., 2021) have been proposed to test the dependence of expression on spatial locations. However, the dependence may be confounded by some biological or technical factors. In this paper, we aim to mitigate the confounding issues in SV gene identification by accounting for two possible confounding factors—cell-type proportions and excessive zeros.

On the one hand, the commonly used spatial transcriptomics (ST) platforms, including ST based on spatially barcoded microarrays (Ståhl et al., 2016), 10x Genomics Visium (Rao et al., 2020), and Slide-seq (Rodriques et al., 2019), profile gene expression from spots that are regularly organized in a grid in a tissue section. Each spot usually consists of dozens of cells, so the observed expression measurements are at the bulk level rather than at single-cell resolution. Since spots in different tissue regions often have different cell-type proportions (Cable et al., 2021; Elosua-Bayes et al., 2021), the latent cellular compositions can induce expression variations even though the spatial locations have no impact on the expression, thus confounding the SV gene detection. In fact, the confounding issue by cell-type proportions has been also observed in other types of association studies, e.g., the epigenome-wide association studies (Zheng et al., 2018; Luo et al., 2019; Rahmani et al., 2019). On the other hand, unlike traditional bulk RNA-seq or microarray data, the bulk ST expression still suffers from zero-inflation because the expression signals for a large proportion of genes within each spot are too weak to be captured by ST technologies. Figure 1(a) shows a bar plot of spot-wise zero proportions in a real bulk ST dataset (Moncada et al., 2020), and we can observe that more than 80% of spots have at least 70% zeros in the expression. Therefore, it is necessary to account for cell-type proportions and sparsity when modeling bulk ST data.

**Fig. 1:**
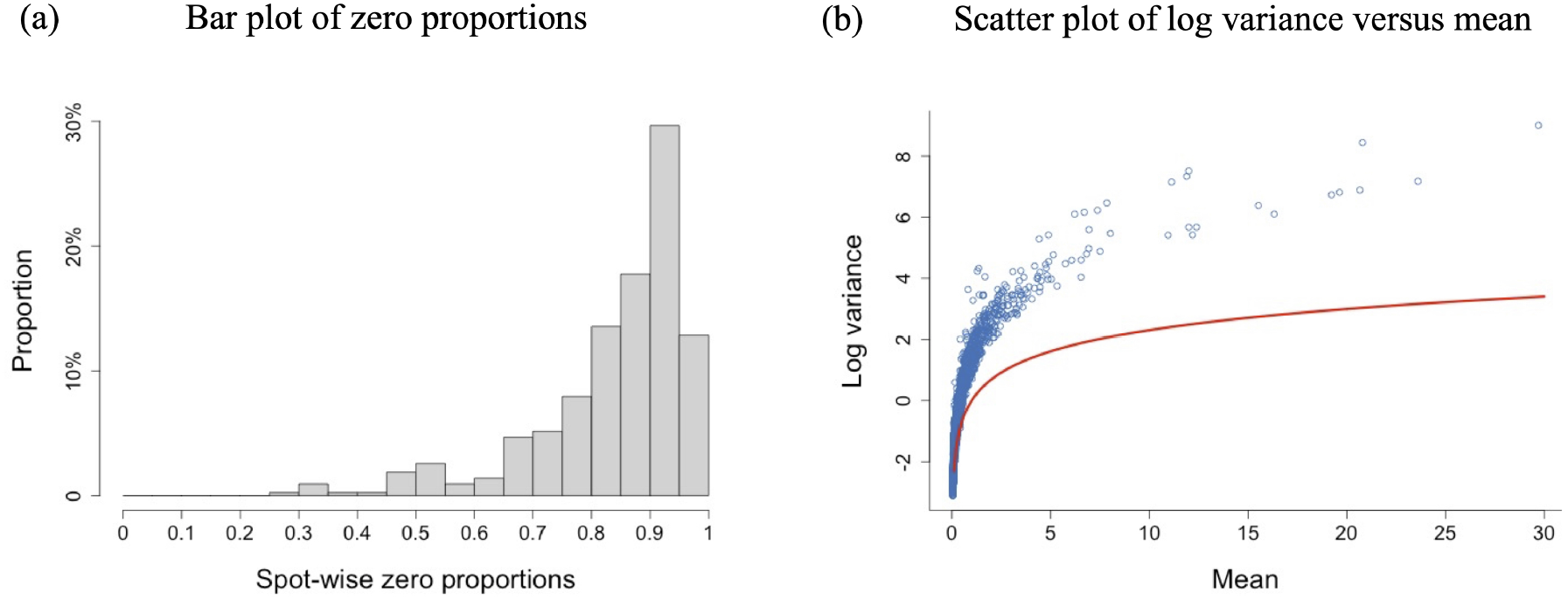
Zero-inflation and overdispersion in the pancreatic ductal adenocarcinoma (PDAC) ST data. (a) Bar plot of spot-wise zero proportions. (b) Scatter plot of genes’ expression variance at logarithmic scale versus expression mean in PDAC data. Each point corresponds to one gene, and the red curve corresponds to the points where the mean equals variance.

In bulk ST data, a gene is called **SV** if it displays an expression pattern that depends on the spatial locations of **spots** in a tissue section. Considering that a tissue consists of diverse cell types, it naturally brings in the concept of **cell-type-specific SV** genes whose expressions are affected by the spatial coordinates of **cells** in one specific cell type. However, an SV gene may not be cell-type-specific SV and vice versa. For a simple illustration, in Figure 2(a), this gene is SV across the spots, but its expression does not vary within each cell type. On the other hand, in Figure 2(b), this gene is cell-type-specific SV in both cell types 1 and 2, but its overall expressions on spots does not change. In this sense, cell-type-specific SV genes are different from SV genes, so there is a pressing need of new statistical methods to capture cell-type-specific SV genes. For common SV gene detection, frequentist methods carry out multiple hypothesis testings (non-SV in the null and SV in the alternative) and determine the p-value threshold by controlling the false discovery rate (FDR), and Bayesian methods calculate the posterior probability of being SV for each gene using posterior samples and identify SV genes based on estimated Bayesian FDR. Specifically, to our knowledge, trendsceek (Edsgärd et al., 2018) and SpatialDE (Svensson et al., 2018) are the first two statistical methods to achieve that. Trendsceek (Edsgärd et al., 2018) was built upon the marked point process to test whether the joint probability of expressions on two locations relies on their distance, calling it a mark segregation. It then makes use of four types of mark-segregation summary statistics to compute p-values through permutations. As trendsceek models the probability density, it can capture spatial expression changes both from mean and covariance. In contrast, SpatialDE (Svensson et al., 2018) only models the spatial covariance structure using zero mean Gaussian process (Williams and Rasmussen, 2006) and fits spatial expression data via a normal distribution, and then compares the result against a null model without spatial effects to calculate p-values. Recently, Hao et al. (2021) proposes SOMDE using self-organizing map to enhance the computational scalability on large-scale data. However, these methods need to first transform raw expression count data to continuous values, and this may lose power in the downstream analysis (Sun et al., 2017).

**Fig. 2:**
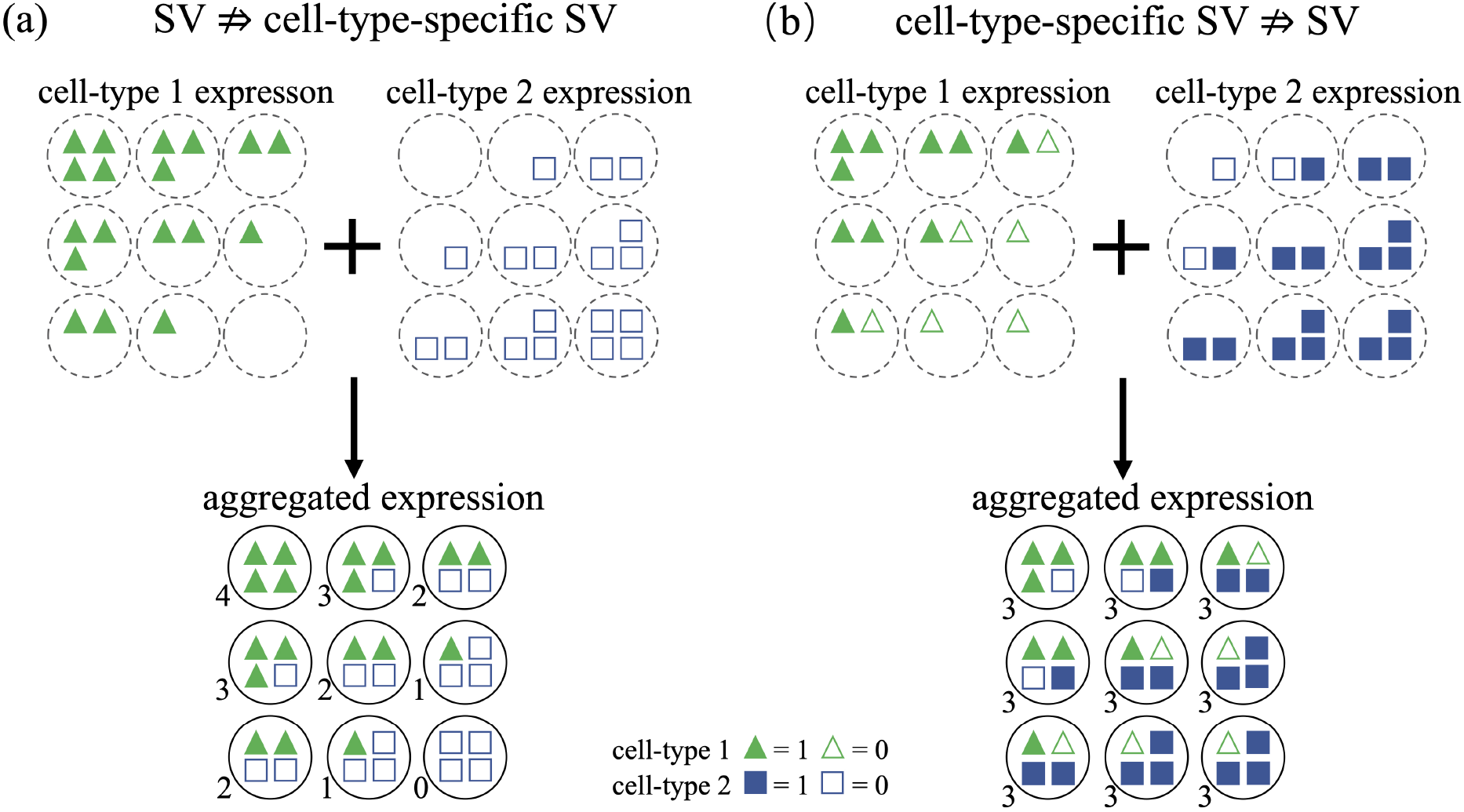
A simple illustration of SV genes and cell-type-specific SV genes. A big circle represents a spot, a solid/empty triangle means a cell of cell type 1 with expression 1/0, and a solid/empty square is a cell of cell type 2 with expression 1/0. The top panels show the gene expression distribution for each cell type, while the bottom panels display the aggregated expression pattern, where the number at the lower left of each spot is the aggregated expression value. (a) This gene is SV at the aggregated level (lower panel), but its expression keeps unchanged within each cell type (upper panel). (b) This gene is not SV at the aggregated level (lower panel), but its expression is associated with cell locations for each cell type (upper panel).

SPARK (Sun et al., 2020) is an elegant and powerful statistical method that directly fits spatial raw counts via the Poisson log linear regression model and uses the zero mean Gaussian process to model spatial effects. Hence, it can achieve more power than trendsceek and SpatialDE. It also maintains robustness by considering multiple kernel choices of the Gaussian process and combining multiple p-values through a Cauchy combination rule (Liu et al., 2019). Nevertheless, a simple Poisson distribution cannot account for excess zeros (Figure 1(a)) and overdispersion (Figure 1(b)) in the ST expression data. Recently, BOOST-GP (Li et al., 2021) explicitly models the sparse spatial expression via a zero-inflated negative binomial distribution, where the negative binomial mean is connected to covariates through a log link. Spatial effects are further incorporated via zero mean Gaussian process, and binary indicators are introduced for SV genes. Subsequently, the inference is performed in the Bayesian framework, and the posterior samples of SV gene indicators are used to calculate the posterior inclusion probability. Finally, SV genes are selected based on a controlled estimated Bayesian FDR.

Instead of the explicit modeling of zero-inflation in BOOST-GP, Zhu et al. (2021) designs a nonparametric approach SPARK-X that does not need to specify the distribution of sparse spatial expression. SPARK-X extends the scalability of SPARK and further improves its robustness on large-scale spatial transcriptomic data. Moreover, as far as we know, currently SPARK-X (Zhu et al., 2021) is the unique SV gene detection method that provides a way to identify cell-type-specific SV genes. Specifically, when applied to Slide-seq v2 data and HDST data, SPARK-X first uses the cell-type proportion estimates from RCTD (Cable et al., 2021) to assign each spot to its major cell type and then detects SV genes for spots of the same labeled cell type. Nevertheless, the assignment procedure ignores the influence of minor cell types in each spot, and thus it is more reasonable to directly utilize the cell-type proportion estimates to identify cell-type-specific SV genes.

In this paper, we develop a simple statistical approach “CTSV” to identify Cell-Type-specific SV genes accounting for excess zeros. CTSV directly fits the sparse expression raw counts using a zero-inflated negative binomial distribution, models the mean as a weighted average of cell-type-specific spatial expression profiles with weights being the cell-type proportions, and for each cell type connects the spatial expression profile to a function of spatial coordinates. By combining these equations in CTSV, the identification of cell-type-specific SV genes is equivalent to testing whether the function of spatial coordinates is zero for each cell type in a zero-inflated negative binomial regression model. Specifically, since there have been several mature bulk ST deconvolution methods (Cable et al., 2021; Elosua-Bayes et al., 2021; Dong and Yuan, 2021), we treat the estimated cell-type proportions as fixed covariates in CTSV. We further model unknown functions to be linear, focal, and periodic, respectively, and combine the p-values from the multiple choices to achieve the robustness to unavailable spatial patterns like in SPARK (Sun et al., 2020). Through simulation studies, CTSV can achieve more power than SPARK-X in detecting cell-type-specific SV genes and also outperforms other methods at the aggregated level. The real data analysis to PDAC ST data also shows the practical utility of CTSV.

The novelty of this work can be reflected in the following two main aspects. First, from the perspective of biology, our paper first introduces the concept of cell-type-specific SV genes and highlights its importance and difference from SV genes. Second, the statistical construction procedure of CTSV is novel. CTSV explicitly incorporates the cell type proportions of spots into a zero-inflated negative binomial distribution and models the spatial effects through the mean vector, whereas existing SV gene detection approaches either do not directly utilize cellular compositions or do not account for excess zeros. It is the subtle construction of CTSV that makes it possible to correctly detect cell-type-specific SV genes from bulk ST data, which will be detailed in the next section.

## 2 Method

### 2.1 The Proposed Approach CTSV

Suppose there are *G* genes, *n* spots, and *K* cell types in the tissue section. Assume that **Y** = {*Y*_*gi*_ : 1 ≤ *g* ≤ *G*, 1 ≤ *i* ≤ *n*} is the bulk ST data matrix, where *Y*_*gi*_ is the observed raw count of gene *g* in spot *i*. Let **S** = {(*s*_*i*1_, *s*_*i*2_) : 1 ≤ *i* ≤ *n*} represent the set of coordinates of spots’ centers, and ***s***_*i*_ = (*s*_*i*1_, *s*_*i*2_) is the two dimensional coordinate of spot *i*’s center. To account for the count nature and overdispersion of ST data, we consider the negative binomial distribution NB(*c*_*i*_*λ*_*gi*_, *ψ*_*g*_) with mean *c*_*i*_*λ*_*gi*_ and shape parameter *ψ*_*g*_ for gene *g* in spot *i*, and its probability mass function is 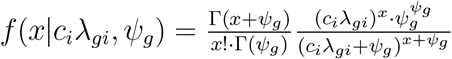 for any non-negative integer *x*. In this way, the variance equals *c*_*i*_*λ*_*gi*_ + (*c*_*i*_*λ*_*gi*_)^2^*/ψ*_*g*_ and thus is larger than the mean *c*_*i*_*λ*_*gi*_. The scalar *c*_*i*_ is a size factor to account for different library sizes of spots, and it is computed to be the ratio of spot *i*’s library size to the median library size across spots, i.e.,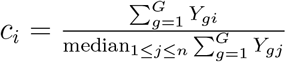.

In addition to overdispersion, bulk ST data may suffer from zero-inflation—the observed zero proportion is much larger than the expected zero proportion of a negative binomial distribution. Typically there are two kinds of zeros in the data. One is called “biological zeros” resulting from genes that do not express, and the other one is “technical zeros” or “dropout zeros,” which means that some genes have relatively low expressions but are not captured. Taking both overdispersion and zero-inflation into consideration, we model the count data *Y*_*gi*_ by a zero-inflated negative binomial distribution,

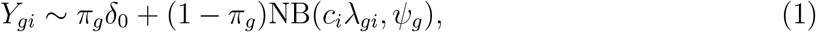

where *π*_*g*_ denotes the probability of being a technical/dropout zero for gene *g* in the spots and *δ*_0_ is a Dirac measure with point mass at zero.

As one spot may consist of dozens of heterogeneous cells, we model the log scale of *λ*_*gi*_ as a mix of cell-type-specific relative expression levels of gene *g* in spot *i*,

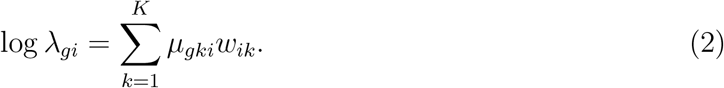

*w*_*ik*_ is the cell-type *k* proportion in spot *i*, and *µ*_*gki*_ represents the relative mean expression level of gene *g* for cell type *k* in spot *i. µ*_*gki*_ depends on the spot *i* through its location **s**_*i*_, and the relationship is modeled as follows using a similar formulation from Luo et al. (2019).

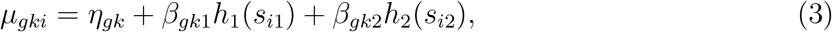

where *η*_*gk*_ is the cell-type-*k* baseline expression level of gene *g*, the two functions *h*_1_(·) and *h*_2_(·) describe the spatial effects on the mean *η*_*gk*_, and the coefficients *β*_*gk*1_ and *β*_*gk*2_ are of our interest that can reflect whether the location **s**_*i*_ affects the expression of gene *g* in cell type *k*. Subsequently, by combining Equations (1)-(3), we arrive at the proposed approach CTSV (Cell-Type-specific Spatially Variable gene detection),

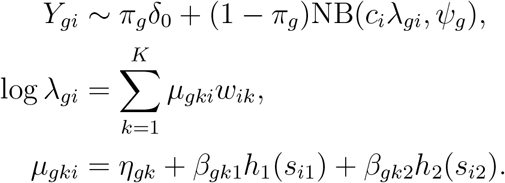

If we integrate the last two equations, CTSV is equivalent to

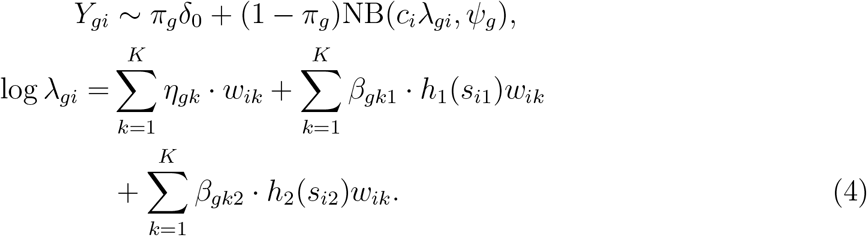

Our next goal is to conduct statistical inference for the coefficients *β*_*gk*1_ and *β*_*gk*2_ to test whether they are zero or not for each gene. Specifically, if at least one of the two null hypotheses *H*_0_ : *β*_*gk*1_ = 0 and *H*_0_ : *β*_*gk*2_ = 0 is rejected, then we believe that gene *g* is SV in cell type *k*.

### 2.2 Statistical Inference

#### 2.2.1 When functions *h*_1_ and *h*_2_ are known

In Equation (4), if we know the cellular compositions {*w*_*ik*_ : *k* = 1, …, *K*} for each spot *i* as well as the functions *h*_1_ and *h*_2_, then we can treat them as covariates and thus the inference for CTSV reduces to the inference for a zero-inflated negative binomial regression model (Preisser et al., 2016), which can be easily conducted by the R package **pscl** (Zeileis et al., 2008). However, the cellular compositions of each spot are often unavailable. Fortunately, there have been several deconvolution methods designed for bulk ST data recently, such as RCTD (Cable et al., 2021), SPOTlight (Elosua-Bayes et al., 2021), and SpatialDWLS (Dong and Yuan, 2021). Subsequently, we treat the estimates for {*w*_*ik*_ : *k* = 1, …, *K*} as fixed covariates and plug them in Equation (4).

The parameter estimation in the zero-inflated negative binomial distribution is not trivial. For example, Miao et al. (2018) used EM algorithm (Dempster et al., 1977) to estimate the dropout zero probability *π*_*g*_. For each gene, CTSV is essentially a zero-inflated negative binomial regression model, and the likelihood function can be written as follows (gene index *g* is suppressed for simplicity).

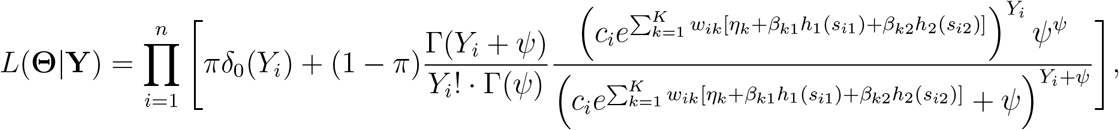

where the parameter set 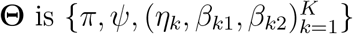. We follow the estimation strategy from Zeileis et al. (2008) to obtain approximated maximum likelihood estimates for **Θ**. Specifically, we utilize the conjugate gradient algorithm (Gilbert and Nocedal, 1992) to minimize the negative logarithmic likelihood (− log *L*(**Θ**|**Y**)) with warm starting values being the iteratively reweighted least squares estimates (Green, 1984).

Next, based on the R package pscl (Zeileis et al., 2008), we can obtain the p-value *p*_*gkℓ*_ for the hypothesis *H*_0_ : *β*_*gkℓ*_ = 0 vs *H*_1_ : *β*_*gkℓ*_ ≠ = 0 for gene *g* in cell type *k* along the *ℓ*-th coordinate (*ℓ* = 1, 2) via Wald tests. Notice that as the inference is carried out for each gene independently, the procedure is highly parallelable. We also remark that the usage of pscl is just a computational tool to realize the statistical inference for regression coefficients ***β*** in CTSV, which does not damage the novelty of CTSV. All the p-values can be organized into a p-value matrix {*p*_*gkℓ*_} with dimension *G×* 2*K*, where the *k*-th (1 ≤ *k* ≤ *K*) column corresponds to the p-value vector in cell type *k* for the *s*_1_ coordinate and the (*K* + *k*)-th (1 ≤ *k* ≤ *K*) column to the p-value vector in cell type *k* for the *s*_2_ coordinate. To control the false discovery rate (FDR) in the multiple hypothesis testings, we convert the p-value matrix to the q-value matrix {*q*_*gkℓ*_}_*G×*2*K*_ using the R package **qvalue** (Storey and Tibshirani, 2003; Storey et al., 2020). In this way, a q-value threshold *α* controls the false discovery rate to be not larger than *α*.

Specifically, for each *g*-th row in the q-value matrix, if there is at least one q-value in this row (*q*_*gkℓ*_ : 1 ≤ *k* ≤ *K, ℓ* = 1, 2) less than *α*, we call the corresponding gene *g* SV at the aggregated level. For each cell type *k*, if there is at least one q-value in (*q*_*gkℓ*_ : *ℓ* = 1, 2) less than *α*, we then identify the gene *g* to be cell-type-*k*-specific SV.

#### 2.2.2 When functions *h*_1_ and *h*_2_ are unknown

In practice, we often do not know what the type of underlying spatial patterns is in the tissue section for each gene. To deal with possible model misspecification and make the CTSV method more robust, we follow the idea from Sun et al. (2020) to choose three types of functions for *h*_1_ and *h*_2_, which can reflect the linear, focal, and periodic spatial expression patterns. Specifically, suppose that **s**_1_ and **s**_2_ are first transformed to have mean zero and standard deviation one. We choose linear functions as *h*_1_(*s*_*i*1_) = *s*_*i*1_ and *h*_2_(*s*_*i*2_) = *s*_*i*2_, squared exponential functions 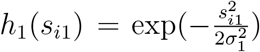 and 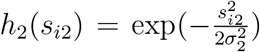, and periodic functions 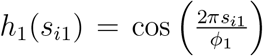 and 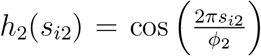. Moreover, for the squared exponential functions, we choose two sets of scale length parameters by (i) letting *σ*_1_ and *σ*_2_ be the 40% quantile of the absolute values of the transformed *s*_*i*1_ and *s*_*i*2_, respectively, denoted by *σ*_1_ = *Q*_40%_(|**s**_1_|), *σ*_2_ = *Q*_40%_(|**s**_2_|); and (ii) letting *σ*_1_ = *Q*_60%_(|**s**_1_|), *σ*_2_ = *Q*_60%_(|**s**_2_|). Similarly, for periodic functions, we set (i) *ϕ*_1_ = *Q*_40%_(|**s**_1_|), *ϕ*_2_ = *Q*_40%_(|**s**_2_|) and (ii) *ϕ*_1_ = *Q*_60%_(|**s**_1_|), *ϕ*_2_ = *Q*_60%_(|**s**_2_|). Hence, for each gene *g* in cell type *k* along *ℓ*-th coordinate, we obtain five p-values.

Accordingly, for gene *g* in cell type *k* along *ℓ*-th coordinate, we combine the five p-values 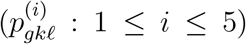 following the Cauchy combination rule ACAT (Liu et al., 2019). We first convert each of the five p-values into a Cauchy statistic 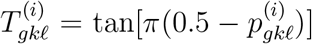, then take an average of them 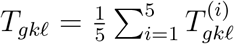, and transform the average into a single p-value *p*_*gkℓ*_ = ℙ(*C* ≥ *T*_*gkℓ*_), where *C* follows the standard Cauchy distribution (Liu et al., 2019; Pillai and Meng, 2016). In this way, we convert five p-value matrices to one p-value matrix (*p*_*gkℓ*_)_*G×*2*K*_, and then the inference is based on the FDR control as discussed before.

## 3 Simulation

In this section, we compared the performance of our method with several state-of-the-art SV gene detection methods. We generated the spatial transcriptomic raw count data following Equation (4), where related parameters are set as follows. Suppose there are *G* = 10, 000 genes, *n* = 600 spots, and *K* = 6 cell types. The cell-type-*k* baseline expression profile ***η***_*k*_ was generated from normal distributions. Specifically, we first independently simulated *η*_*g*1_ from N(2, 0.2^2^) for *g* = 1, …, *G* in cell type 1 and then randomly sampled 300 differentially expressed (DE) genes for each cell type *k* (2 ≤ *k* ≤ *K*). Next, on the cell-type-*k* DE genes (2 ≤ *k* ≤ *K*), we sampled *η*_*gk*_ from 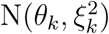 independently, where (*θ*_2_, *ξ*_2_) = (3, 0.2), (*θ*_3_, *ξ*_3_) = (2, 0.2), (*θ*_4_, *ξ*_4_) = (4, 0.2), (*θ*_5_, *ξ*_5_) = (3, 0.2), (*θ*_6_, *ξ*_6_) = (4, 0.2). For expressions on the remaining genes, we set *η*_*gk*_ = *η*_*g*1_. The explanations for the parameter choices are given in Supplementary Section S1. Moreover, we partitioned the spot region into four regions as displayed in Figure 3(a) and then sampled cell-type proportions ***w***_*i*_ of spot *i* from Dirichlet distributions. Cell-type proportions of spots in regions from 1 to 4 were independently sampled from Dir(1, 1, 1, 1, 1, 1), Dir(1, 3, 5, 7, 9, 11), Dir(16, 14, 12, 10, 8, 6), and Dir(1, 4, 4, 4, 4, 1), respectively. For coefficients *β*_*gk*_, we set 200 SV genes in each cell type, and there were 700 SV genes at the aggregated level. Figure 3(b) shows the SV gene distribution patterns in each cell type. We further consider the following three simulation settings to specify the spatial effects *h*_1_ and *h*_2_.

**Fig. 3:**
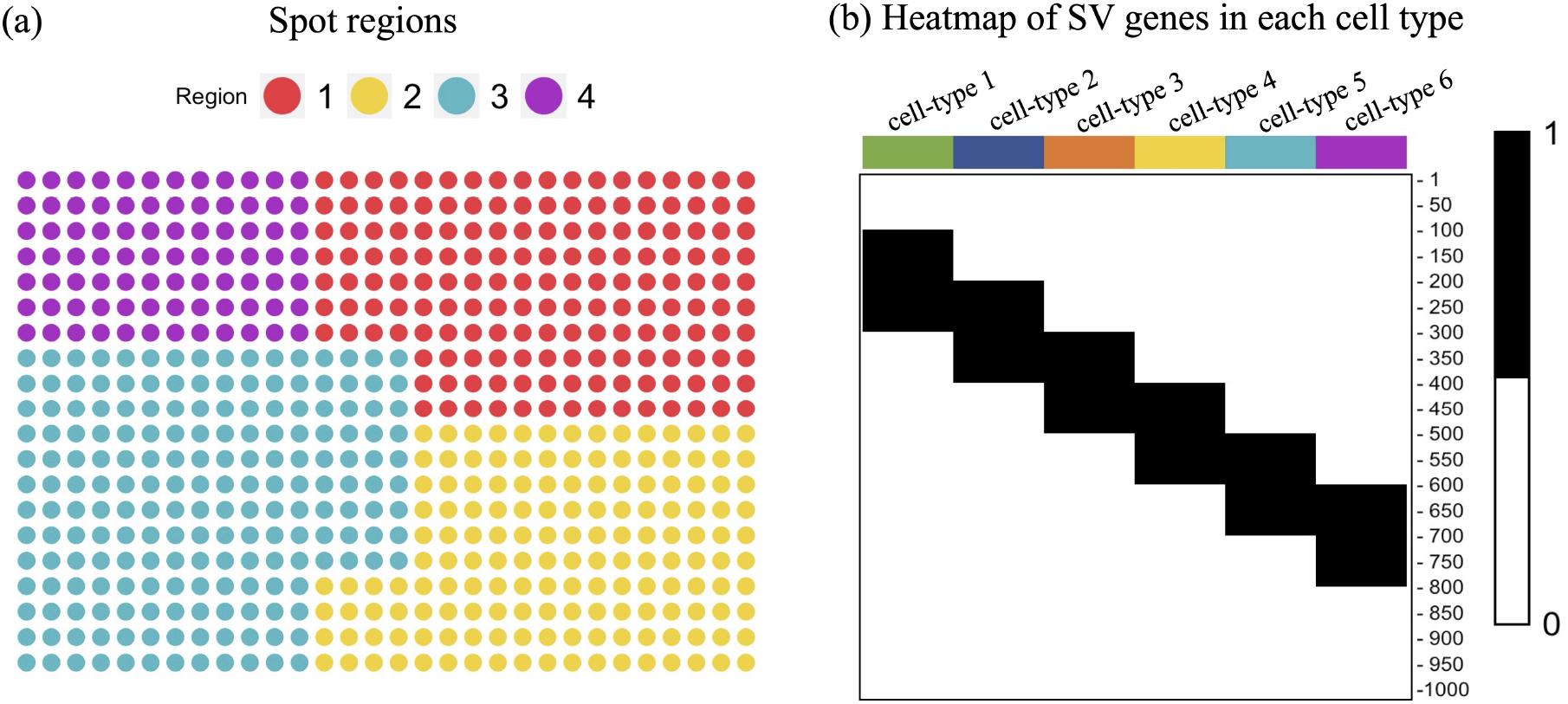
Spot regions and the heatmap of cell-type-specific SV gene pattern. (a) Four spot regions with different colors. (b) Heatmap of the SV gene pattern. If one gene in a cell type is SV, then it is colored by black. Only the first 1,000 genes are shown for a good visualization because all the remaining genes are not SV.

1. For the linear spatial pattern as shown in Figure 4(a), we chose *h*_1_(*s*_*i*1_) = *s*_*i*1_ and *h*_2_(*s*_*i*2_) = *s*_*i*2_. For SV genes, we set *β*_*gk*1_ = 1.8 and *β*_*gk*2_ = 0.8 for each cell type. For non SV genes, *β*_*gkℓ*_ was set to be zero.

**Fig. 4:**
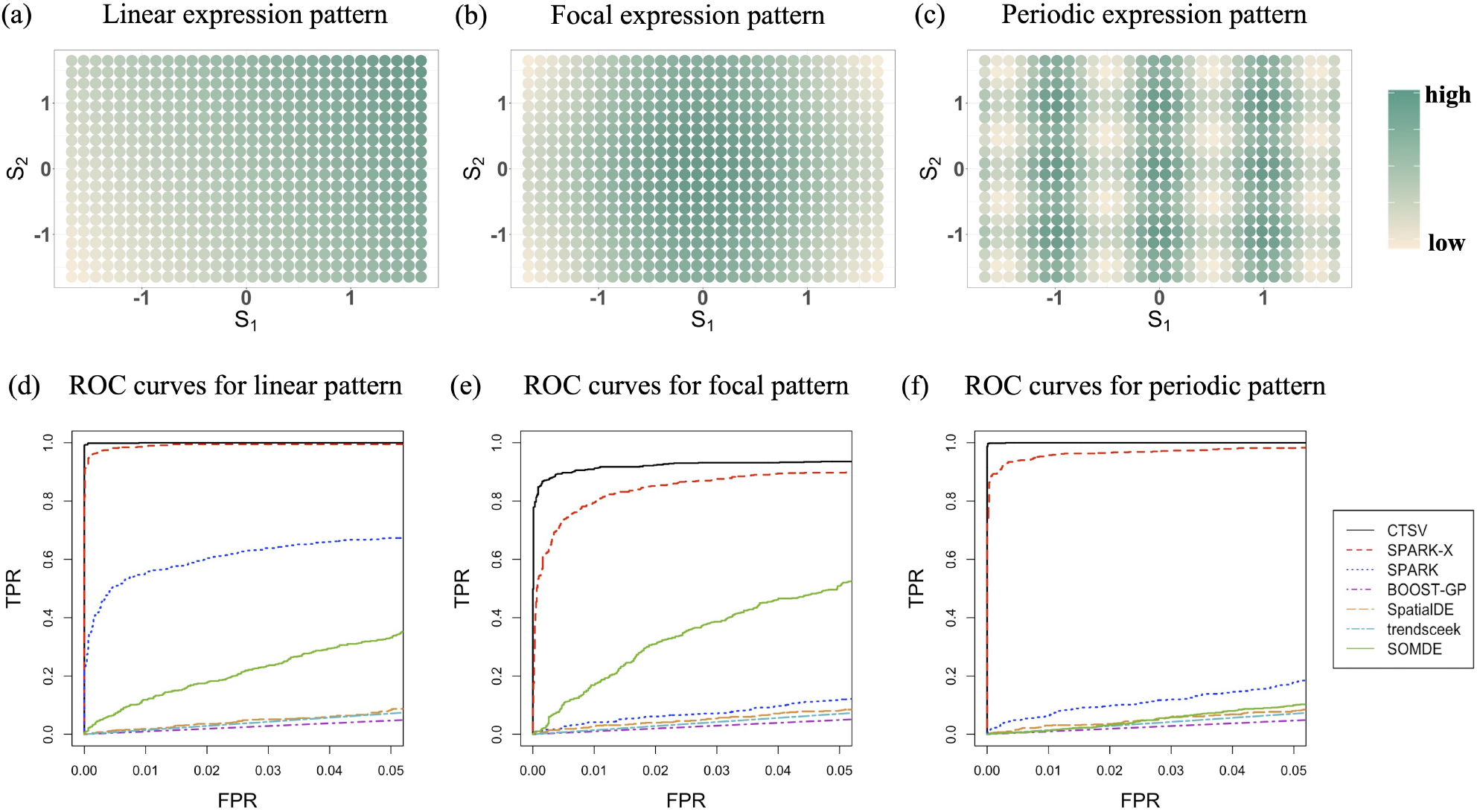
SV genes’ spatial expressions in (a) linear pattern, (b) focal pattern, and (c) periodic pattern, where the coordinates are scaled to have mean zero and standard deviation one. (d-f) The ROC curves with FPR controlled to be less than 0.05 for CTSV, SPARK-X, SPARK, BOOST-GP, SpatialDE, trendsceek, and SOMDE in the three spatial expression patterns. Only the FPR range (0, 0.05) is shown because in medical and clinical practice we often need to control FPR to be less than a threshold and some range of thresholds may be more important than others (Pencina et al., 2008). To provide more information, the ROC curves over the whole FPR range (0, 1) are also given in the Supplementary Figure S2.
2. For the focal spatial pattern as shown in Figure 4(b), we set 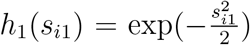 and 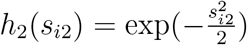. For SV genes in each cell type, we set *β* = 3 and *β* = 1. For non SV genes, *β*_*gkℓ*_ was set to be zero.
3. For the periodic spatial pattern as shown in Figure 4(c), we have *h*_1_(*s*_*i*1_) = cos (2*πs*_*i*1_), *h*_2_(*s*_*i*2_) = cos (2*πs*_*i*2_). For SV genes in each cell type, we set *β*_*gk*1_ = 2.5 and *β*_*gk*2_ = 1. For non SV genes, *β*_*gkℓ*_ was set to be zero.

After obtaining ***η***_*k*_, ***w***_*i*_, *h*_1_(*s*_*i*1_), *h*_2_(*s*_*i*2_), and *β*_*gkℓ*_, we can calculate log *λ*_*gi*_ and then sample *Y*_*gi*_ from NB(*c*_*i*_*λ*_*gi*_, *ψ*_*g*_), where the shape parameter is *ψ*_*g*_ = 100 and *c*_*i*_ = 1. Considering ST data have a large proportion of zeros, we set *π*_*g*_ (*g* = 1, …, *G*) to be 0.6 in each spatial pattern. Therefore, for each gene, the count data was set to be dropout zero with a probability 0.6. Subsequently, we applied the proposed method CTSV to the three types of simulated ST data and compared the performance with trendsceek (Edsgärd et al., 2018), SpatialDE (Svensson et al., 2018), SPARK (Sun et al., 2020), SPARK-X (Zhu et al., 2021), BOOST-GP (Li et al., 2021), and SOMDE (Hao et al., 2021). Their implementation details are given in Supplementary Section S2.

When implementing CTSV, we considered the estimate error for the cell-type proportions and sampled 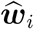 from Dir(*α*_0_***w***_*i*_) with *α*_0_ = 100. In addition, if NA (Not Available) is returned by the function *zeroinfl* in R package **pscl** (Zeileis et al., 2008), the corresponding p-value is recorded as one. In the argument of function *zeroinfl*, some commonly used optimization methods can be used, such as BFGS, conjugate gradient (CG), or Nelder-Mead, and we applied CG algorithm for its stability during the optimization procedure. We displayed the histogram for the absolute estimation error 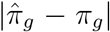 in Supplementary Figure S1, showing that the estimation errors concentrate on very small values. Hence, the estimation for the dropout zero probability has slight effects on the detection of cell-type-specific SV genes.

The receiver operating characteristic (ROC) curves for identifying SV genes at the aggregated level in the three simulation settings were reported in Figure 4(d)-(f), respectively, where the false positive rate (FPR) is controlled to be less than 0.05 for a good visualization of the performance comparison. The partial ROC curves indicate that CTSV uniformly outperformed other methods in SV gene detection at the aggregated level. In each setting, the performance of CTSV was followed by SPARK-X, which also performs well due to its nonparametric nature. SPARK ranked the third for the linear and periodic settings, while SOMDE ranked the third in the focal spatial pattern. SpatialDE, trendsceek, and BOOST-GP fail to achieve enough power in all the three simulation settings. Note that trendsceek has four types of statistics, and we only showed the best one. When controlling the FDR less than 0.01 for each method (i.e., the q-value threshold is 0.01), Table 1 demonstrates the true positive rates (TPR) and the number of false positives (FP) in the three spatial expression patterns for all the methods. CTSV and SPARK-X gave much higher TPR than other methods, while the FP of CTSV was slightly larger than SPARK-X. We also observed that trendsceek, SpatialDE, and SOMDE cannot identify any SV gene with FDR less than 0.01. Therefore, at the aggregated level, CTSV can provide a high power with controlled FP and FDR owing to its ability to handle excess zeros and account for cell-type proportions.

**Table 1:** The comparisons of true positive rate (TPR) and the number of false positives (FP) in SV gene detection at the aggregated level.

|  | Pattern | CTSV | SPARK-X | SPARK | BOOST-GP | SpatialDE | SOMDE | trendsceek |
| --- | --- | --- | --- | --- | --- | --- | --- | --- |
| TPR | Linear | 0.999 | 0.907 | 0.178 | 0.001 | 0 | 0 | 0 |
|  | Focal | 0.871 | 0.293 | 0 | 0.001 | 0 | 0 | 0 |
|  | Periodic | 0.999 | 0.819 | 0 | 0.003 | 0 | 0 | 0 |
| FP | Linear | 33 | 1 | 0 | 5 | 0 | 0 | 0 |
|  | Focal | 19 | 3 | 0 | 5 | 0 | 0 | 0 |
|  | Periodic | 21 | 3 | 0 | 4 | 0 | 0 | 0 |

Regarding the detection of cell-type-specific SV genes, as SPARK-X is currently the only method that can achieve the function, we compared CTSV and SPARK-X. In SPARK-X (Zhu et al., 2021), if one spot was dominated by a cell type, which has the maximal proportion in that spot, SPARK-X assigned the spot to the cell type. Subsequently, SPARK-X performed the detection task on spots with the same cell type. Figure 5 displays the heatmaps of −log_10_(*p*_*gk*_) (*g* = 1, …, 1000) of CTSV and SPARK-X, where *p*_*gk*_ is the p-value of gene *g* in cell-type *k* for SPARK-X, and *p*_*gk*_ = min(*p*_*gk*1_, *p*_*gk*2_) for CTSV. The darker the color, the more significant that the corresponding gene is SV in that cell type. Compared with the underlying truth (Figure 3(b)), CTSV obtained more accurate results in identifying cell-type-specific SV genes than SPARK-X. Table 2 indicates that when FDR is controlled to be less than 0.01, CTSV yielded higher power than SPARK-X for all the cell types in the three simulation settings, but CTSV did not perform very well in the focal spatial expression pattern. The results showed that CTSV is good at identifying cell-type-specific SV genes by directly modeling cell-type proportions rather than transforming them to one-hot code like in SPARK-X, which may lose some information.

**Table 2:** Cell-type-specific TPR and FP of CTSV and SPARK-X.

|  | Pattern | Linear |  | Focal |  | Periodic |  |
| --- | --- | --- | --- | --- | --- | --- | --- |
|  | Methods | CTSV | SPARK-X | CTSV | SPARK-X | CTSV | SPARK-X |
| TPR | cell-type 1 | 1 | 0.375 | 0.800 | 0 | 1 | 0 |
|  | cell-type 2 | 0.995 | 0.095 | 0.785 | 0 | 0.940 | 0 |
|  | cell-type 3 | 0.980 | 0 | 0.605 | 0 | 0.970 | 0 |
|  | cell-type 4 | 0.975 | 0 | 0.515 | 0 | 0.980 | 0 |
|  | cell-type 5 | 0.995 | 0 | 0.780 | 0 | 0.970 | 0 |
|  | cell-type 6 | 0.995 | 0 | 0.905 | 0 | 0.995 | 0 |
| FP | cell-type 1 | 35 | 33 | 9 | 0 | 1 | 0 |
|  | cell-type 2 | 22 | 4 | 9 | 0 | 1 | 0 |
|  | cell-type 3 | 10 | 0 | 5 | 0 | 7 | 0 |
|  | cell-type 4 | 11 | 0 | 8 | 0 | 6 | 0 |
|  | cell-type 5 | 3 | 0 | 9 | 0 | 3 | 0 |
|  | cell-type 6 | 21 | 0 | 119 | 0 | 5 | 0 |

**Fig. 5:**
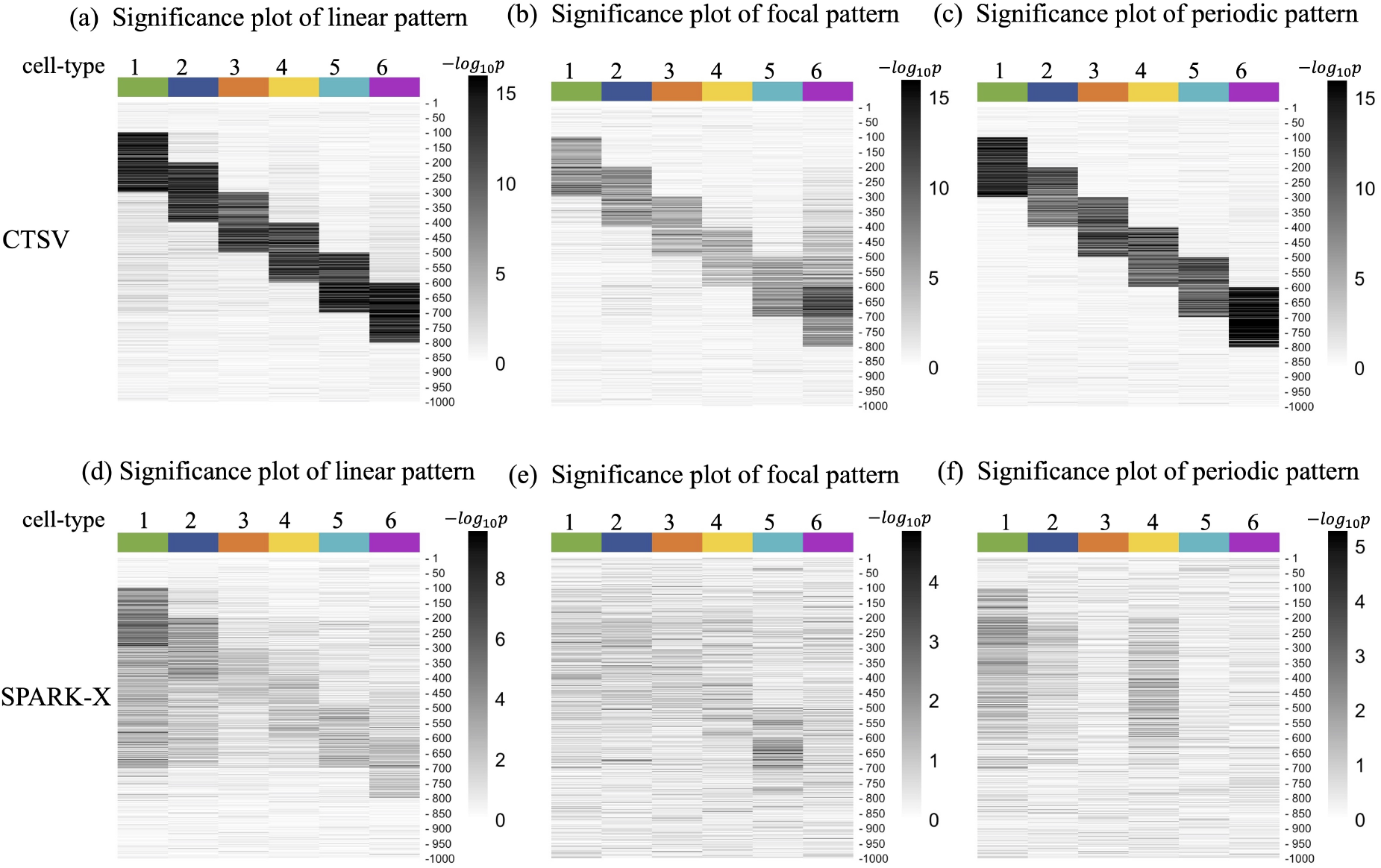
(a-c) Significance plots of CTSV and (d-f) significance plots of SPARK-X in the three spatial expression patterns for the first 1,000 genes. Values in the heatmaps are −*log*_10_*p* of the corresponding gene in each cell type. The darker the color, the more likely the corresponding gene is to be SV in that cell type.

### Imperfect deconvolution

To explore the effects of imperfect cell-type proportion estimations on the SV gene discovery, we performed additional experiments under the three spatial patterns. When implementing CTSV, we sampled the cell-type proportion estimates for each spot 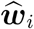 from Dir(*α*_0_***w***_*i*_) (***w***_*i*_ is the underlying truth) with the concentration parameter *α*_0_ = 100, 80, 60, 40, 20, 10, 5, 1, respectively. The lower the *α*_0_, the less accurate the deconvolution estimates. Supplementary Figure S3 show that the imperfect deconvolution may lead to more false positives for linear and periodic patterns and decrease power for the focal pattern. Fortunately, when *α*_0_ ≥ 20 (i.e., the deconvolution is not much bad), the performances of CTSV in most cell types are satisfactory.

### Model misspecification

We carried out model misspecification experiments where data were generated from a different model. Specifically, we introduced the zero-inflated Poisson log-normal regression model to generate the expression count data, *Y*_*gi*_ *∼ π*_*g*_*δ*_0_ + (1 − *π*_*g*_)Poi(*c*_*i*_*λ*_*gi*_) and 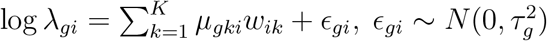. In each spatial pattern, we set the standard deviation *τ*_*g*_ = 0.1, 0.2, 0.3 and other parameters are the same as those in the original simulation study. CTSV and competing approaches were then applied. The ROC curves in Supplementary Figure S4 and TPR/FP comparison Supplementary Table S1 show that CTSV can outperform SPARK-X and remaining methods for *τ*_*g*_ = 0.1. However, when *τ*_*g*_ increases, the performance of SPARK-X begins to be better than CTSV due to a larger gap between the generating distribution and the assumed zero-inflated negative binomial distribution. Fortunately, from the perspective of detecting cell-type-specific SV genes, CTSV can still achieve relatively high accuracy for *τ*_*g*_ = 0.1, 0.2, 0.3 (Supplementary Tables S2-S4). Therefore, when ST data do not follow the zero-inflated negative binomial distribution, the nonparametric approach SPARK-X may outperform CTSV at the aggregated level. We leave the extension of CTSV to a nonparametric approach as a future work.

### Missing cell types

The notation *K* is the number of cell types across all spots in the studied tissue section. We acknowledge that it is possible that each spot *i* can have its own cell type number *K*_*i*_ (*K*_*i*_ ≤ *K*) due to the increasing resolution of spatial transcriptomics. Fortunately, our model can be easily adapted to this situation. For example, if there are *K* = 6 cell types in total and spot *i* only has three cell types (e.g., 30% cell type 1, 45% cell type 3, and 25% cell type 6), then we can let the cell-type proportion ***ω***_*i*_ for spot *i* be (0.3, 0, 0.45, 0, 0, 0.25), which is also a *K* = 6 dimensional vector. To evaluate the performance of CTSV in this “missing cell type” case, we randomly chose some spots with missing cell type number from one to five (i.e., *K*_*i*_ *∈* {1, 2, 3, 4, 5, 6}), resulting in 21 spots with only one cell type, 40 spots with two cell types, 61 spots with three cell types, 58 spots with four cell types, 52 spots with five cell types, and 368 spots with all the six cell types. For the three spatial patterns, Supplementary Figure S5 and Supplementary Table S5 show that CTSV also achieves good performances in detecting cell-type-specific SV genes.

### Mixed spatial patterns

We considered three types of mixed spatial patterns: (1) *h*_1_ is linear and *h*_2_ is periodic, where *h*_1_(*s*_*i*1_) = *s*_*i*1_ and *h*_2_(*s*_*i*2_) = cos(2*πs*_*i*2_); (2) *h*_1_ is linear and *h*_2_ is focal, where *h*_1_(*s*_*i*1_) = *s*_*i*1_ and 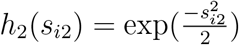 and *h*_1_ is periodic and *h*_2_ is focal, where *h*_1_(*s*_*i*1_) = cos(2*πs*_*i*1_) and 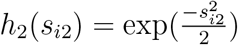. In each setting, we implemented the CTSV method described in Sections 2 and 3, where *h*_1_ and *h*_2_ used in CTSV still belong to the same pattern, so these experiments are actually also model misspecification cases. Supplementary Figure S6 and Supplementary Table S6 illustrate that under mixed spatial patterns, CTSV also outperforms other competing methods and achieves higher TPR with q-value threshold 0.01. Importantly, CTSV can still achieve relatively high accuracy in detecting cell-type-specific SV genes (Supplementary Table S7).

### Increased dropout zero proportions

To evaluate whether the FP number of CTSV increases with the dropout zero proportion *π*_*g*_, we implemented two additional settings where *π*_*g*_ = 0.7 and 0.8. The corresponding results are shown in Supplementary Figure S7 and Supplementary Tables S8-S10. Compared with other methods, we observe that CTSV is more robust to the dropout zero proportion, and the number of FP does not increase with *π*_*g*_.

### Computational speed

We set the spot size as 600, 1,000, 2,000, and 5,000 to investigate the computational time of CTSV. The average execution time per gene were 9.608 seconds, 12.187 seconds, 22.447 seconds, and 45.673 seconds, respectively, using 4 cores for paralleling. The experiments were implemented on a MacBook Pro computer with Intel Core i5, 4 cores, 8GB memory, and 2.40GHz.

## 4 Real data analysis

We applied CTSV to pancreatic ductal adenocarcinoma (PDAC) ST data (Moncada et al., 2020), which can be downloaded from Gene Expression Omnibus (Edgar et al., 2002) with accession code GSE111672, and our analysis focuses on the ST1 data from PDAC patient A. As there are associated scRNA-seq data with 18 cell types for patient A, we employed the deconvolution approach SPOTlight (Elosua-Bayes et al., 2021) to obtain cell-type proportion estimates 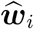 of each spot. SPOTlight is based on a seeded non-negative matrix factorization regression algorithm. It uses the ST data, scRNA-seq data, and a set of marker genes as input, and applies non-negative least squares iteratively to carry out the deconvolution.

We remark here that the deconvolution is a nontrival task and current methods do not perform equally well in different situations, so data analysts need careful considerations in choosing suitable deconvolution tools in their own problems. Here, in the pancreatic ductal adenocarcinoma (PDAC) data analysis, we chose SPOTlight (Elosua-Bayes et al., 2021) mainly for two reasons. First, SPOTlight was shown to have higher accuracy and sensitivity than other state-of-the-art deconvolution approaches based on synthetic mixture data, and it can be flexibly applied to different technical conditions and protocols. Second, the performance of SPOTlight on the PDAC data has been biologically validated, and the deconvolution results provide many insights into tumor regions (Elosua-Bayes et al., 2021).

We then merged cancer clones A and B into one cell type denoted by “cancer cell,” and combined macrophages A and B to one cell type named “macrophages.” To alleviate the effects of rare cell types, we calculated the 80th percentile of proportions across spots for each cell type and removed cell types whose 80th percentile is less than 0.1. After the procedure, six cell types—antigen presenting ductal cells, centroacinar ductal cells, high/hypoxic ductal cells, terminal ductal cells, cancer cells, and macrophages—were remained for downstream analysis, and their proportions were adjusted such that they are positive and summed to be one.

Subsequently, we filtered out genes that are expressed in less than 20 spots and kept all spots, resulting in 4,070 genes and 428 spots. The justification for using a zero-inflated distribution in CTSV in this dataset is provided in Supplementary Section S3. We afterward applied CTSV, trendsceek (Edsgärd et al., 2018), SpatialDE (Svensson et al., 2018), SPARK (Sun et al., 2020), SPARK-X (Zhu et al., 2021), SOMDE (Hao et al., 2021), and BOOST-GP (Li et al., 2021) to the processed bulk ST data. Because trendsceek and SOMDE did not detect any SV gene in PDAC dataset, we did not display them in the downstream comparisons. The Venn plot (Figure 6) shows the SV gene overlap among CTSV, SpatialDE, SPARK, SPARK-X, and BOOST-GP. When q-value threshold is 0.05, CTSV identified 61 SV genes from 4,070 genes at the aggregated level, around a half of which were also detected by SpatialDE, SPARK, SPARK-X, and BOOST-GP. In contrast, each of the competing methods detected more than 800 SV genes.

**Fig. 6:**
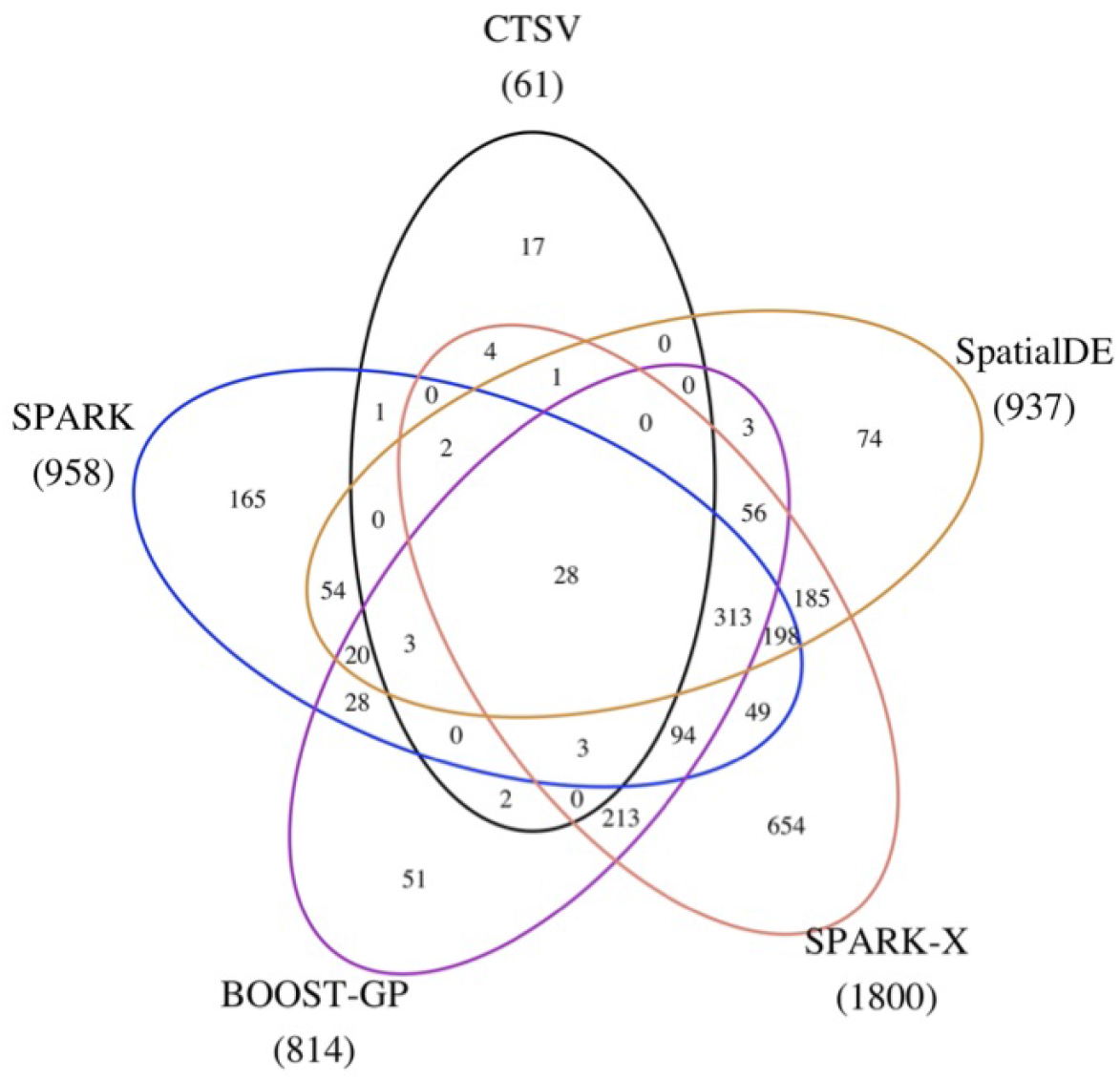
Venn plot of SV genes detected by CTSV, SPARK, BOOST-GP, SPARK-X, and SpatialDE in the PDAC data. The number in the parentheses indicates the total number of SV genes detected by that method.

For the identification of cell-type-specific SV genes, we compared the performance between CTSV and SPARK-X. In SPARK-X, each spot was assigned to the major cell type of that spot, and then SPARK-X was applied to spots that belong to the same cell type. Table 3 shows the SV gene number in each cell type for the two methods as well as the number of overlapping SV genes. We also provided the spatial expression patterns of cancer-cell-specific SV genes detected by CTSV for spots with cancer cells being the major cell type component (Figure 7). We observe that the expressions show spatial changes in the cancer regions. Specifically, genes like *CEL,CPA1* and *CLU* show relatively low expression levels in the upper right of the cancer region and have relatively high expression values in the lower middle, indicating the cancer-region-specific spatial expression variation of genes identified by CTSV. The spatial expression patterns of 673 cancer-cell-specific SV genes detected by SPARK-X are also given in Supplementary Figure S8, where more than one half of detected SV genes (e.g., *AQP8, HMGB1*, and *NDN*) show insignificant spatial variation.

**Table 3:**
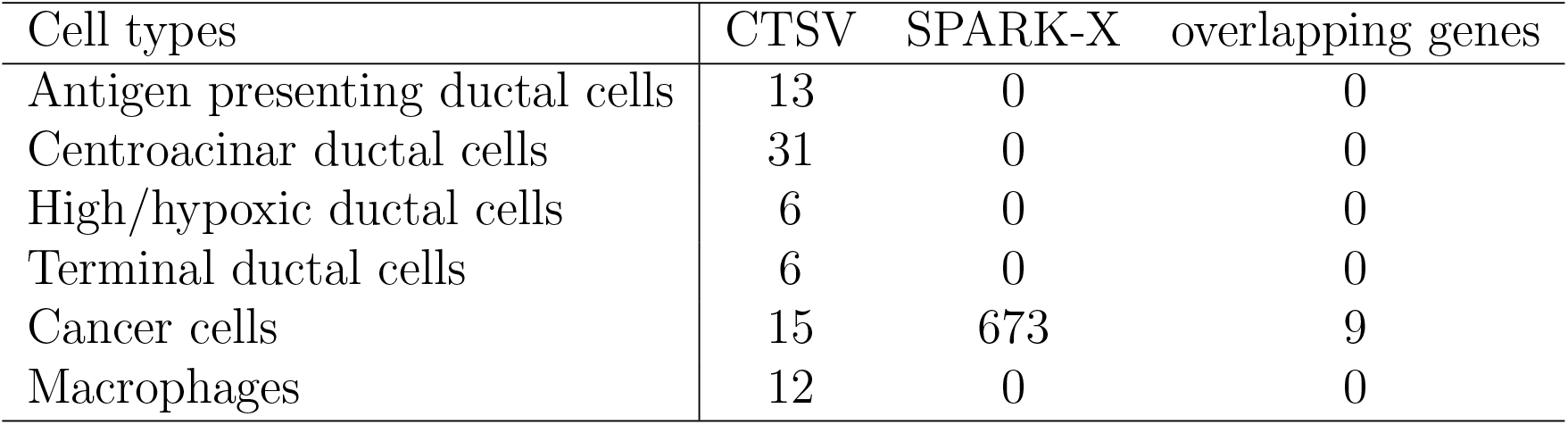
Number of SV genes in each cell type by CTSV and SPARK-X.

| Cell types | CTSV | SPARK-X | overlapping genes |
| --- | --- | --- | --- |
| Antigen presenting ductal cells | 13 | 0 | 0 |
| Centroacinar ductal cells | 31 | 0 | 0 |
| High/hypoxic ductal cells | 6 | 0 | 0 |
| Terminal ductal cells | 6 | 0 | 0 |
| Cancer cells | 15 | 673 | 9 |
| Macrophages | 12 | 0 | 0 |

**Fig. 7:**
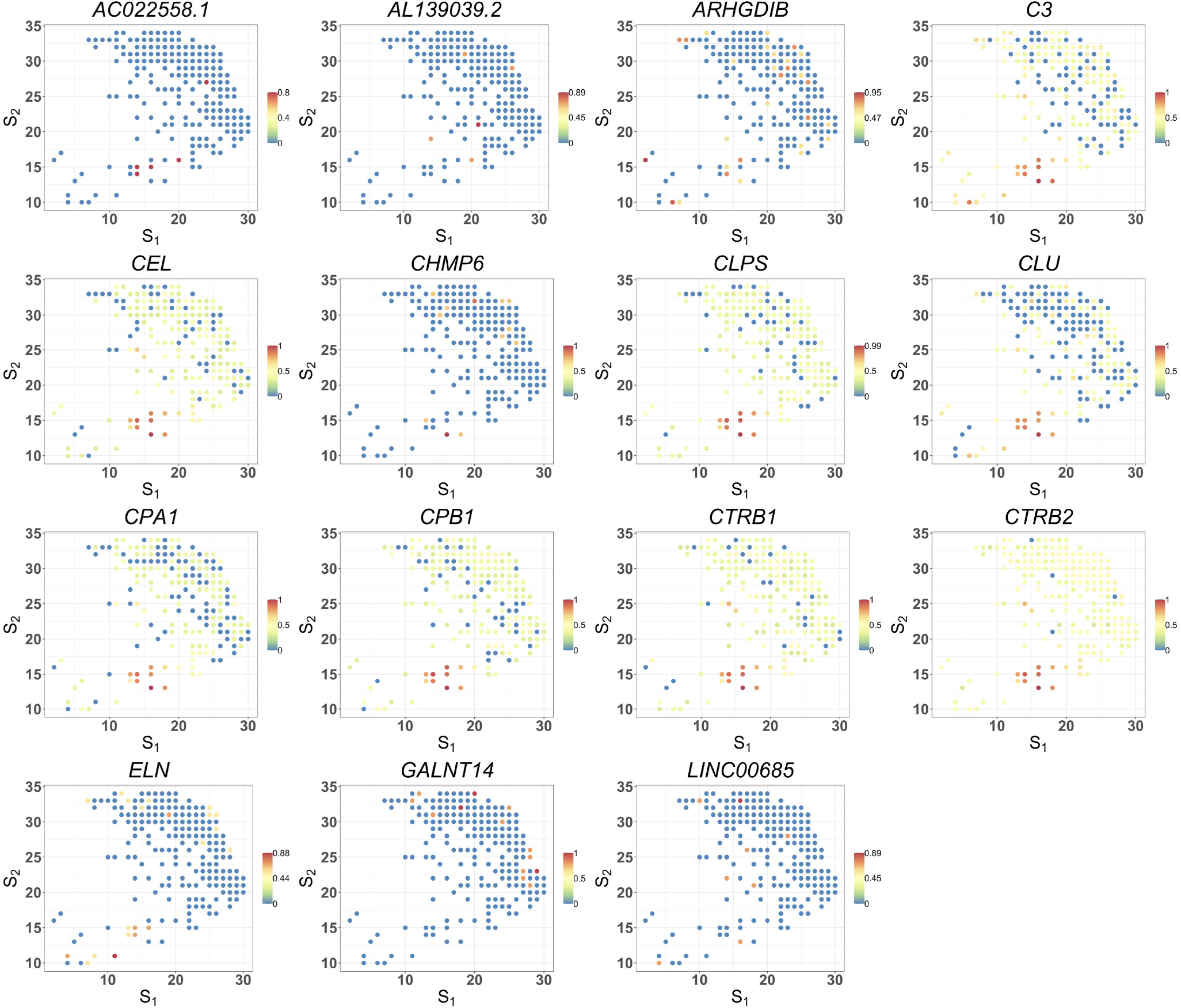
The spatial expression patterns of cancer-cell-specific SV genes detected by CTSV in the cancer region of PDAC data. Values are relative expressions, and the calculation details are given in Supplementary Section S4.

In addition, some cell-type-specific SV genes of CTSV provide some connections with tumor or PDAC. Table 4 displays these genes. For example, *ARHGDIB* in cancer cells, which was not identified by SPARK-X, encodes the protein RhoGDI2 that functions as a metastasis suppressor in human cancer (Gildea et al., 2002) and plays an important role in tumor dormancy regulation (Said et al., 2011). *ISG15* found in antigen presenting ductal cells is associated with the reinforcement of cancer stem cells’ self-renewal, invasive capacity, and tumorigenic potential in PDAC (Sainz et al., 2014). In terminal ductal cells, *JADE1* may contribute to the development of pancreatic cancer (Liu et al., 2015). *CLPS* was detected as an SV gene in more than one cell type, and the pancreatic lipase requires the colipase protein encoded by *CLPS* for efficient dietary lipid hydrolysis (Lowe, 1997; Van Tilbeurgh et al., 1999). Thus, the results by CTSV provide some clues for clarifying the underlying tumor mechanisms, which requires further validations by biological experiments.

**Table 4:**
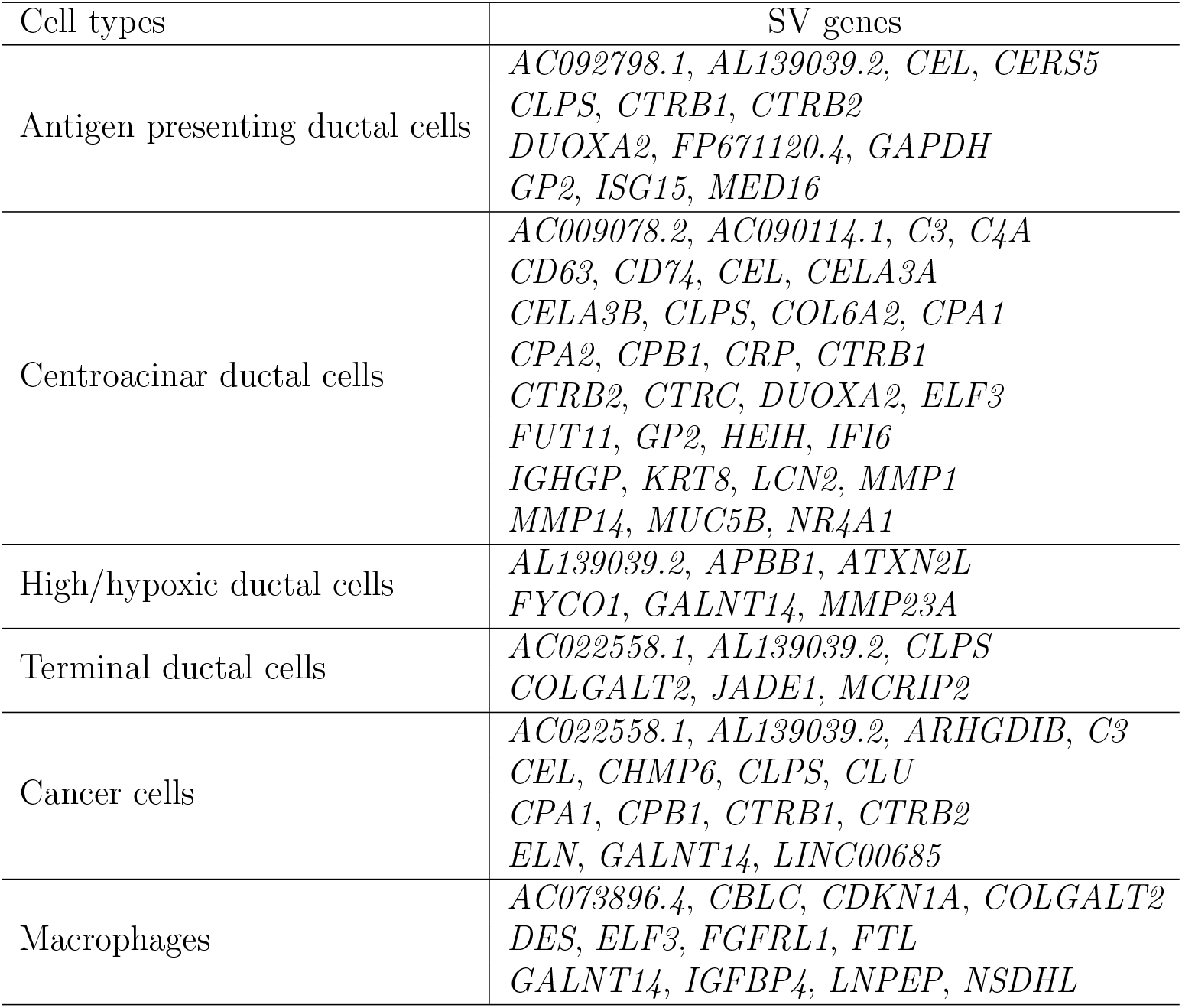
Cell-type-specific SV genes detected by CTSV

| Cell types | SV genes |
| --- | --- |
| Antigen presenting ductal cells | *AC092798.1, AL139039.2, CEL, CERS5*<br>*CLPS, CTRB1, CTRB2*<br>*DUOXA2, FP671120.4, GAPDH*<br>*GP2, ISG15, MED16* |
| Centroacinar ductal cells | *AC009078.2, AC090114.1, C3, C4A*<br>*CD63, CD74, CEL, CELA3A*<br>*CELA3B, CLPS, COL6A2, CPA1*<br>*CPA2, CPB1, CRP, CTRB1*<br>*CTRB2, CTRC, DUOXA2, ELF3*<br>*FUT11, GP2, HEIH, IFI6*<br>*IGHGP, KRT8, LCN2, MMP1*<br>*MMP14, MUC5B, NR4A1* |
| High/hypoxic ductal cells | *AL139039.2, APBB1, ATXN2L*<br>*FYCO1, GALNT14, MMP23A* |
| Terminal ductal cells | *AC022558.1, AL139039.2, CLPS*<br>*COLGALT2, JADE1, MCRIP2* |
| Cancer cells | *AC022558.1, AL139039.2, ARHGDIB, C3*<br>*CEL, CHMP6, CLPS, CLU*<br>*CPA1, CPB1, CTRB1, CTRB2*<br>*ELN, GALNT14, LINC00685* |
| Macrophages | *AC073896.4, CBLC, CDKN1A, COLGALT2*<br>*DES, ELF3, FGFRL1, FTL*<br>*GALNT14, IGFBP4, LNPEP, NSDHL* |

## 5 Conclusion

In this paper, we developed a cell-type-specific SV gene detection method (CTSV) for bulk ST data. CTSV directly models raw count data through a zero-inflated negative binomial distribution, incorporates cell-type proportions, and relies on the R package **pscl** (Zeileis et al., 2008) to fit the model. To capture different types of spatial patterns, five spatial effect functions are used, and then CTSV applied the Cauchy combination rule (Liu et al., 2019) to obtain p-values for robustness.

In simulation studies, CTSV was not only shown to be the most powerful approach at the aggregated level in the three spatial expression settings, but it also outperformed SPARK-X in terms of cell-type-specific SV gene detection, perhaps due to the direct consideration of cell-type proportions. In the analysis for pancreatic ductal adenocarcinoma data, CTSV also identified reasonable cell-type-specific SV genes that are related to meaningful biological functions.

In fact, the spatial information can be incorporated into the Gaussian process in two ways— the spatial effect on the mean vector or the spatial dependency induced by the covariance matrix. Previous methods including SpatialDE and SPARK used the covariance matrix modeling, while CTSV chose the mean to reflect spatial effects for two reasons. First, from the perspective of statistics, it is easier to test the regression coefficients *β*_*gkℓ*_ (*β*_*gkℓ*_ *∈* (−*∞*, +*∞*) with null hypothesis *H*_0_ : *β*_*gkℓ*_ = 0) in the mean function than the scale parameter *τ*_*g*1_ (*τ*_*g*1_ *∈* [0, +*∞*) with null hypothesis *H*_0_ : *τ*_*g*1_ = 0) in the covariance matrix (e.g., SPARK), as the latter is a hypothesis testing at the parameter space boundary and thus needs more complicated statistical techniques. Second, from the perspective of biology, by modeling two axes *s*_*i*1_ and *s*_*i*2_ separately, we have the opportunity to distinguish which axis may affect the gene expression based on *H*_0_ : *β*_*gk*1_ = 0 and *H*_0_ : *β*_*gk*2_ = 0. For example, it is possible that the expression changes only with *s*_*i*1_ and keeps invariant with *s*_*i*2_ (see Supplementary Figure S9). We acknowledge that CTSV can be further equipped with the spatial dependency via the covariance matrix, but this makes the statistical inference difficult and computationally inefficient. Hence, we leave it as a future work.

In addition, the performance of SPARK-X and CTSV are close in the simulation but are very different on the real data for the following reasons. On the one hand, the difference between simulation and real data may be due to the different zero-inflation rate. In pancreatic ductal adenocarcinoma ST data, the gene-wise zero proportions have the inter-quartile range [0.8014, 0.9322], while in simulation the inter-quartile range of gene-wise zero proportions is [0.5867, 0.6150] with the dropout probability *π* = 0.6. When we increase *π* from 0.6 to 0.7 and 0.8, the gap between SPARK-X and CTSV becomes larger in the ROC curves (Supplementary Figure S7). On the other hand, in the simulation setting, we let the spatial pattern be linear, focal and periodic, respectively, while in the real data analysis the spatial pattern of SV gene expressions can be more complex, e.g., the combination of two or more patterns. Therefore, these factors may make the statistical performances of CTSV and SPARK-X different between simulation and real data application.

Several extensions are worth exploring in the future. First, for robustness, we choose five simple spatial effect functions for *h*_1_ and *h*_2_, and it is better to utilize nonparametric statistical methods to directly fit the functions, such as splines or wavelets. Second, it is more helpful to incorporate prior knowledge of the tissue images (Hu et al., 2021). Third, when it comes to single-cell spatial expression data, we can also apply CTSV by setting the proportion of the cell type to which this cell belongs as one and the proportions of other cell types as zero.

Moreover, integrating multiple datasets can borrow strengths across different platforms to increase statistical power. However, due to different protocols, it may suffer from platform effects. In principle, CTSV may incorporate the platform effects *γ*_*bg*_ in platform *b* through the following modeling.

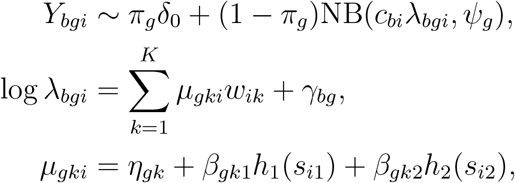

where *Y*_*bgi*_ is the read count of gene *g* for spot *i* in platform *b*, platform *b* has an additive effect *γ*_*bg*_ on the gene expression, and *γ*_1*g*_ on the platform one is fixed at zero for identifiability. Moreover, the platform may also affect the variance or the dropout zero proportion *π*_*g*_, which makes the statistical inference for CTSV more complex. Therefore, our future direction is to equip CTSV with the ability to address platform effects.

## Acknowledgements

We thank the authors of SPOTlight (Elosua-Bayes et al., 2021) for generously providing annotated single-cell RNA-seq PDAC data. This research was supported by Public Computing Cloud, Renmin University of China.

## Funding

This work was supported by the National Key R&D Program of China (Grant No. 2018YFC2000302), National Natural Science Foundation of China (Grant No. 11901572), and the fund for building world-class universities (disciplines) of Renmin University of China.

## Data availability

The PDAC datasets are publicly available in Gene Expression Omnibus with accession code GSE111672.

## Notes

### Competing Interest Statement

The authors have declared no competing interest.

