## Supplementary Materials for "Identification of Cell-Type-Specific Spatially Variable Genes Accounting for Excess Zeros"

Jinge Yu  Xiangyu Luo*

Institute of Statistics and Big Data, Renmin University of China

#### S1 Parameter choices in the simulation

Motivated by the genomic data analysis, after preprocessing, the number of genes often ranges from thousands to tens of thousands, and the spot number often ranges from hundreds (e.g., pancreatic ductal adenocarcinoma data) to thousands (e.g., mouse brain tissue data), so we set the gene number to be $G = 10,000$ and the spot number $n = 600$. We chose a mediate cell-type number $K = 6$ and simulated the mean of relative expression profile of each cell type $\boldsymbol{\eta_k}$ from normal distributions. For each cell type we chose 200 SV genes, and we set 700 SV genes at the aggregated level, which is motivated by a simulation study in Zhu et al. (2021). Furthermore, to model the biological fact that one gene may be SV in more than one cell type, we assumed there were overlaps of SV genes between different cell types. We chose three types of spatial functions $h_1$ and $h_2$ to model three possible practical expression patterns, and we set the coefficients $\beta_{gk1}$ and $\beta_{gk2}$ for SV genes to ensure that the spatial expression count data vary within a reasonable range. Moreover, we set the parameter $\psi_g$ to be 100 in the negative binomial distribution to guarantee the overdispersion of observations.

#### S2 Implementation detials for the competing methods

We provide the implementation details for the competing methods as follows.

*trendsceek*. Trendsceek was based on a point process to test whether the joint probability of expressions on two locations relies on their distance. First we converted

cell positions to point pattern using function `pos2pp`, and then normalized count expression data through `deseq_norm`, where parameters "counts" and "min.count" were set to gene expression count data $\boldsymbol{Y}$ and 1, respectively. "min.count = 1" means that we kept genes that are expressed in at least one spot. Subsequently, we used point pattern and the transformed data as input and set the parameter "log.fcn = log10" to obtain the mark distribution of the point pattern. In the end, we ran the function `trendsceek_test`, where the derived point pattern was the input and other parameters were set to default values. The R package is available on `https://github.com/edsgard/trendsceek`.

*SpatialDE*. SpatialDE measures genes' spatial effects through a zero mean Gaussian process. We first used `NaiveDE.stabilize` function to transform expression count data to continuous data, and used it as the argument "expression_matrix" parameter. Subsequently, we computed the summation of expression data of all genes in one spot and combined this information with spot locations as the argument "sample_info" of function `NavieDE.regress_out`. The "covariate_formula" was set to "np.log(total_counts)", as stated in the test example of SpatialDE on the authors' Github. The Python package is available on `https://github.com/Teichlab/SpatialDE`.

*SPARK*. SPARK was based on a Poisson log linear model and used Gaussian process to capture genes' spatial effects. When applying SPARK, we first created a SPARK object through the function `CreateSPARKObject`, the "counts" and "location" parameters were set to gene expression count data $\boldsymbol{Y}$ and spots' location matrix $\boldsymbol{S}$, respectively. The "percentage" and "min_total_counts" were both set to 0. We then used the SPARK object for fitting the count-based spatial model to estimate the parameters through the R fcuntion `spark.vc`. Lastly, we let the fitted object be the "object" input of the function `spark.test`. Other parameters were set to be the same as default values. The R package is available on `https://github.com/xzhoulab/SPARK`.

*SPARK-X*. SPARK-X is a nonparametric method built for identifying SV genes. We used the function `sparkx` in R package SPARK to perform SV gene detection both at the aggregated level and at the cell-type-specific level. When implementing SPARK-X at the aggregated level, we set the "count_in" parameter to gene expression count data $\boldsymbol{Y}$, and the "locus_in" parameter to location matrix $\boldsymbol{S}$. When conducting cell-type-specific SV gene detection, we first assigned each spot to its major cell type and then performed SPARK-X on those spots dominated by the same cell type. For each cell type, the input of R function `sparkx` were $\boldsymbol{Y}_{sub}$ and $\boldsymbol{S}_{sub}$—the gene expression data of spots belonging to one cell type and their spot locations. Under each scenario, we chose the parameter "numCores = 10", and let other parameters be the same as default values. The R package is available on `https://github.com/xzhoulab/SPARK`.

*BOOST-GP*. BOOST-GP was based on a zero-inflated negative binomial model and captured spatial effects of genes via a zero mean Gaussian process. It performed inference in the Bayesian framework, and we applied BOOST-GP through function boost.gp. We set the iteration number 100, "burn = 50" in boost.gp, and "cutoff" parameter 0.2 in function distance for fast computation. The codes are available on https://github.com/Minzhe/BOOST-GP.

*SOMDE.* SOMDE used self-organizing map to cluster neighboring cells, and then utilized a Gaussian process for SV gene detection. First, we built SOM through the function SomNode with spot locations as input and set the parameter "k = 20" as well as the function mtx. Subsequently, we normalized the count data using norm and then identified SV gene using run. For other implementation steps, we followed the tutorial on the author's Github. The Python package is available on https://github.com/XuegongLab/somde.

### S3 Justification for using a zero-inflated distribution

In the negative binomial (NB) distribution, for gene $g$, if its expression count $Y_{gi}$ follows $\mathrm{NB}(\theta_g, \psi_g)$ with mean $\theta_g$ and dispersion $\psi_g$, then the theoretical probability of $Y_{gi}$ being zero equals $\Pr(Y_{gi} = 0) = \psi_g^{\psi_g}/(\theta_g + \psi_g)^{\psi_g}$. If the observed zero proportion is larger than the estimated $\Pr(Y_{gi} = 0)$ under the NB model in real data, then we believe the ST data for gene $g$ is zero-inflated.

Subsequently, in real PDAC data, we use the NB distribution to fit the ST count data for each gene. The theoretical zero probability under NB $\psi_g^{\psi_g}/(\theta_g + \psi_g)^{\psi_g}$ is estimated as follows. First, we calculate the sample mean and the sample variance of the expression data, denoted by $\overline{\mathbf{Y}}_g$ and $\mathrm{var}(\mathbf{Y}_g)$. Next, we employ the method of moments estimator for $\theta_g$ and $\psi_g$ by equating $\widehat{\theta}_g = \overline{\mathbf{Y}}_g$, $\widehat{\psi}_g = \frac{\hat{\theta}_g^2}{\mathrm{var}(\mathbf{Y}_g) - \hat{\theta}_g}$. Finally, we use $\widehat{\psi}_g^{\widehat{\psi}_g}/(\widehat{\theta}_g + \widehat{\psi}_g)^{\widehat{\psi}_g}$ as the estimated probability of $Y_{gi}$ being zero. Among the 4,049 genes after data preprocessing, there are 1,835 genes with the observed zero proportion larger than the estimated zero probability under NB. Thus, for a large amount of genes, the NB distribution is not satisfactory to capture extra dropout zeros in the real PDAC data.

On the other hand, the NB distribution is a special case of the zero-inflated NB distribution by letting $\pi_g = 0$. Therefore, fitting ST data by zero-inflated NB can also estimate associated parameters very well. Moreover, the zero-inflated NB distribution has been employed in previous works such as BOOST-GP (Li et al., 2021) for modeling ST data. Therefore, at least for the current PDAC data, it is necessary to model the ST count data via the zero-inflated NB distribution.

### S4 Values in Figure 7

Values in Figure 7 are the relative expression values of genes, which range from zero to one. To obtain the relative expression values, we first transform the gene expression count data $\boldsymbol{Y}$ to continuous values $\boldsymbol{Y}^*$, and then for each gene $g$ we normalize $\boldsymbol{Y}_g^* = (Y_{g1}^*, \ldots, Y_{gn}^*)$ to $\boldsymbol{Y}_g^{**}$ to make sure $0 \leq Y_{gi}^{**} \leq 1$, $i = 1, 2, \ldots n$, which follows Sun et al. (2020). In the first step, we calculate the mean and variance of expression denoted by $\overline{Y_g}$ and $\mathrm{var}(\boldsymbol{Y}_g)$ for each gene, and apply nonlinear least square regression through R function `nls` to obtain the regression coefficient $\rho$ of the regression formula $\mathrm{var}(\boldsymbol{Y}_g) \sim \overline{Y_g} + \rho\overline{Y_g}^2$. We obtain continuous values $Y_{gi}^* = \log(Y_{gi} + \frac{1}{2\rho})$. In the second step, we have $Y_{gi}^{**} = \frac{Y_{gi}^* - \min_{1\leq i\leq n}\{Y_{gi}^*\}}{\max_{1\leq i\leq n}\{Y_{gi}^*\} - \min_{1\leq i\leq n}\{Y_{gi}^*\}}$.

Table S1: The TPR and FP comparisons in the SV gene detection at the aggregated level under model misspecification.

| | Pattern | CTSV | SPARK-X | SPARK | BOOST-GP | SpatialDE | SOMDE | trendsceek |
|---|---|---|---|---|---|---|---|---|
| $\tau_g = 0.1$ | | | | | | | | |
| TPR | Linear | 0.991 | 0.853 | 0.054 | 0 | 0 | 0 | 0 |
| | Focal | 0.613 | 0.124 | 0 | 0.001 | 0 | 0 | 0 |
| | Periodic | 0.999 | 0.817 | 0 | 0.001 | 0 | 0 | 0 |
| FP | Linear | 25 | 3 | 0 | 9 | 0 | 0 | 0 |
| | Focal | 9 | 2 | 0 | 2 | 0 | 0 | 0 |
| | Periodic | 28 | 4 | 0 | 4 | 0 | 0 | 0 |
| $\tau_g = 0.2$ | | | | | | | | |
| TPR | Linear | 0.950 | 0.844 | 0.071 | 0 | 0 | 0 | 0 |
| | Focal | 0.351 | 0.076 | 0 | 0.001 | 0 | 0 | 0 |
| | Periodic | 0.996 | 0.817 | 0 | 0.001 | 0 | 0 | 0 |
| FP | Linear | 32 | 2 | 0 | 9 | 0 | 0 | 0 |
| | Focal | 11 | 1 | 0 | 2 | 0 | 0 | 0 |
| | Periodic | 35 | 5 | 0 | 4 | 0 | 0 | 0 |
| $\tau_g = 0.3$ | | | | | | | | |
| TPR | Linear | 0.883 | 0.820 | 0.050 | 0 | 0 | 0 | 0 |
| | Focal | 0.176 | 0.066 | 0 | 0.001 | 0 | 0 | 0 |
| | Periodic | 0.976 | 0.800 | 0 | 0.001 | 0 | 0 | 0 |
| FP | Linear | 74 | 2 | 0 | 9 | 0 | 0 | 0 |
| | Focal | 35 | 1 | 0 | 2 | 0 | 0 | 0 |
| | Periodic | 106 | 1 | 0 | 4 | 0 | 0 | 0 |

Table S2: Cell-type-specific TPR and FP of CTSV and SPARK-X under model misspecification with $\tau_g = 0.1$.

| Spatial pattern | | Linear | | Focal | | Periodic | |
|---|---|---|---|---|---|---|---|
| Methods | | CTSV | SPARK-X | CTSV | SPARK-X | CTSV | SPARK-X |
| TPR | cell-type 1 | 0.960 | 0.300 | 0.535 | 0 | 1.000 | 0 |
| | cell-type 2 | 0.975 | 0.060 | 0.390 | 0 | 0.955 | 0 |
| | cell-type 3 | 0.935 | 0 | 0.255 | 0 | 0.965 | 0 |
| | cell-type 4 | 0.950 | 0 | 0.220 | 0 | 0.970 | 0 |
| | cell-type 5 | 0.965 | 0 | 0.440 | 0 | 0.965 | 0 |
| | cell-type 6 | 0.985 | 0 | 0.750 | 0 | 1.000 | 0 |
| FP | cell-type 1 | 33 | 20 | 3 | 0 | 4 | 0 |
| | cell-type 2 | 12 | 0 | 0 | 0 | 3 | 0 |
| | cell-type 3 | 4 | 0 | 4 | 0 | 3 | 0 |
| | cell-type 4 | 7 | 0 | 6 | 0 | 4 | 0 |
| | cell-type 5 | 5 | 0 | 2 | 0 | 5 | 0 |
| | cell-type 6 | 11 | 0 | 54 | 0 | 10 | 0 |

Table S3: Cell-type-specific TPR and FP of CTSV and SPARK-X under model misspecification with $\tau_g = 0.2$.

| Spatial pattern | | Linear | | Focal | | Periodic | |
|---|---|---|---|---|---|---|---|
| Methods | | CTSV | SPARK-X | CTSV | SPARK-X | CTSV | SPARK-X |
| TPR | cell-type 1 | 0.940 | 0.260 | 0.235 | 0 | 0.980 | 0 |
| | cell-type 2 | 0.920 | 0.055 | 0.170 | 0 | 0.860 | 0 |
| | cell-type 3 | 0.790 | 0 | 0.125 | 0 | 0.925 | 0 |
| | cell-type 4 | 0.845 | 0 | 0.085 | 0 | 0.950 | 0 |
| | cell-type 5 | 0.915 | 0 | 0.240 | 0 | 0.925 | 0 |
| | cell-type 6 | 0.890 | 0 | 0.465 | 0 | 1.000 | 0 |
| FP | cell-type 1 | 22 | 12 | 4 | 0 | 5 | 0 |
| | cell-type 2 | 17 | 0 | 2 | 0 | 6 | 0 |
| | cell-type 3 | 5 | 0 | 0 | 0 | 6 | 0 |
| | cell-type 4 | 5 | 0 | 7 | 0 | 7 | 0 |
| | cell-type 5 | 8 | 0 | 2 | 0 | 6 | 0 |
| | cell-type 6 | 12 | 0 | 30 | 0 | 8 | 0 |

Table S4: Cell-type-specific TPR and FP of CTSV and SPARK-X under model misspecification with $\tau_g = 0.3$.

| Spatial pattern | | Linear | | Focal | | Periodic | |
|---|---|---|---|---|---|---|---|
| Methods | | CTSV | SPARK-X | CTSV | SPARK-X | CTSV | SPARK-X |
| TPR | cell-type 1 | 0.815 | 0.255 | 0.075 | 0 | 0.965 | 0 |
| | cell-type 2 | 0.750 | 0.035 | 0.070 | 0 | 0.740 | 0 |
| | cell-type 3 | 0.600 | 0 | 0.045 | 0 | 0.840 | 0 |
| | cell-type 4 | 0.660 | 0 | 0.055 | 0 | 0.810 | 0 |
| | cell-type 5 | 0.715 | 0 | 0.055 | 0 | 0.880 | 0 |
| | cell-type 6 | 0.800 | 0 | 0.300 | 0 | 0.980 | 0 |
| FP | cell-type 1 | 20 | 12 | 6 | 0 | 23 | 0 |
| | cell-type 2 | 19 | 0 | 5 | 0 | 15 | 0 |
| | cell-type 3 | 24 | 0 | 3 | 0 | 23 | 0 |
| | cell-type 4 | 13 | 0 | 9 | 0 | 17 | 0 |
| | cell-type 5 | 11 | 0 | 12 | 0 | 20 | 0 |
| | cell-type 6 | 20 | 0 | 12 | 0 | 21 | 0 |

Table S5: Cell-type-specific TPR and FP of CTSV in the missing cell type cases.

| | Pattern | cell-type 1 | cell-type 2 | cell-type 3 | cell-type 4 | cell-type 5 | cell-type 6 |
|---|---|---|---|---|---|---|---|
| TPR | Linear | 0.855 | 0.975 | 0.970 | 0.980 | 0.990 | 0.890 |
| | Focal | 0.525 | 0.265 | 0.225 | 0.290 | 0.540 | 0.230 |
| | Periodic | 1.000 | 1.000 | 1.000 | 1.000 | 1.000 | 1.000 |
| FP | Linear | 12 | 8 | 9 | 3 | 2 | 2 |
| | Focal | 0 | 2 | 1 | 1 | 1 | 3 |
| | Periodic | 3 | 5 | 4 | 3 | 5 | 3 |

Table S6: The comparisons of TPR and the number of FP in SV gene detection at the aggregated level in the mixed spatial pattern cases. "Lin-Per" means that $h_1$ is linear and $h_2$ is periodic; "Lin-Foc" means that $h_1$ is linear and $h_2$ is focal; "Per-Foc" means that $h_1$ is periodic and $h_2$ is focal.

| | Pattern | CTSV | SPARK-X | SPARK | BOOST-GP | SpatialDE | SOMDE | trendsceek |
|---|---|---|---|---|---|---|---|---|
| TPR | Lin-Per | 0.971 | 0.814 | 0.134 | 0.003 | 0 | 0 | 0 |
| | Lin-Foc | 0.999 | 0.850 | 0.060 | 0.010 | 0 | 0 | 0 |
| | Per-Foc | 1.000 | 0.816 | 0 | 0 | 0 | 0 | 0 |
| FP | Lin-Per | 20 | 3 | 0 | 9 | 0 | 0 | 0 |
| | Lin-Foc | 38 | 3 | 0 | 6 | 0 | 0 | 0 |
| | Per-Foc | 24 | 3 | 0 | 4 | 0 | 0 | 0 |

Table S7: Cell-type-specific TPR and FP of CTSV and SPARK-X in the mixed spatial pattern cases.

| Spatial pattern | | Lin-Per | | Lin-Foc | | Per-Foc | |
|---|---|---|---|---|---|---|---|
| Methods | | CTSV | SPARK-X | CTSV | SPARK-X | CTSV | SPARK-X |
| TPR | cell-type 1 | 0.885 | 0.160 | 0.995 | 0.260 | 1.000 | 0 |
| | cell-type 2 | 0.940 | 0.005 | 0.995 | 0.050 | 0.970 | 0 |
| | cell-type 3 | 0.785 | 0 | 0.960 | 0 | 0.990 | 0 |
| | cell-type 4 | 0.805 | 0 | 0.980 | 0 | 0.995 | 0 |
| | cell-type 5 | 0.940 | 0 | 0.990 | 0 | 0.980 | 0 |
| | cell-type 6 | 0.875 | 0 | 1.000 | 0 | 1.000 | 0 |
| FP | cell-type 1 | 21 | 5 | 28 | 9 | 1 | 0 |
| | cell-type 2 | 17 | 0 | 19 | 0 | 6 | 0 |
| | cell-type 3 | 2 | 0 | 3 | 0 | 6 | 0 |
| | cell-type 4 | 7 | 0 | 10 | 0 | 4 | 0 |
| | cell-type 5 | 2 | 0 | 5 | 0 | 5 | 0 |
| | cell-type 6 | 15 | 0 | 53 | 0 | 2 | 0 |

Table S8: The comparisons of TPR and FP in SV gene detection at the aggregated level when $\pi_g$ is 0.7 and 0.8.

| | | CTSV | SPARK-X | SPARK | BOOST-GP | SpatialDE | SOMDE | trendsceek |
|---|---|---|---|---|---|---|---|---|
| $\pi_g = 0.7$ | | | | | | | | |
| TPR | Linear | 0.980 | 0.781 | 0.010 | 0 | 0 | 0 | 0 |
| | Focal | 0.703 | 0.034 | 0 | 0 | 0 | 0 | 0 |
| | Periodic | 0.976 | 0.736 | 0 | 0 | 0 | 0 | 0 |
| FP | Linear | 38 | 0 | 0 | 7 | 0 | 0 | 0 |
| | Focal | 13 | 0 | 0 | 7 | 0 | 0 | 0 |
| | Periodic | 19 | 2 | 0 | 8 | 0 | 0 | 0 |
| $\pi_g = 0.8$ | | | | | | | | |
| TPR | Linear | 0.890 | 0.446 | 0.001 | 0 | 0 | 0 | 0 |
| | Focal | 0.487 | 0.004 | 0 | 0 | 0 | 0 | 0 |
| | Periodic | 0.814 | 0.473 | 0 | 0 | 0 | 0 | 0 |
| FP | Linear | 24 | 0 | 0 | 4 | 0 | 0 | 0 |
| | Focal | 9 | 0 | 0 | 5 | 0 | 0 | 0 |
| | Periodic | 12 | 1 | 0 | 6 | 0 | 0 | 0 |

Table S9: Cell-type-specific TPR and FP of CTSV and SPARK-X with $\pi_g = 0.7$.

| Spatial pattern | | Linear | | Focal | | Periodic | |
|---|---|---|---|---|---|---|---|
| Methods | | CTSV | SPARK-X | CTSV | SPARK-X | CTSV | SPARK-X |
| TPR | cell-type 1 | 0.985 | 0.120 | 0.495 | 0 | 0.975 | 0 |
| | cell-type 2 | 0.945 | 0 | 0.525 | 0 | 0.770 | 0 |
| | cell-type 3 | 0.925 | 0 | 0.400 | 0 | 0.880 | 0 |
| | cell-type 4 | 0.915 | 0 | 0.310 | 0 | 0.890 | 0 |
| | cell-type 5 | 0.945 | 0 | 0.495 | 0 | 0.875 | 0 |
| | cell-type 6 | 0.970 | 0 | 0.755 | 0 | 0.980 | 0 |
| FP | cell-type 1 | 27 | 12 | 7 | 0 | 4 | 0 |
| | cell-type 2 | 11 | 0 | 6 | 0 | 3 | 0 |
| | cell-type 3 | 10 | 0 | 3 | 0 | 4 | 0 |
| | cell-type 4 | 9 | 0 | 4 | 0 | 4 | 0 |
| | cell-type 5 | 7 | 0 | 6 | 0 | 1 | 0 |
| | cell-type 6 | 17 | 0 | 69 | 0 | 3 | 0 |

Table S10: Cell-type-specific TPR and FP of CTSV and SPARK-X with $\pi_g = 0.8$.

| Spatial pattern | | Linear | | Focal | | Periodic | |
|---|---|---|---|---|---|---|---|
| Methods | | CTSV | SPARK-X | CTSV | SPARK-X | CTSV | SPARK-X |
| TPR | cell-type 1 | 0.795 | 0.010 | 0.200 | 0 | 0.820 | 0 |
| | cell-type 2 | 0.730 | 0 | 0.305 | 0 | 0.450 | 0 |
| | cell-type 3 | 0.635 | 0 | 0.245 | 0 | 0.525 | 0 |
| | cell-type 4 | 0.645 | 0 | 0.225 | 0 | 0.510 | 0 |
| | cell-type 5 | 0.775 | 0 | 0.290 | 0 | 0.550 | 0 |
| | cell-type 6 | 0.845 | 0 | 0.475 | 0 | 0.890 | 0 |
| FP | cell-type 1 | 20 | 0 | 11 | 0 | 5 | 0 |
| | cell-type 2 | 11 | 0 | 16 | 0 | 1 | 0 |
| | cell-type 3 | 4 | 0 | 9 | 0 | 1 | 0 |
| | cell-type 4 | 11 | 0 | 8 | 0 | 0 | 0 |
| | cell-type 5 | 1 | 0 | 8 | 0 | 4 | 0 |
| | cell-type 6 | 13 | 0 | 49 | 0 | 4 | 0 |

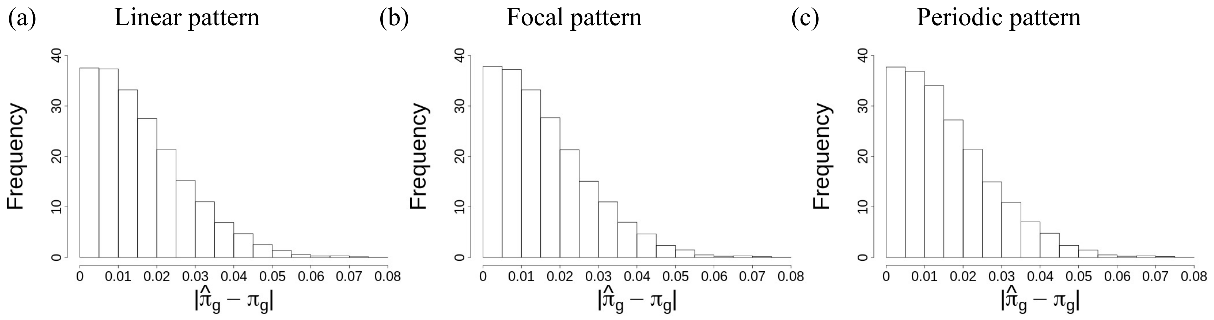


Figure S1: Histograms for absolute estimation errors $|\hat{\pi}_g - \pi_g|$ in different spatial expression patterns.

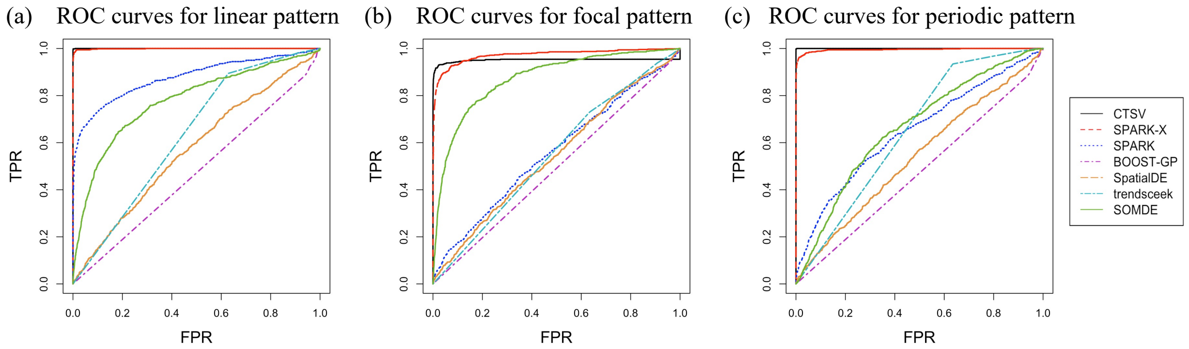


Figure S2: ROC curves over the whole FPR range $(0, 1)$ for CTSV, SPARK-X, SPARK, BOOST-GP, SpatialDE, trendsceek, and SOMDE in the three spatial expression patterns.

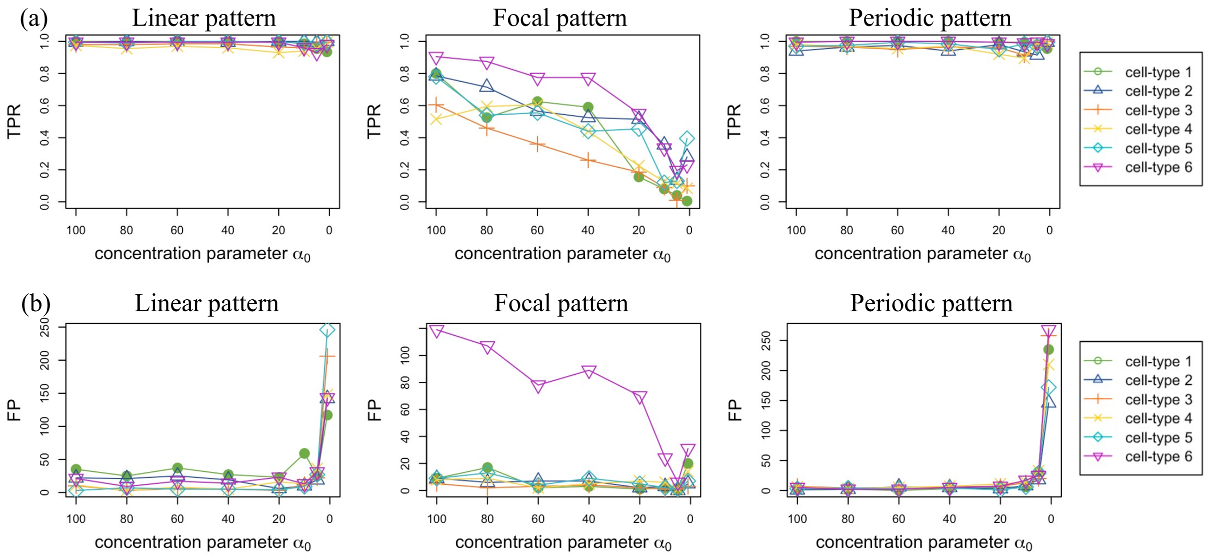


Figure S3: CTSV performance with imperfect deconvolution. Different colors and line types correspond to different cell types. (a) The TPR plot with different values of $\alpha_0$ in the linear, focal, and periodic pattern. (b) The FP plot with different values of $\alpha_0$ in the linear, focal, and periodic pattern.

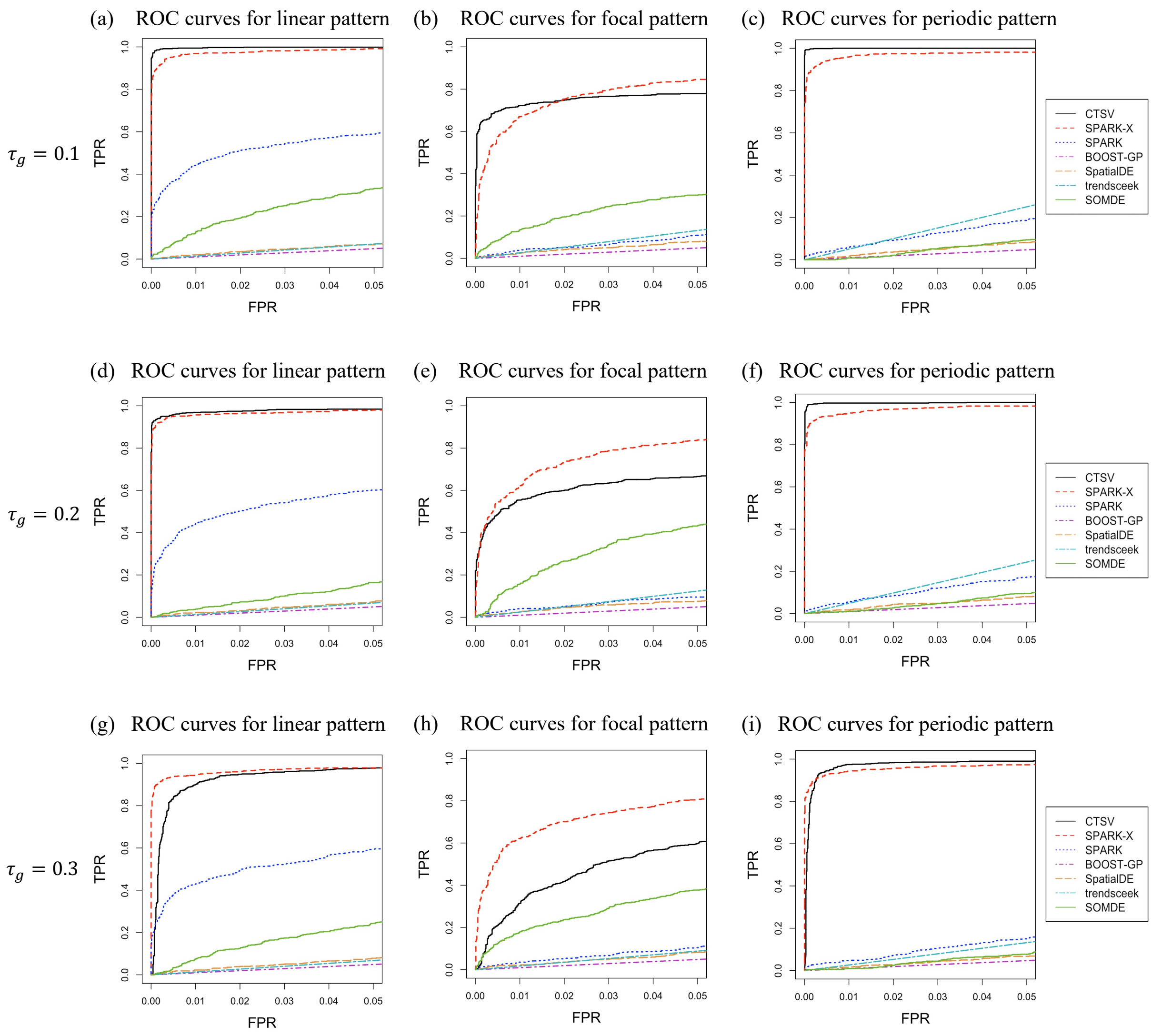


Figure S4: ROC curves with FPR less than 0.05 for three spatial patterns in the model misspecification cases with (a-c) $\tau_g = 0.1$, (d-f) $\tau_g = 0.2$, and (g-i) $\tau_g = 0.3$.

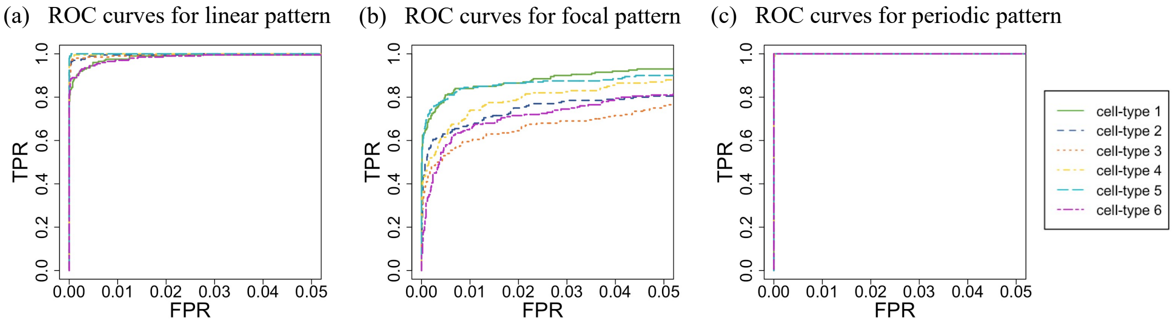


Figure S5: Cell-type-specific ROC curves with FPR less than 0.05 in the missing cell type cases.

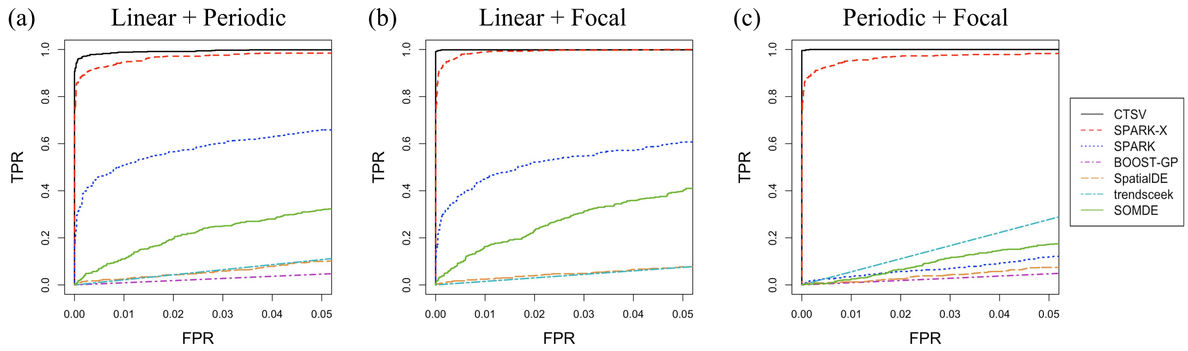


Figure S6: ROC curves with FPR less than 0.05 in the mixed spatial pattern cases. (a) $h_1$ is linear and $h_2$ is periodic. (b) $h_1$ is linear and $h_2$ is focal. (c) $h_1$ is periodic and $h_2$ is focal.

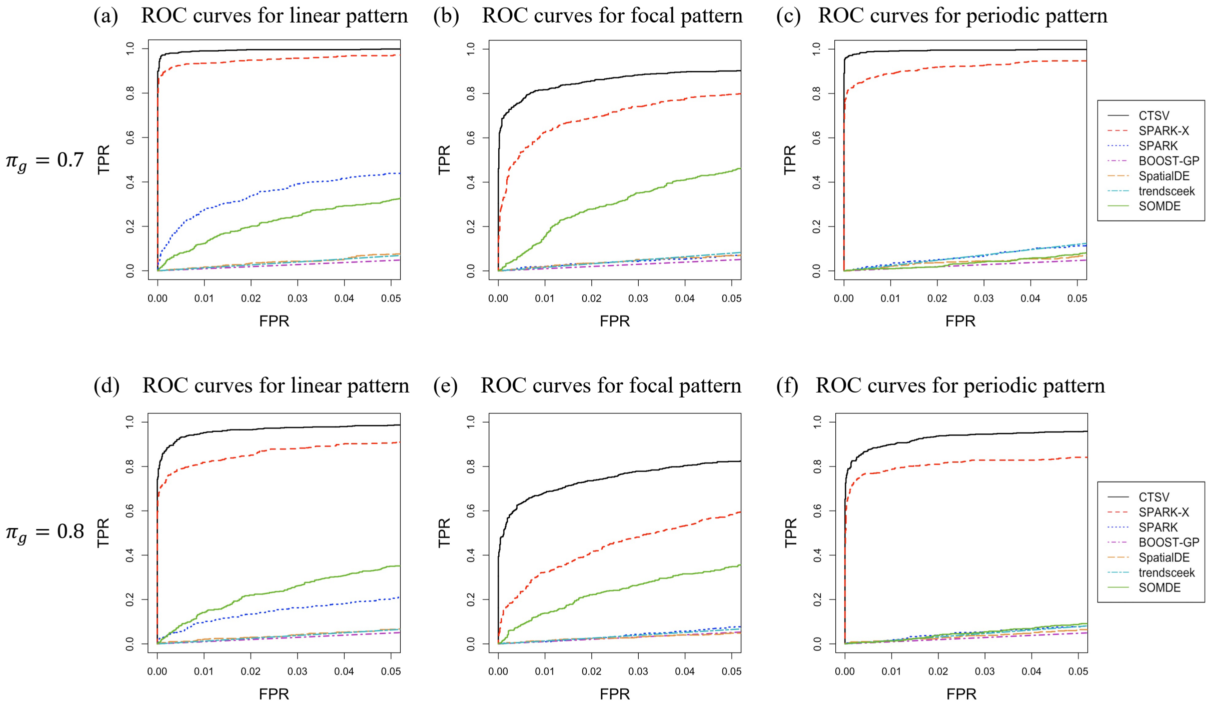


Figure S7: ROC curves with FPR less than 0.05 for CTSV, SPARK-X, SPARK, BOOST-GP, SpatialDE, trendsceek, and SOMDE with $\pi_g = 0.7, 0.8$. (a-c) The ROC curves in the three spatial expression patterns with $\pi_g = 0.7$. (d-f) The ROC curves in the three spatial expression patterns with $\pi_g = 0.8$.

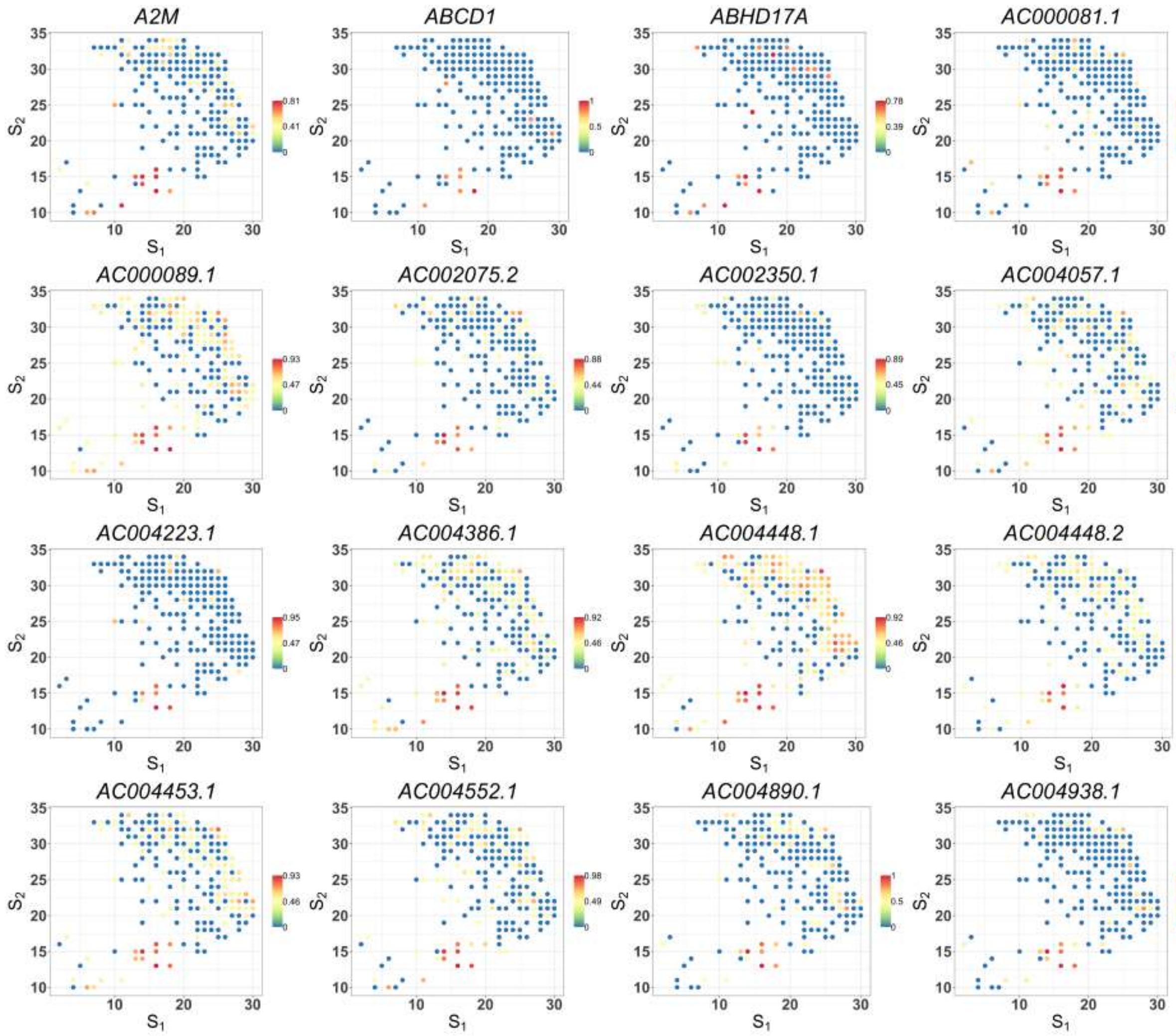


Figure S8: Spatial expression patterns of cancer-cell-specific SV genes detected by SPARK-X in the cancer region of PDAC data (part 1). Values are relative expressions.

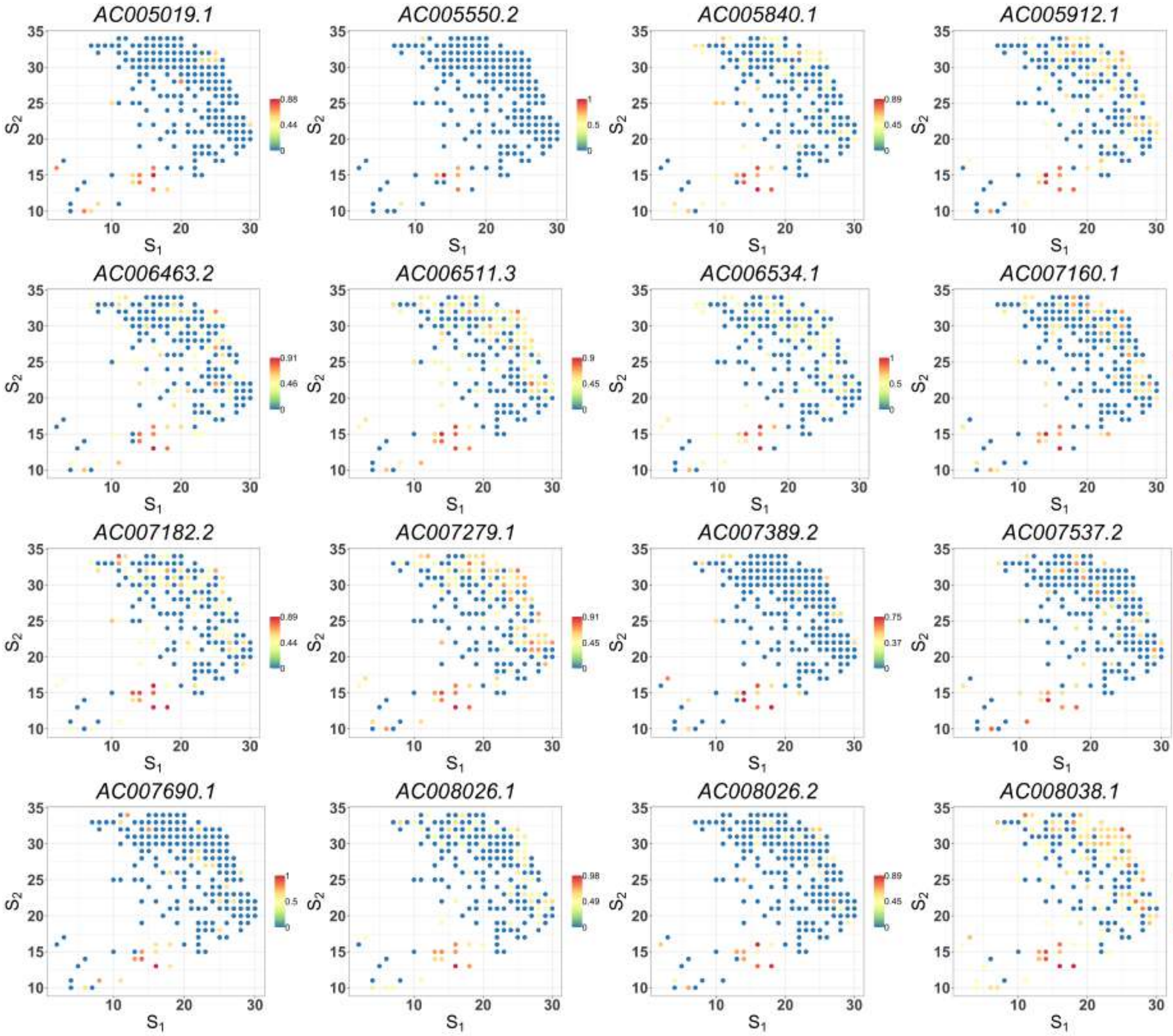


Figure S8: Spatial expression patterns of cancer-cell-specific SV genes detected by SPARK-X in the cancer region of PDAC data (part 2). Values are relative expressions.

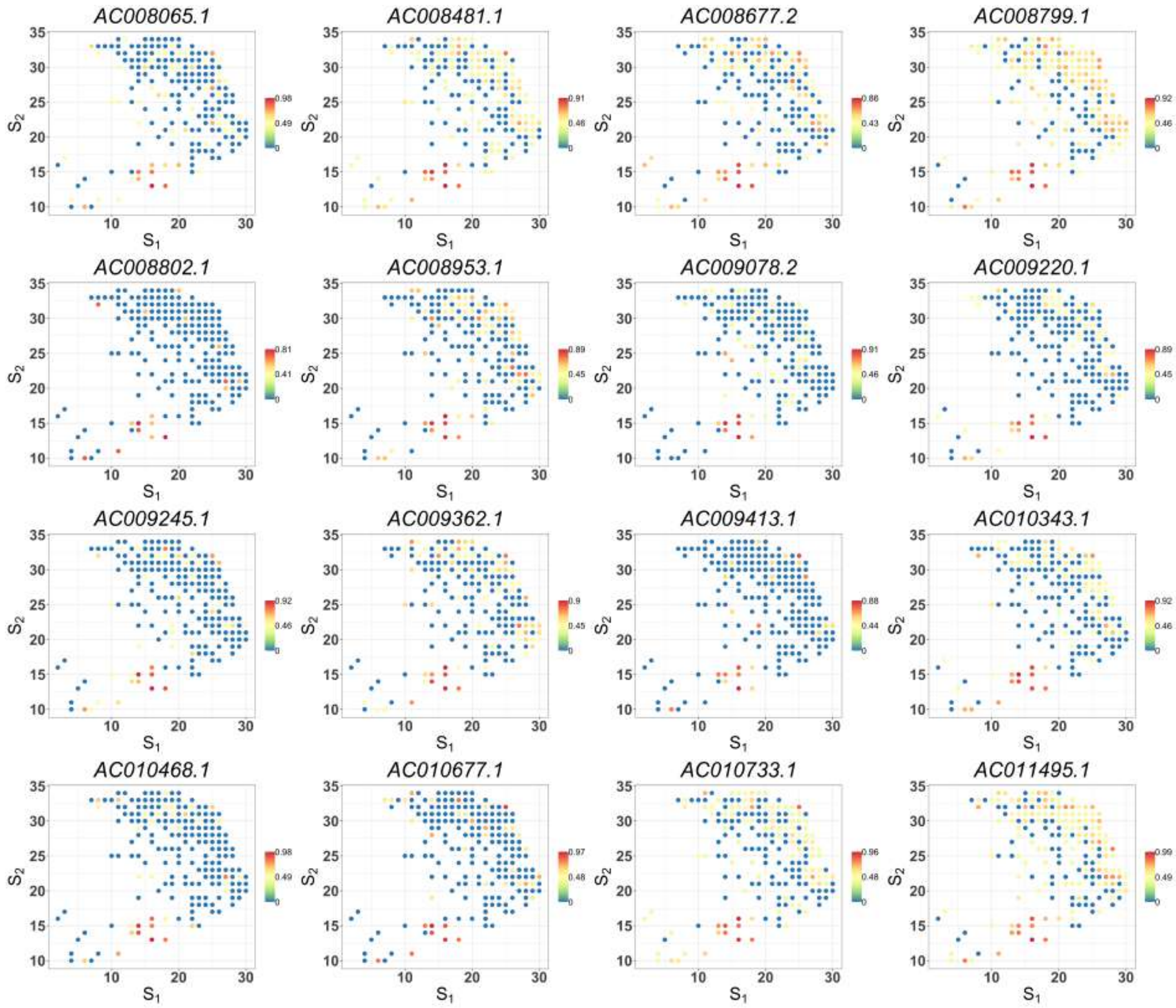


Figure S8: Spatial expression patterns of cancer-cell-specific SV genes detected by SPARK-X in the cancer region of PDAC data (part 3). Values are relative expressions.

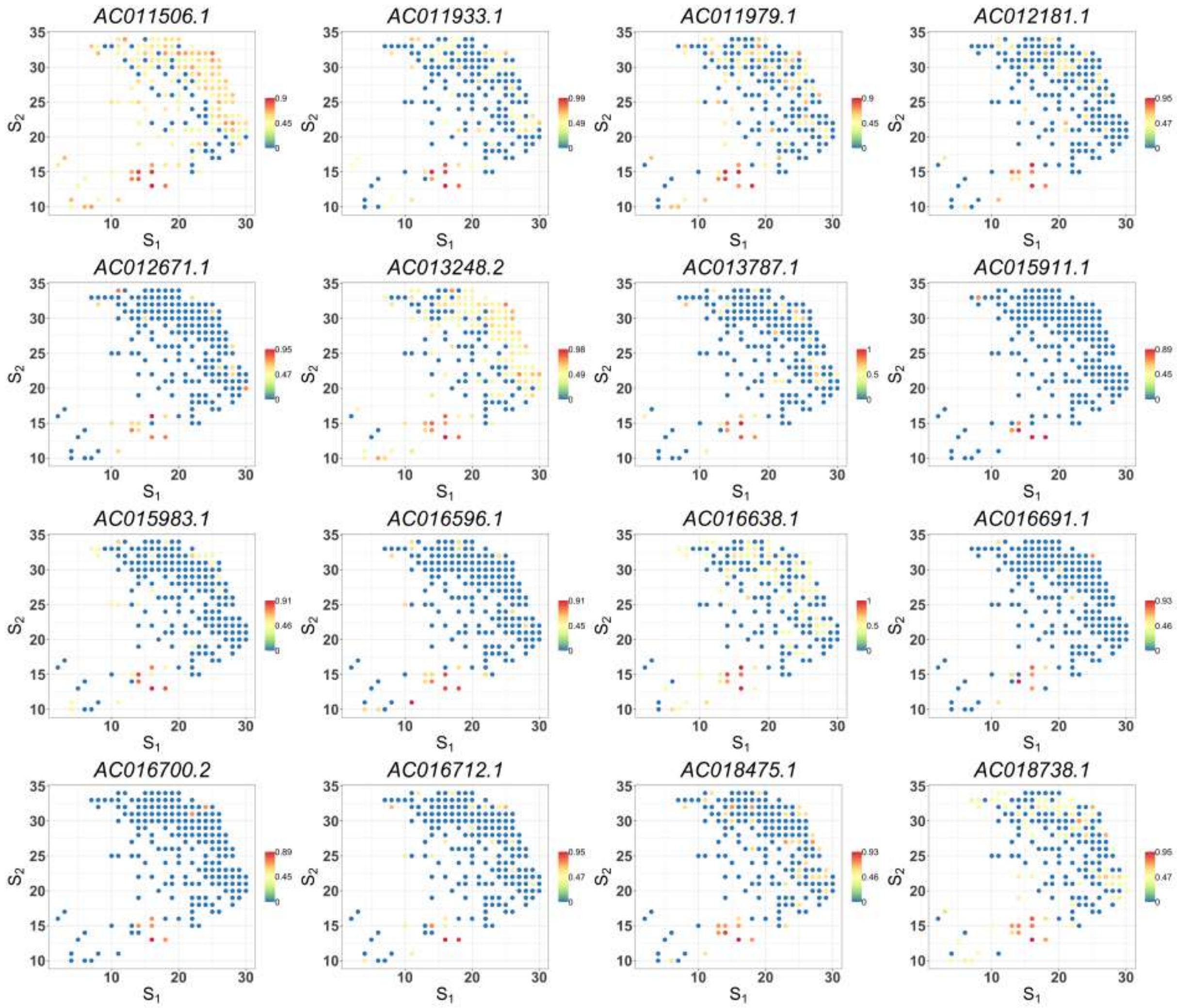


Figure S8: Spatial expression patterns of cancer-cell-specific SV genes detected by SPARK-X in the cancer region of PDAC data (part 4). Values are relative expressions.

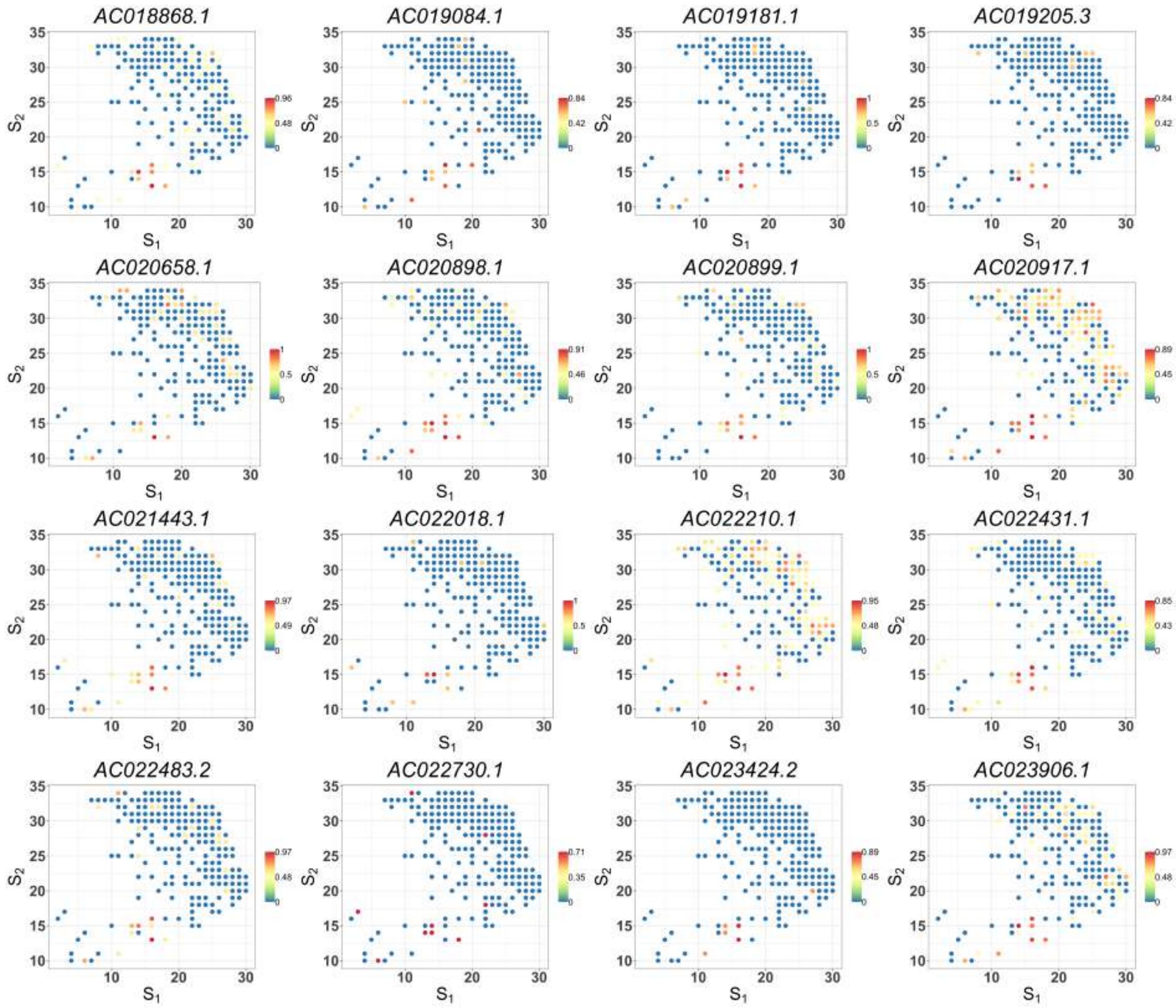


Figure S8: Spatial expression patterns of cancer-cell-specific SV genes detected by SPARK-X in the cancer region of PDAC data (part 5). Values are relative expressions.

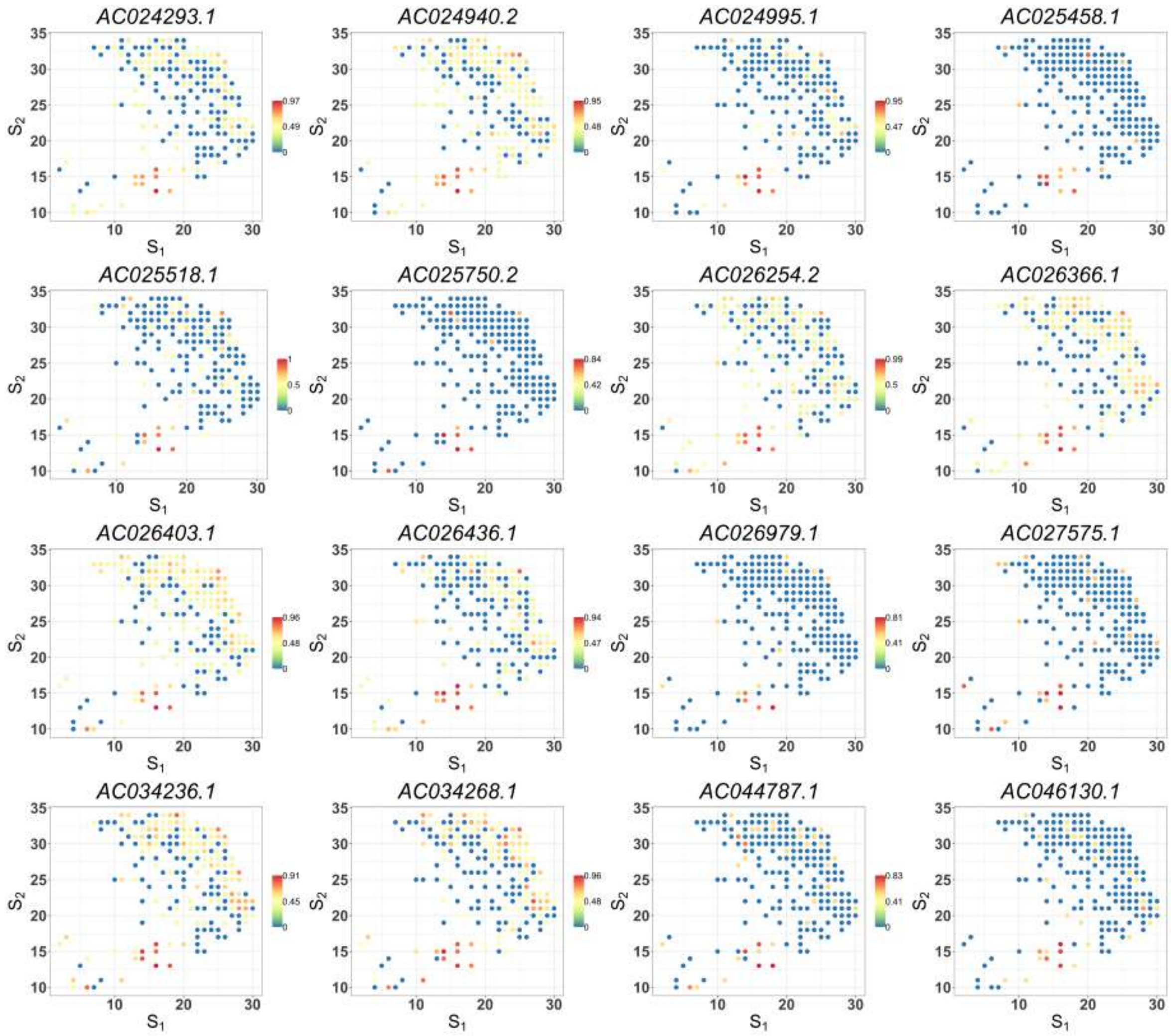


Figure S8: Spatial expression patterns of cancer-cell-specific SV genes detected by SPARK-X in the cancer region of PDAC data (part 6). Values are relative expressions.

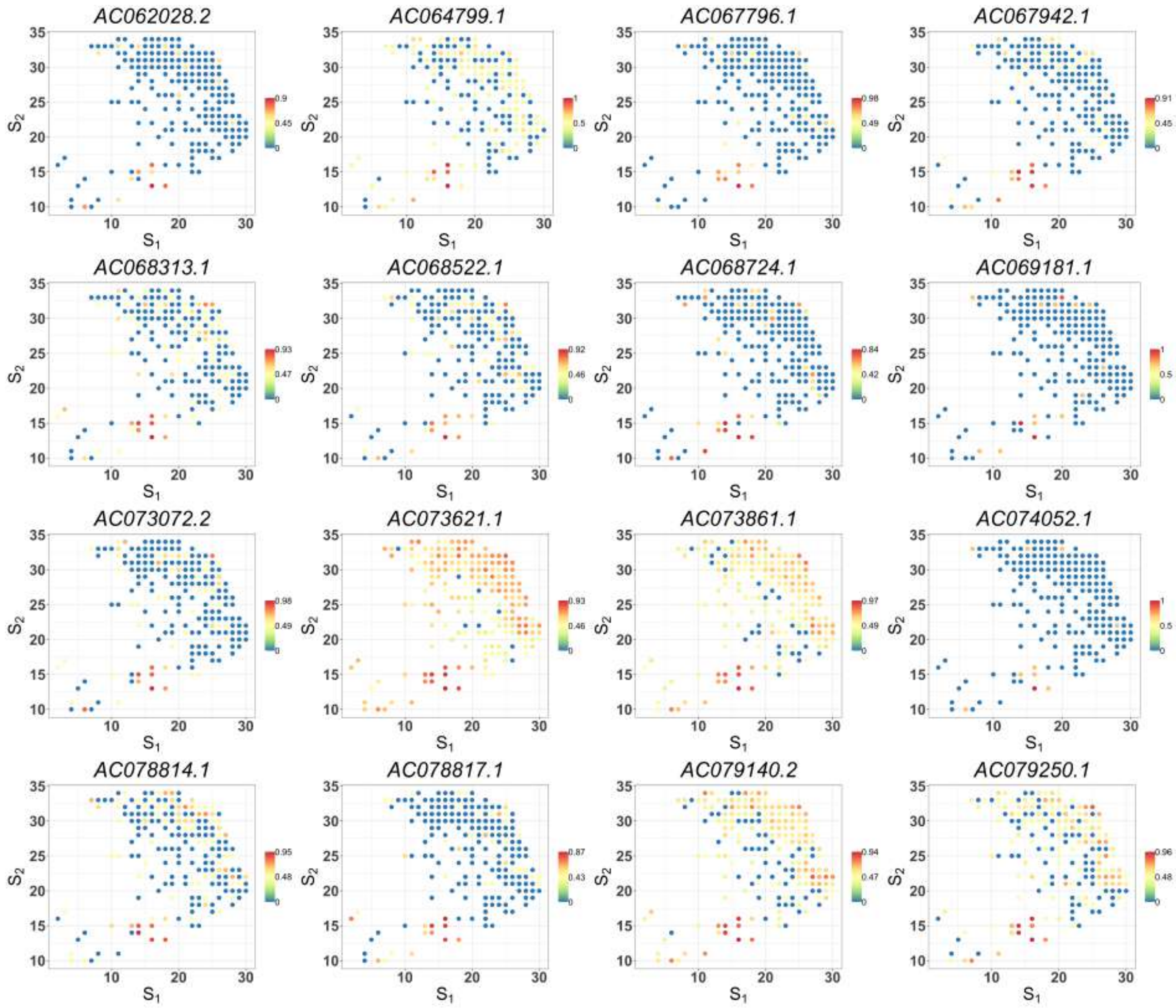


Figure S8: Spatial expression patterns of cancer-cell-specific SV genes detected by SPARK-X in the cancer region of PDAC data (part 7). Values are relative expressions.

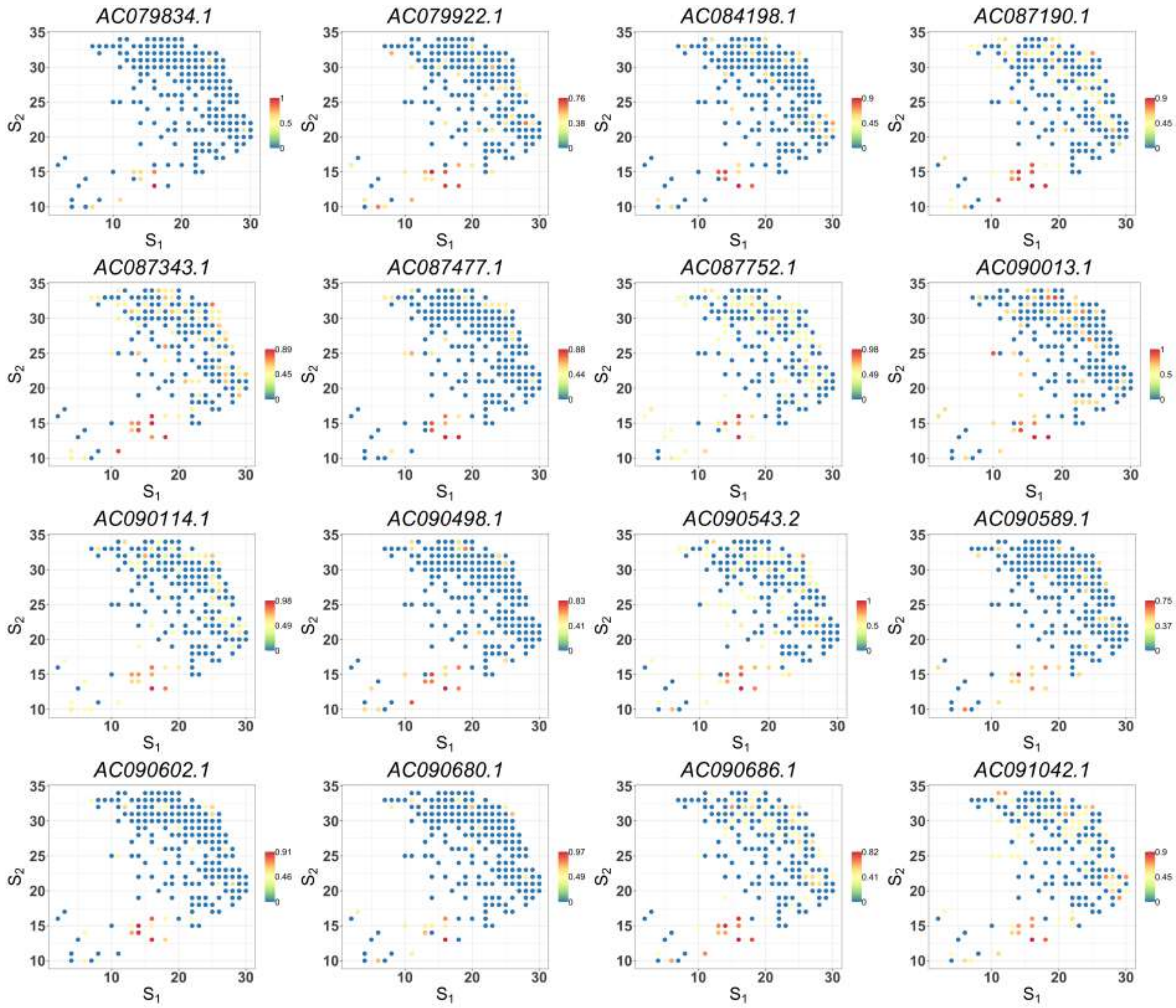


Figure S8: Spatial expression patterns of cancer-cell-specific SV genes detected by SPARK-X in the cancer region of PDAC data (part 8). Values are relative expressions.

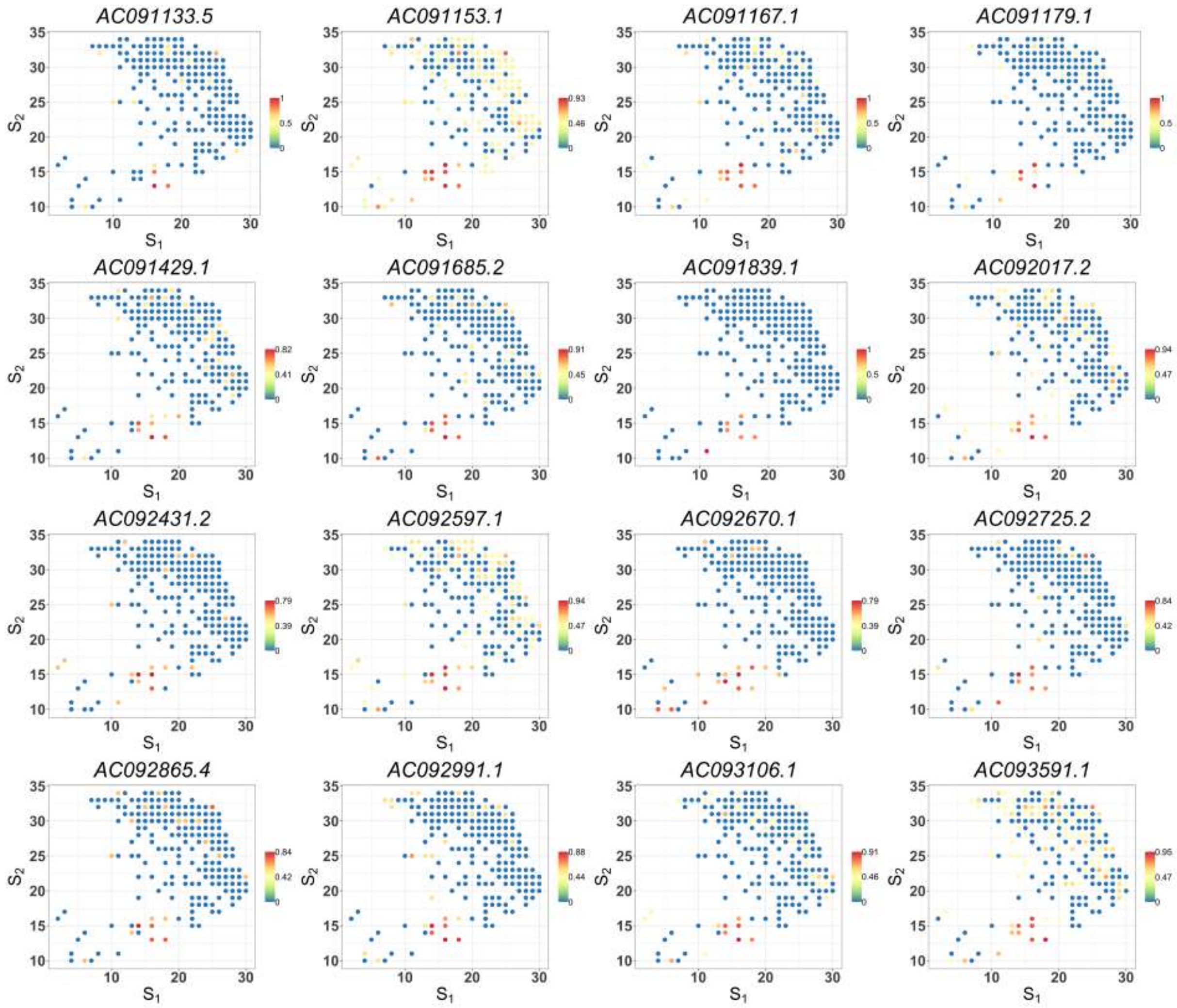


Figure S8: Spatial expression patterns of cancer-cell-specific SV genes detected by SPARK-X in the cancer region of PDAC data (part 9). Values are relative expressions.

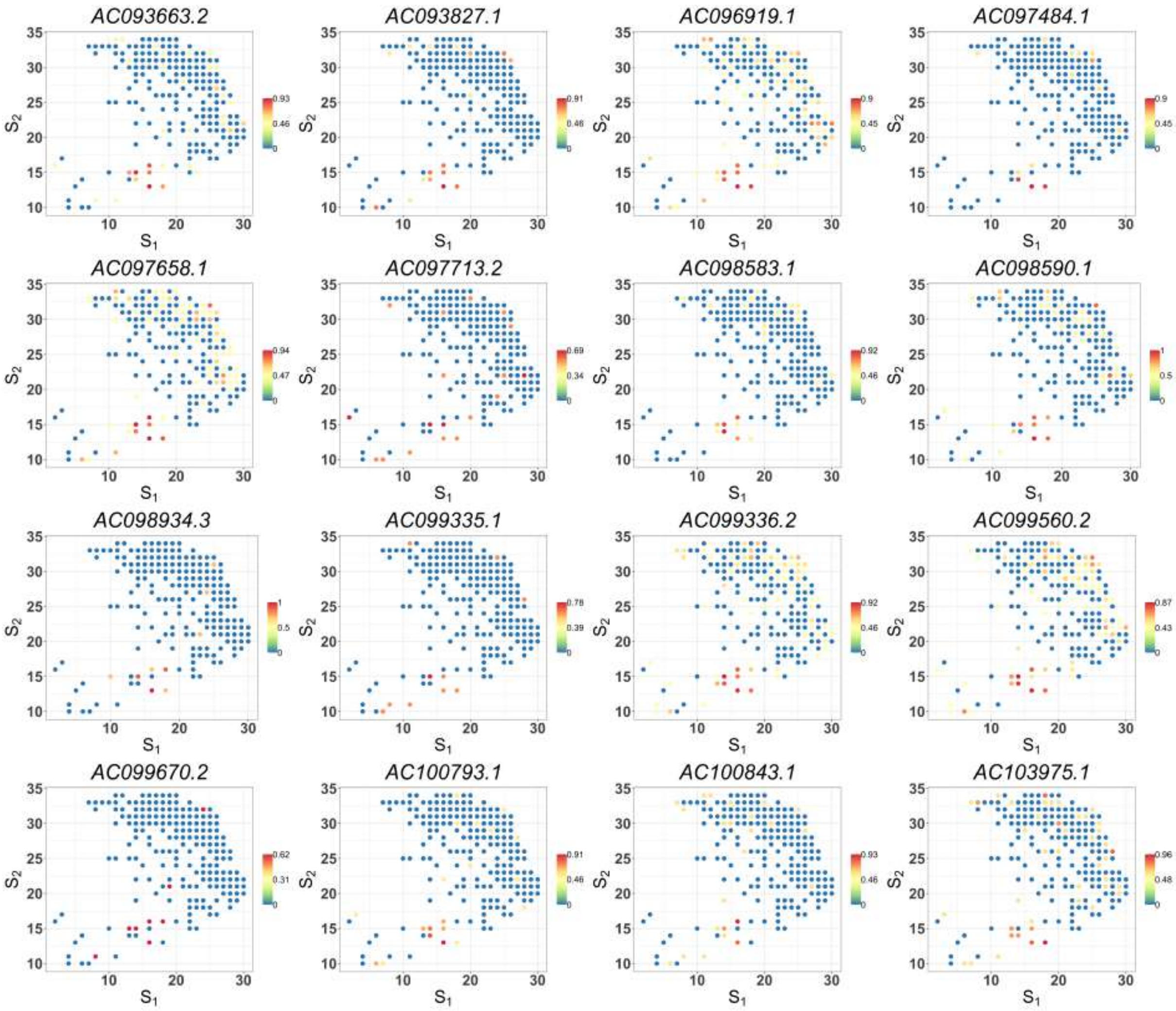


Figure S8: Spatial expression patterns of cancer-cell-specific SV genes detected by SPARK-X in the cancer region of PDAC data (part 10). Values are relative expressions.

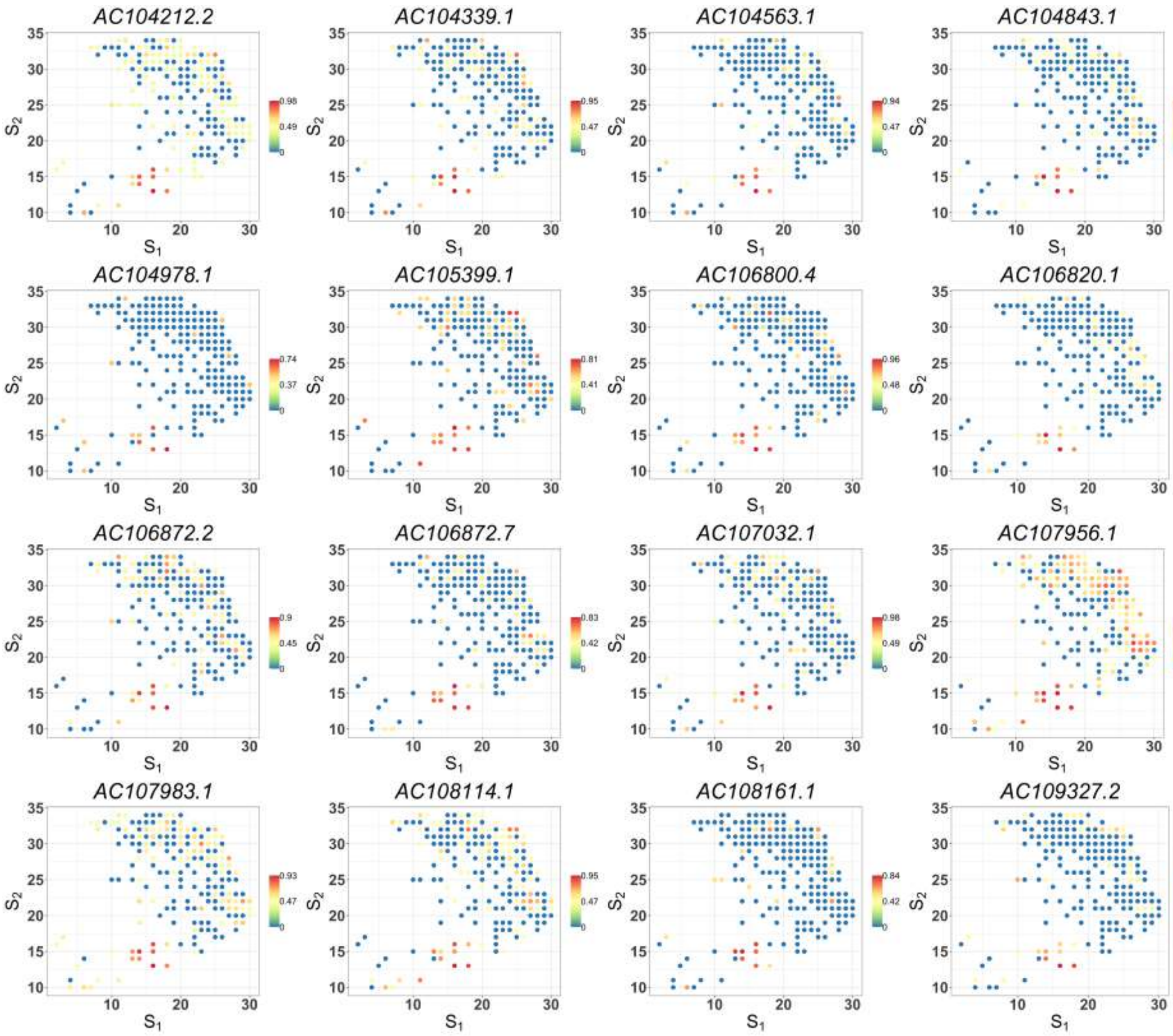


Figure S8: Spatial expression patterns of cancer-cell-specific SV genes detected by SPARK-X in the cancer region of PDAC data (part 11). Values are relative expressions.

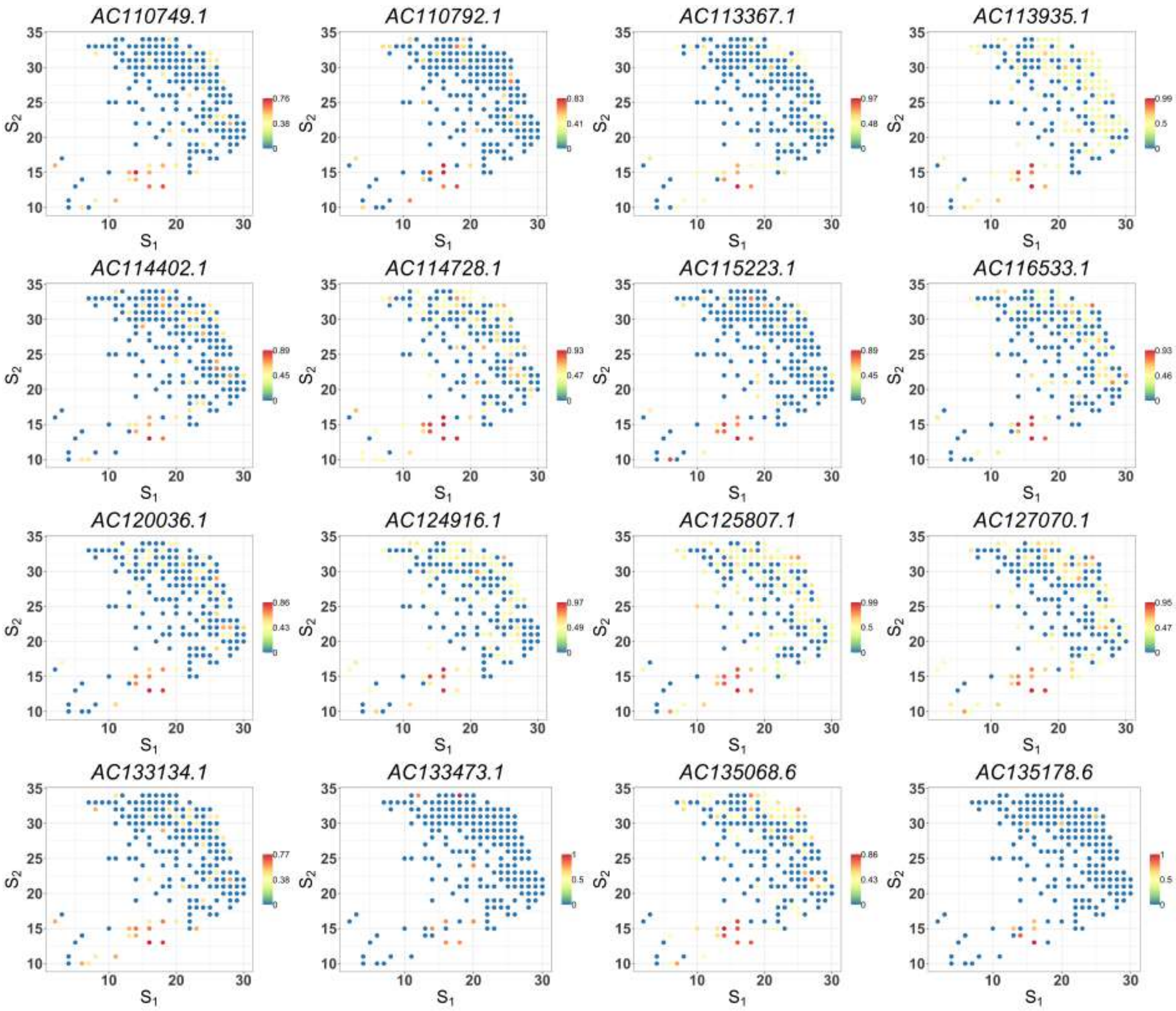


Figure S8: Spatial expression patterns of cancer-cell-specific SV genes detected by SPARK-X in the cancer region of PDAC data (part 12). Values are relative expressions.

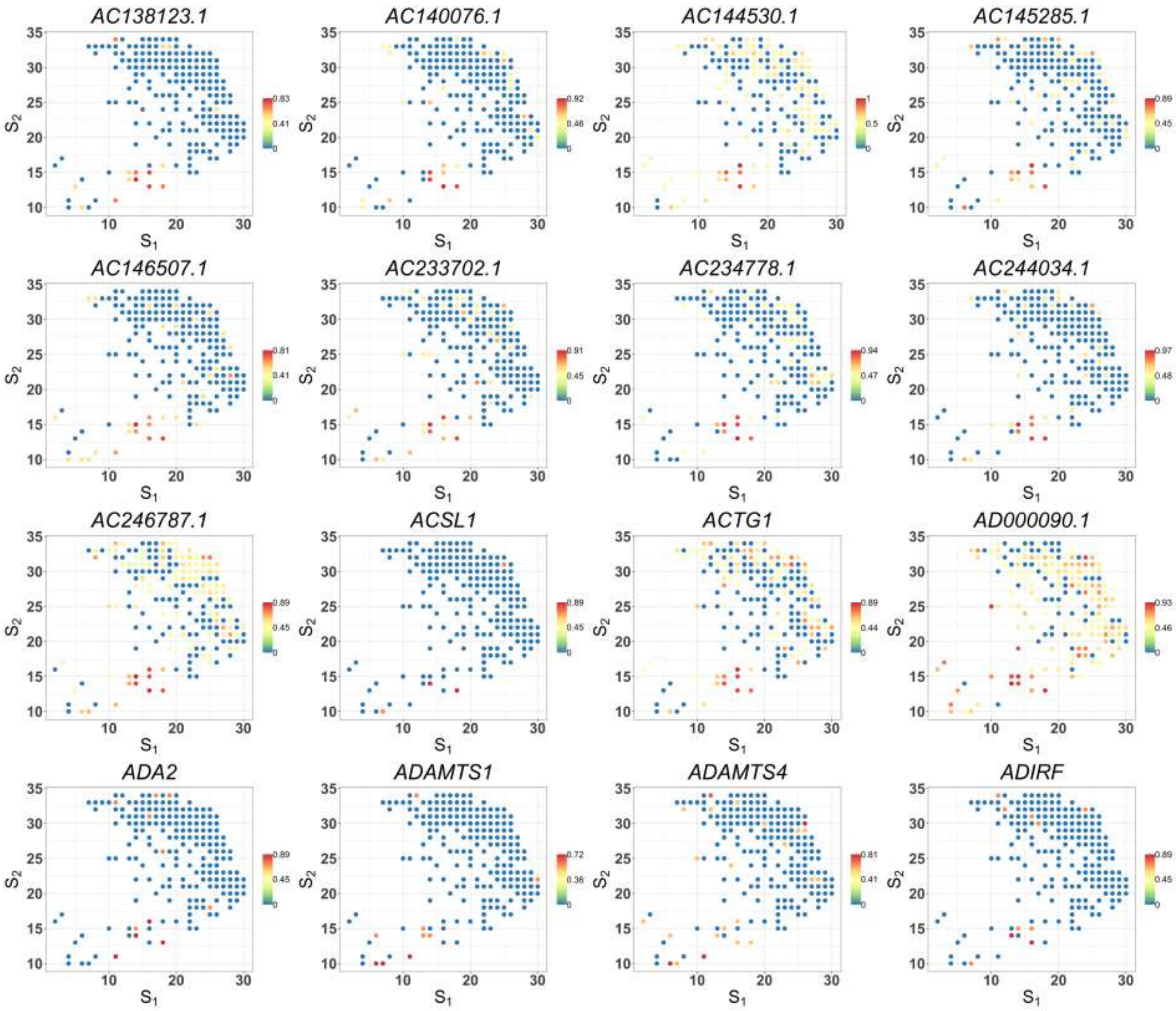


Figure S8: Spatial expression patterns of cancer-cell-specific SV genes detected by SPARK-X in the cancer region of PDAC data (part 13). Values are relative expressions.

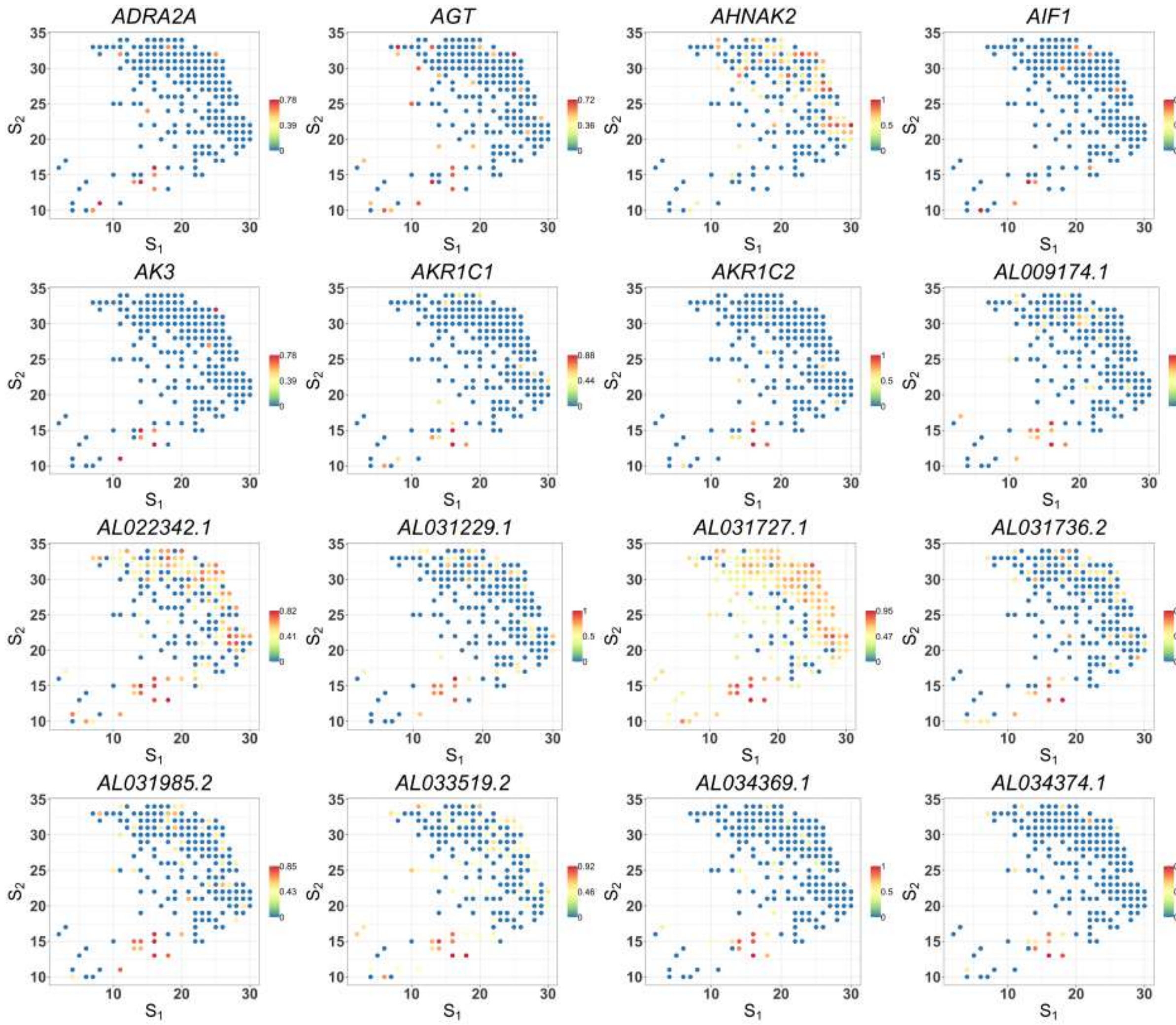


Figure S8: Spatial expression patterns of cancer-cell-specific SV genes detected by SPARK-X in the cancer region of PDAC data (part 14). Values are relative expressions.

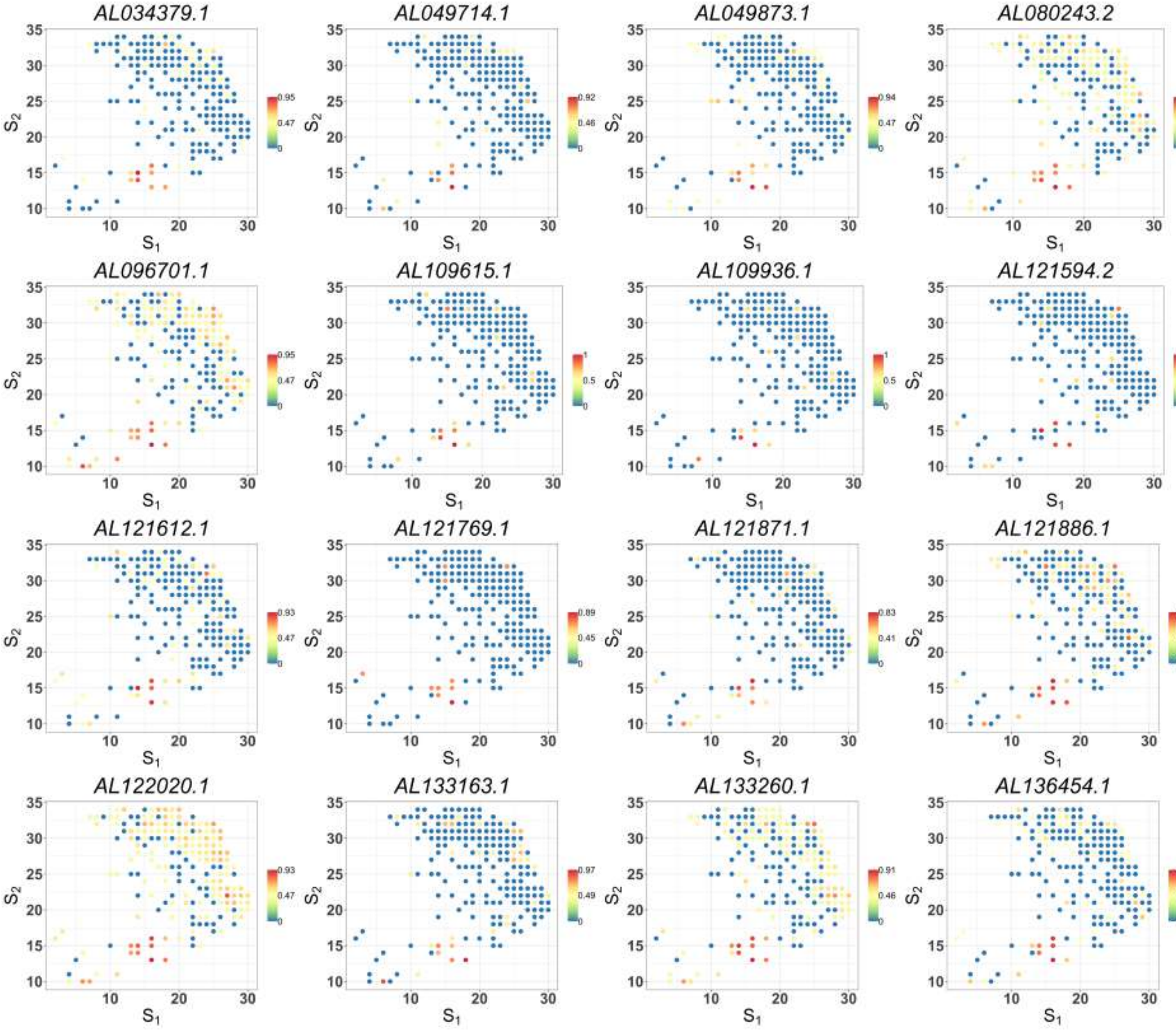


Figure S8: Spatial expression patterns of cancer-cell-specific SV genes detected by SPARK-X in the cancer region of PDAC data (part 15). Values are relative expressions.

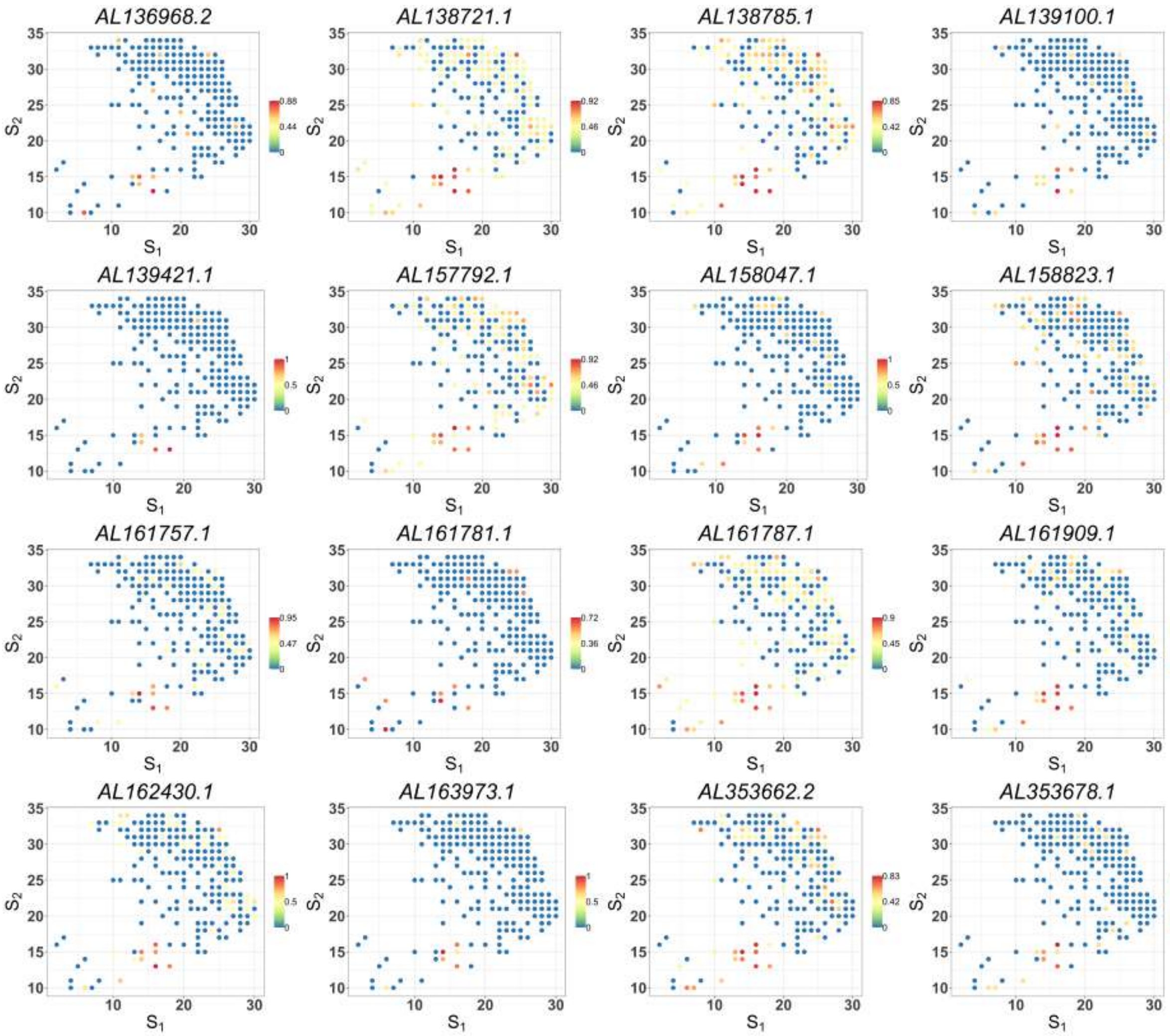


Figure S8: Spatial expression patterns of cancer-cell-specific SV genes detected by SPARK-X in the cancer region of PDAC data (part 16). Values are relative expressions.

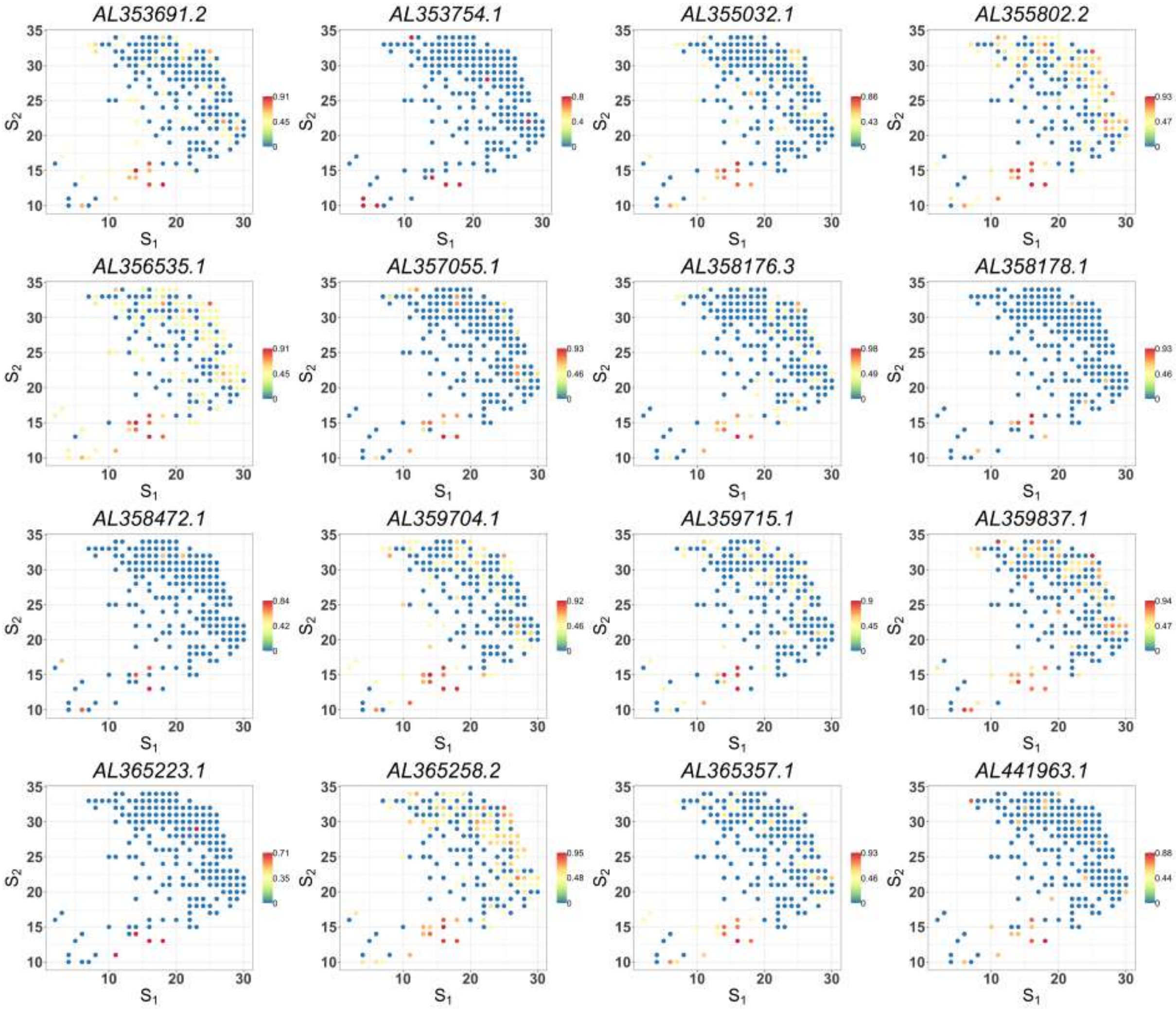


Figure S8: Spatial expression patterns of cancer-cell-specific SV genes detected by SPARK-X in the cancer region of PDAC data (part 17). Values are relative expressions.

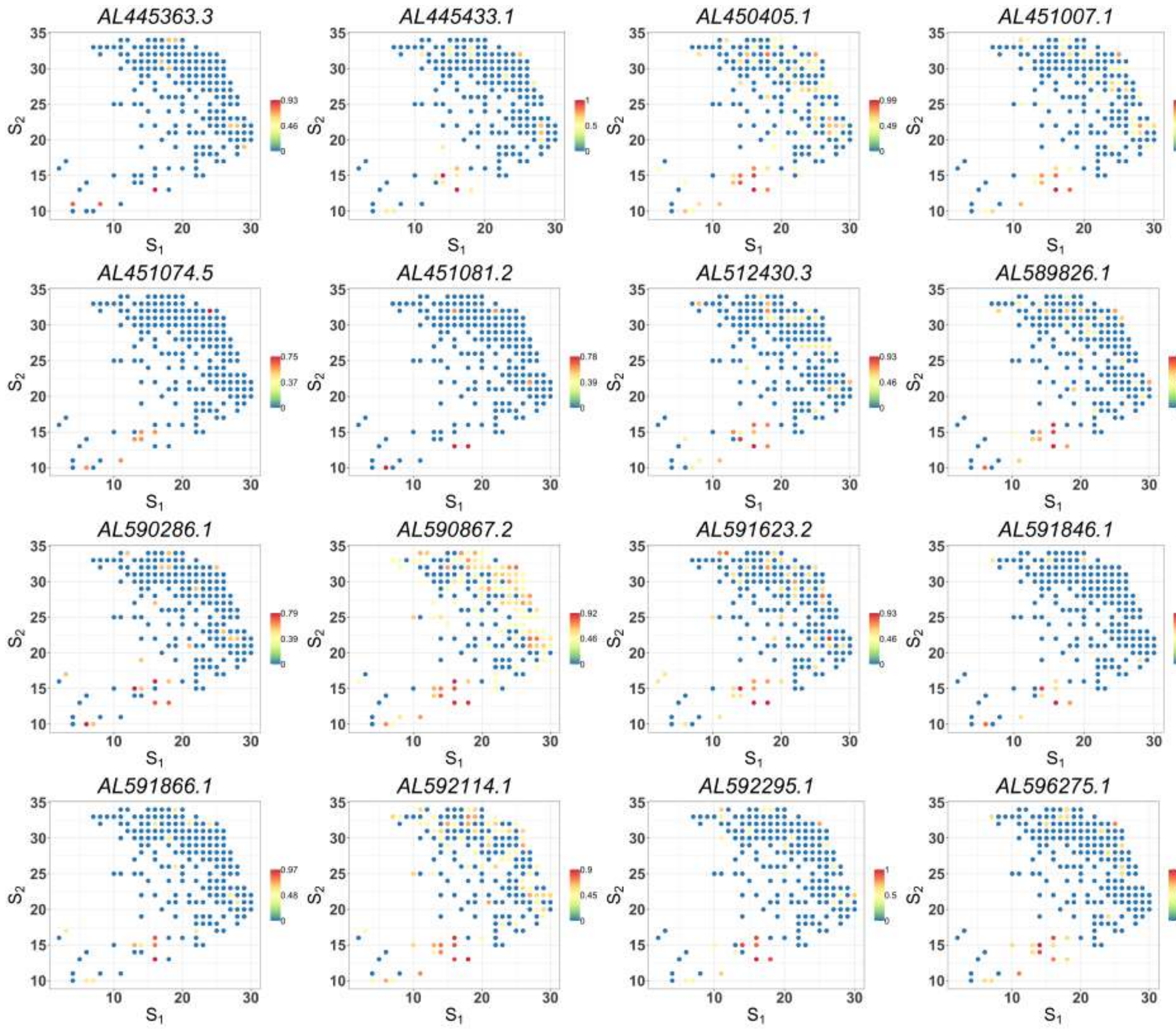


Figure S8: Spatial expression patterns of cancer-cell-specific SV genes detected by SPARK-X in the cancer region of PDAC data (part 18). Values are relative expressions.

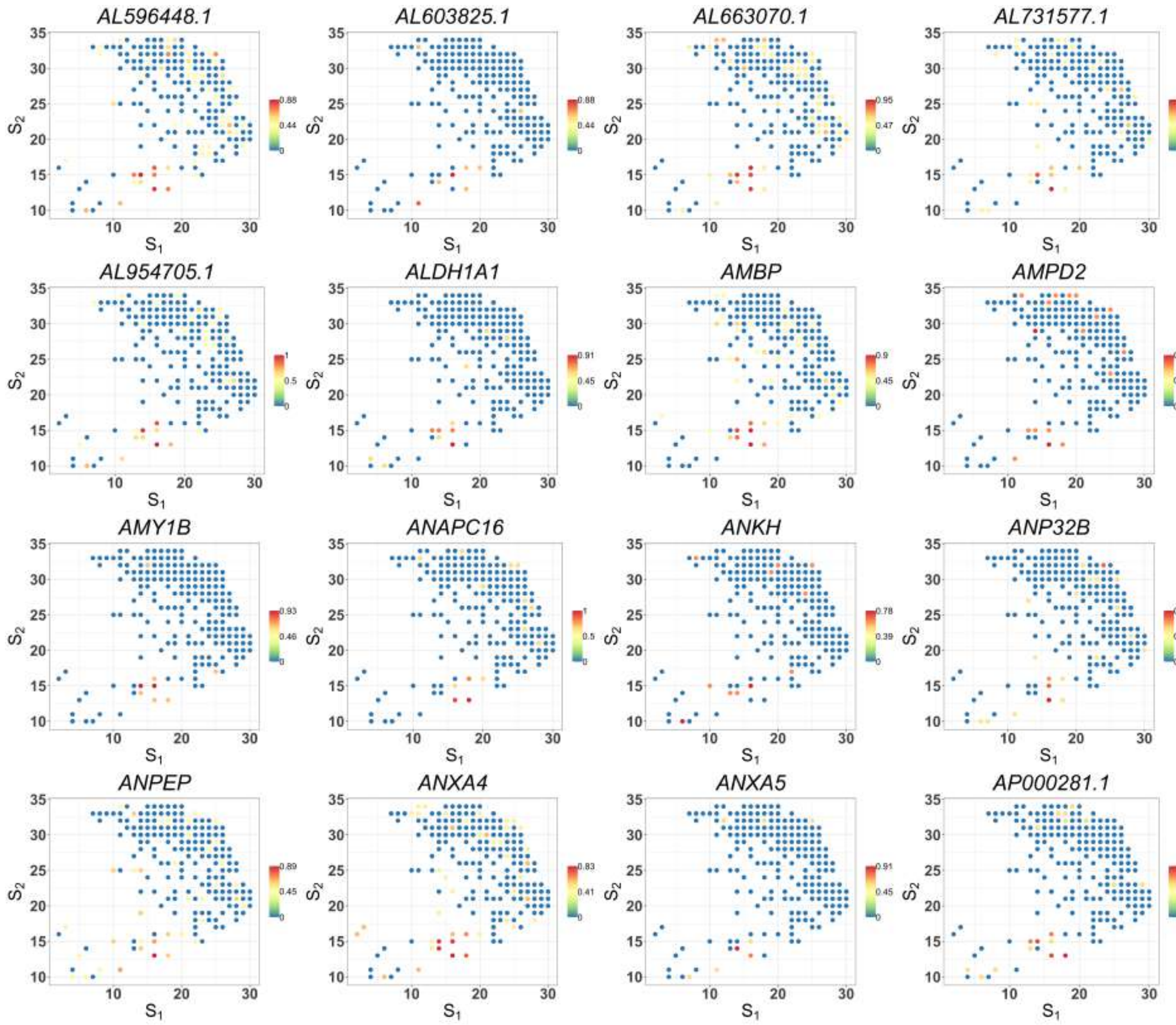


Figure S8: Spatial expression patterns of cancer-cell-specific SV genes detected by SPARK-X in the cancer region of PDAC data (part 19). Values are relative expressions.

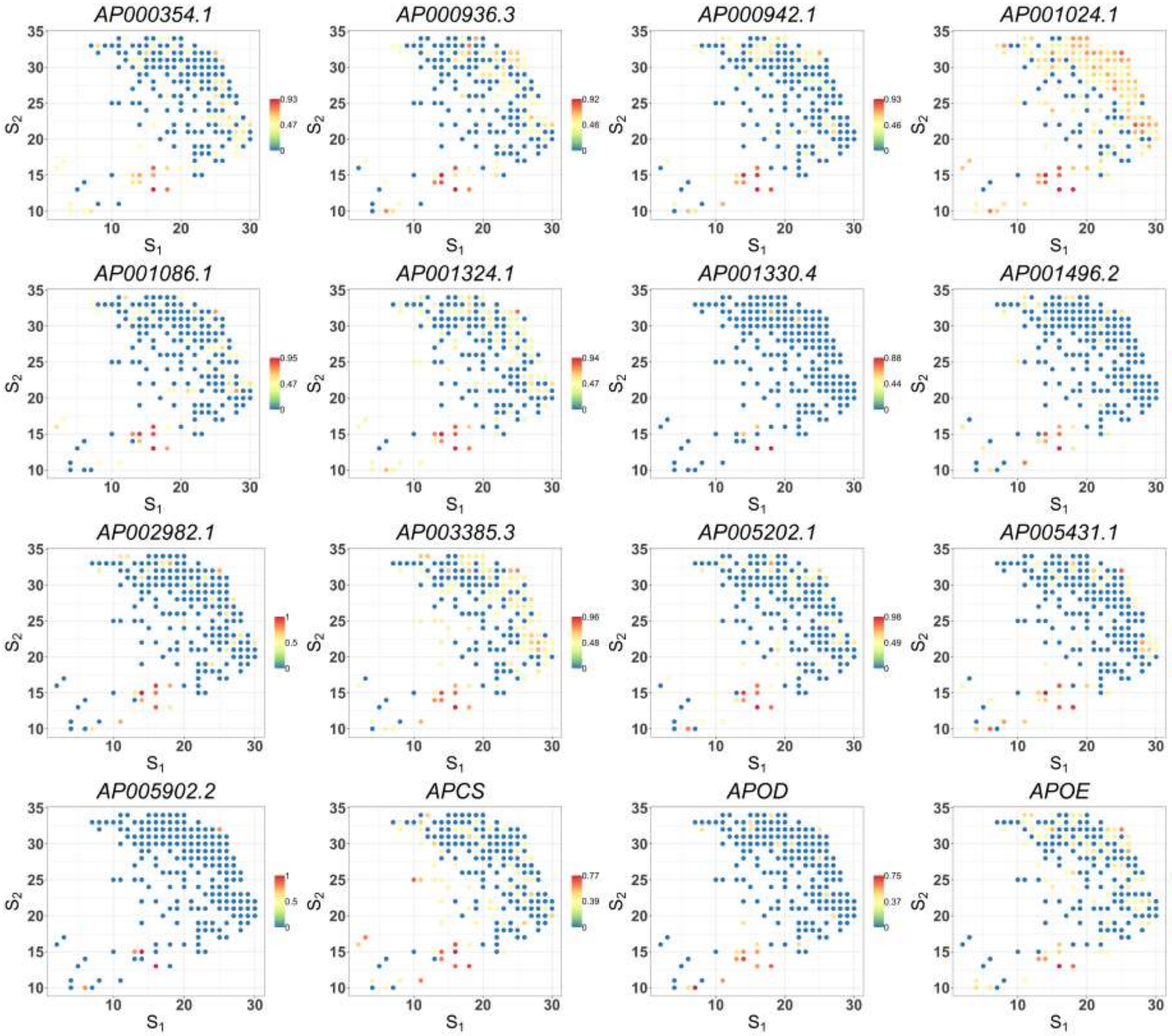


Figure S8: Spatial expression patterns of cancer-cell-specific SV genes detected by SPARK-X in the cancer region of PDAC data (part 20). Values are relative expressions.

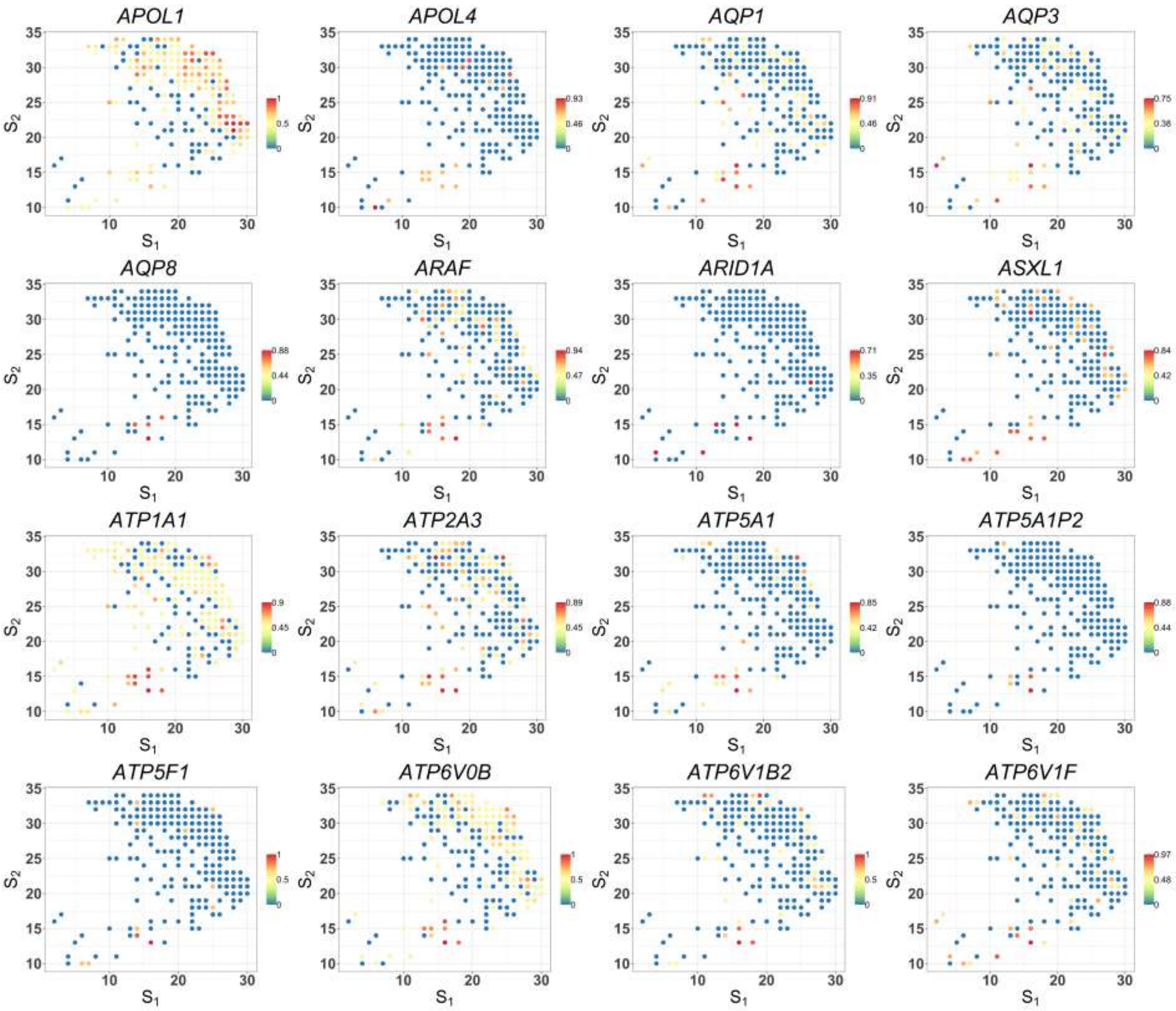


Figure S8: Spatial expression patterns of cancer-cell-specific SV genes detected by SPARK-X in the cancer region of PDAC data (part 21). Values are relative expressions.

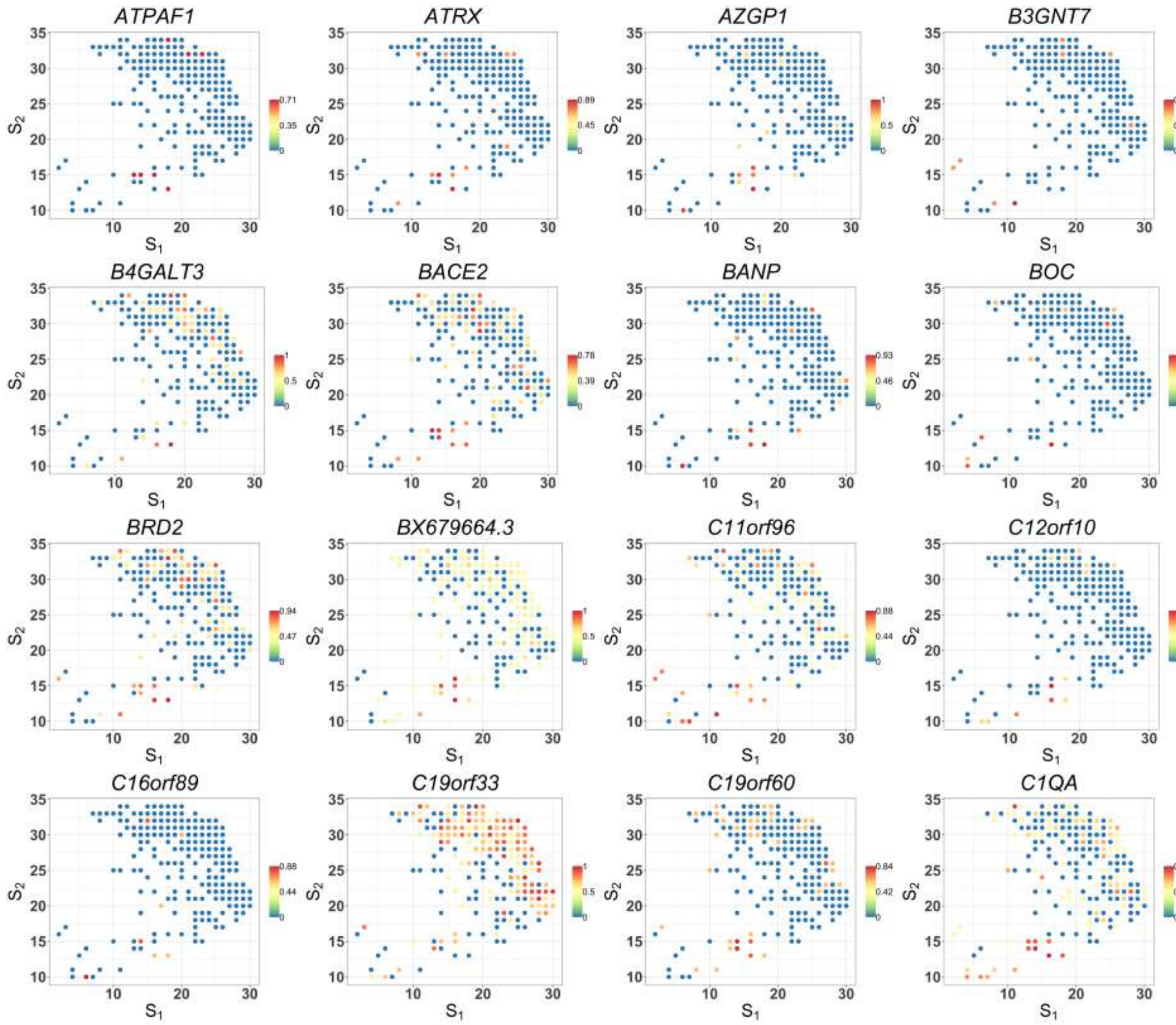


Figure S8: Spatial expression patterns of cancer-cell-specific SV genes detected by SPARK-X in the cancer region of PDAC data (part 22). Values are relative expressions.

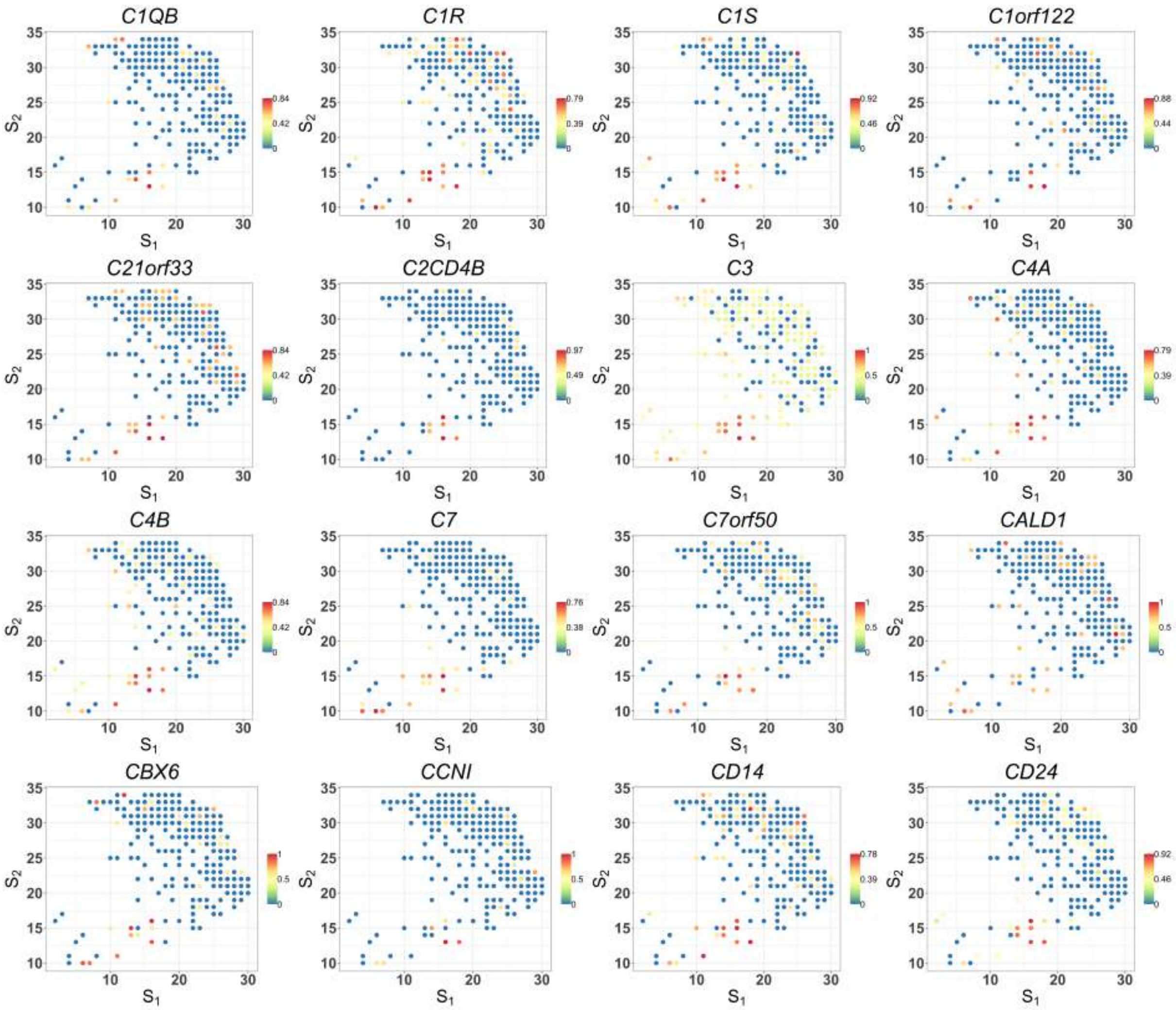


Figure S8: Spatial expression patterns of cancer-cell-specific SV genes detected by SPARK-X in the cancer region of PDAC data (part 23). Values are relative expressions.

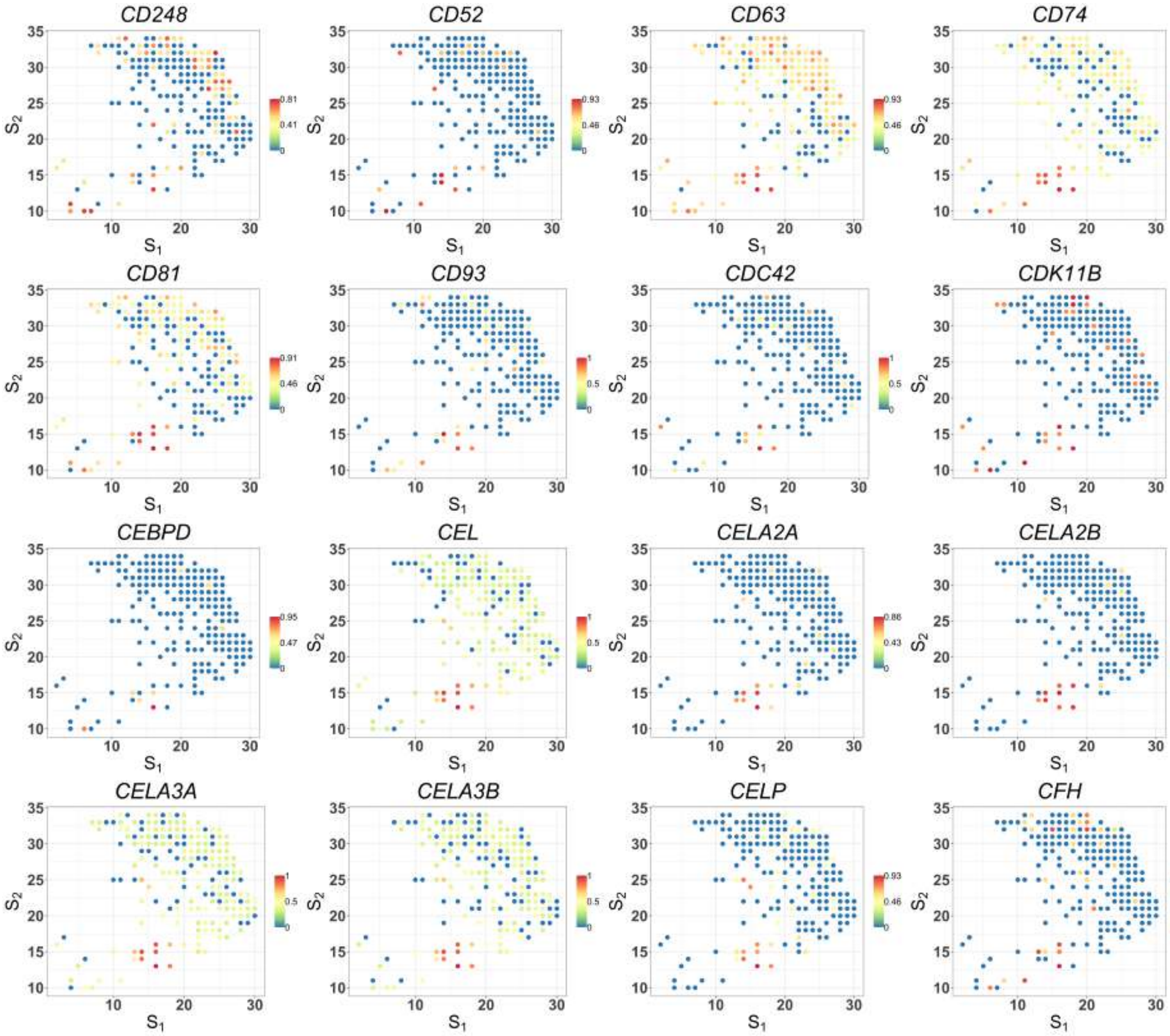


Figure S8: Spatial expression patterns of cancer-cell-specific SV genes detected by SPARK-X in the cancer region of PDAC data (part 24). Values are relative expressions.

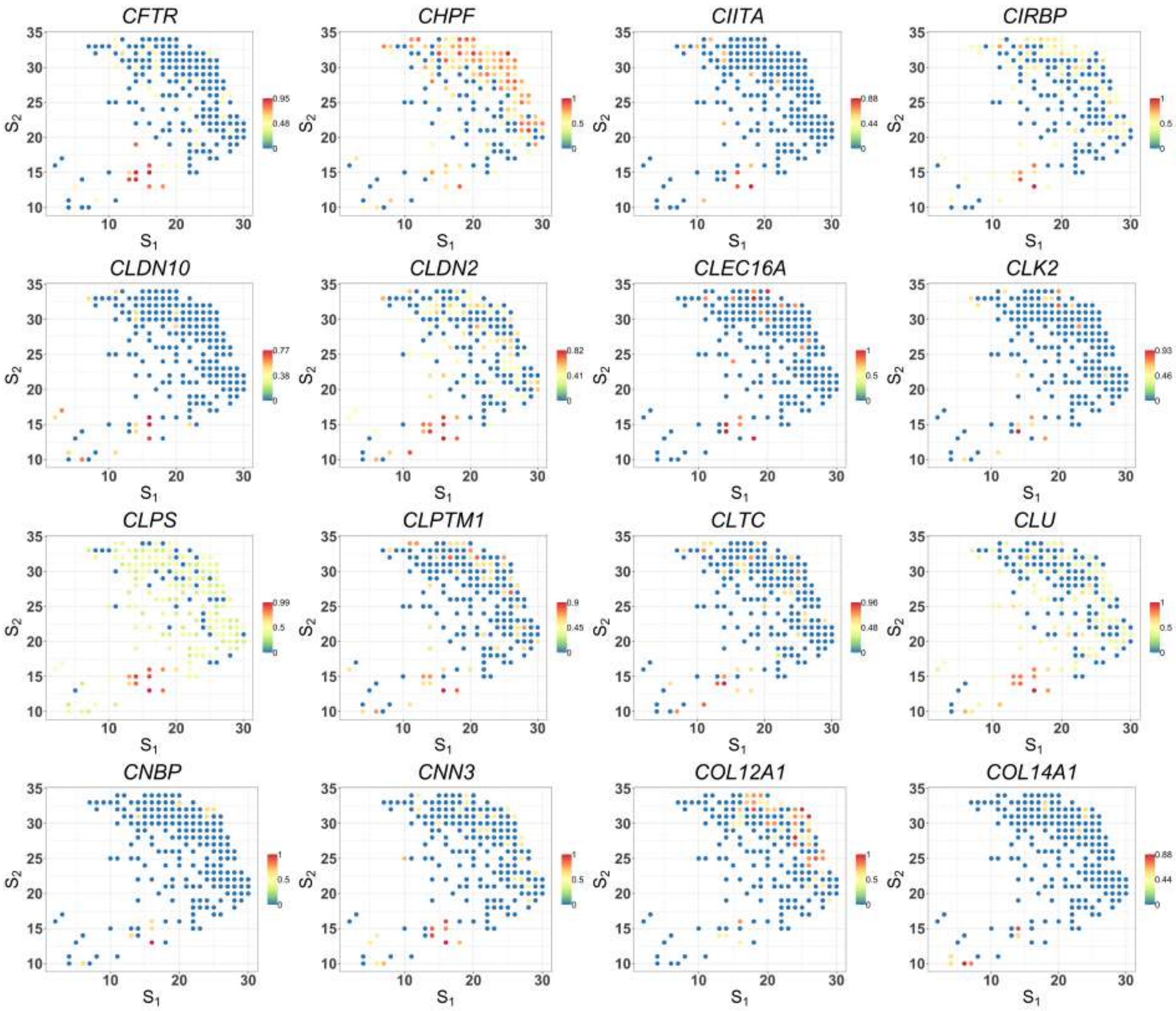


Figure S8: Spatial expression patterns of cancer-cell-specific SV genes detected by SPARK-X in the cancer region of PDAC data (part 25). Values are relative expressions.

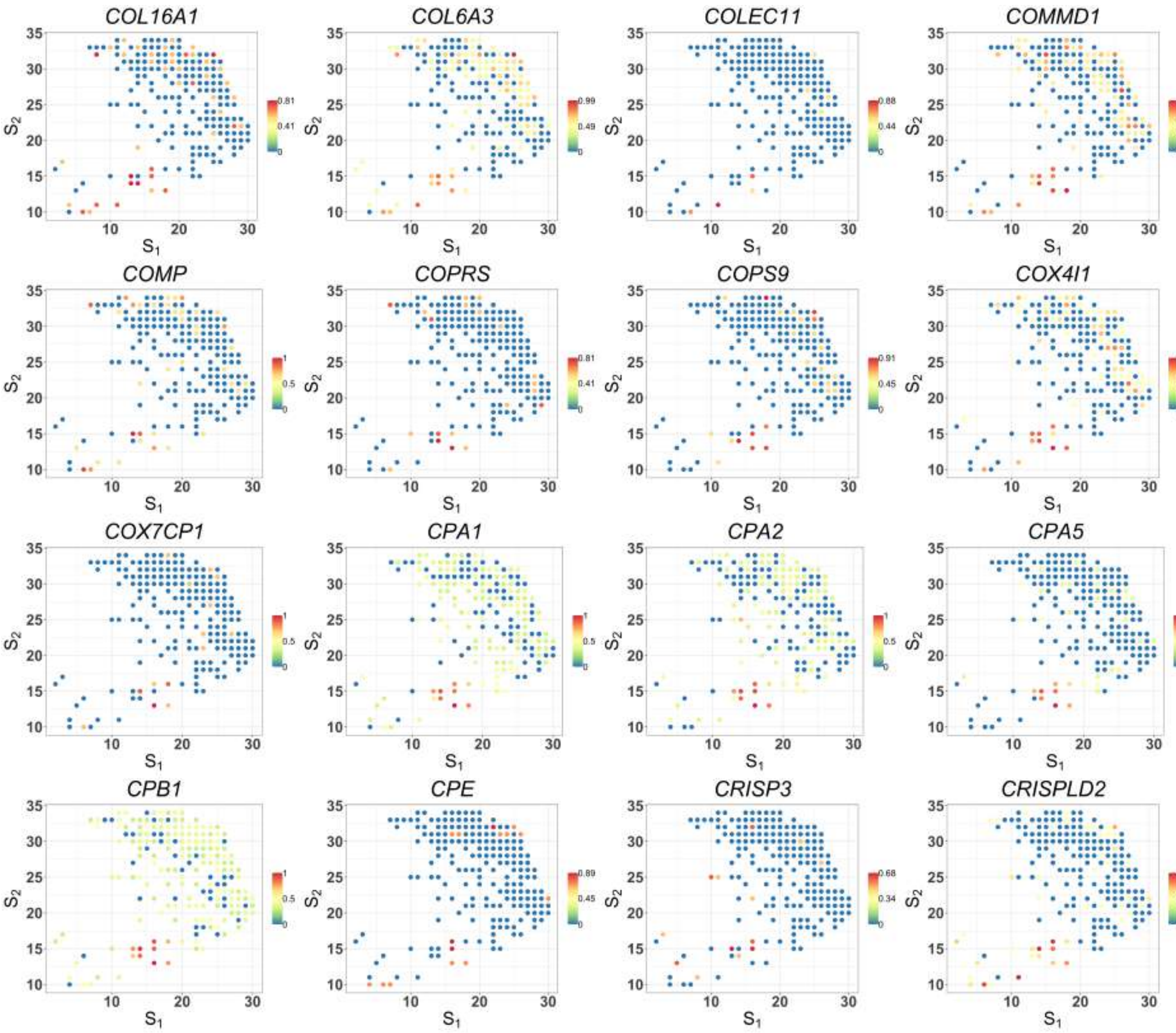


Figure S8: Spatial expression patterns of cancer-cell-specific SV genes detected by SPARK-X in the cancer region of PDAC data (part 26). Values are relative expressions.

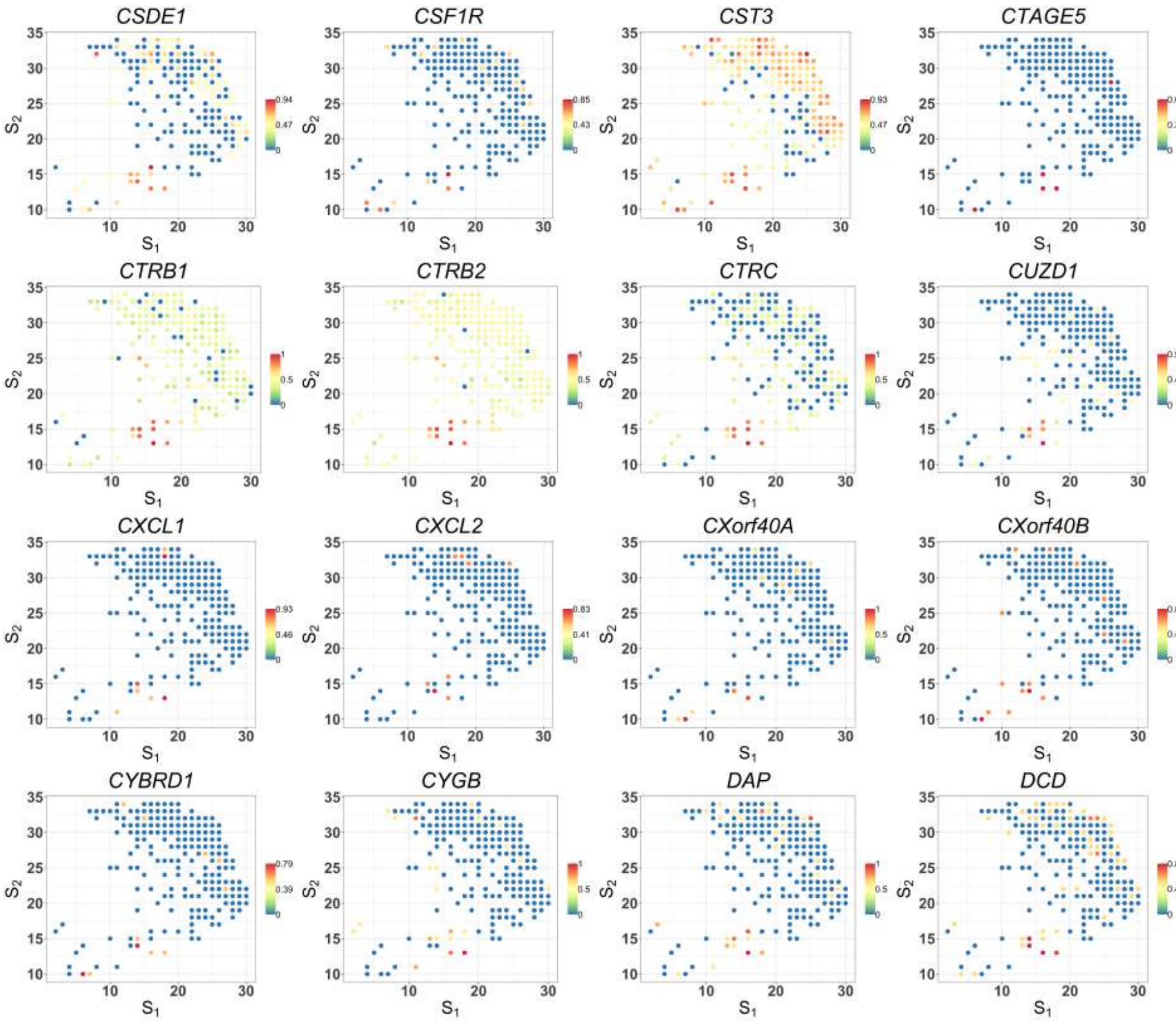


Figure S8: Spatial expression patterns of SV genes detected by SPARK-X in the cancer region of PDAC data (part 27). Values are relative expressions.

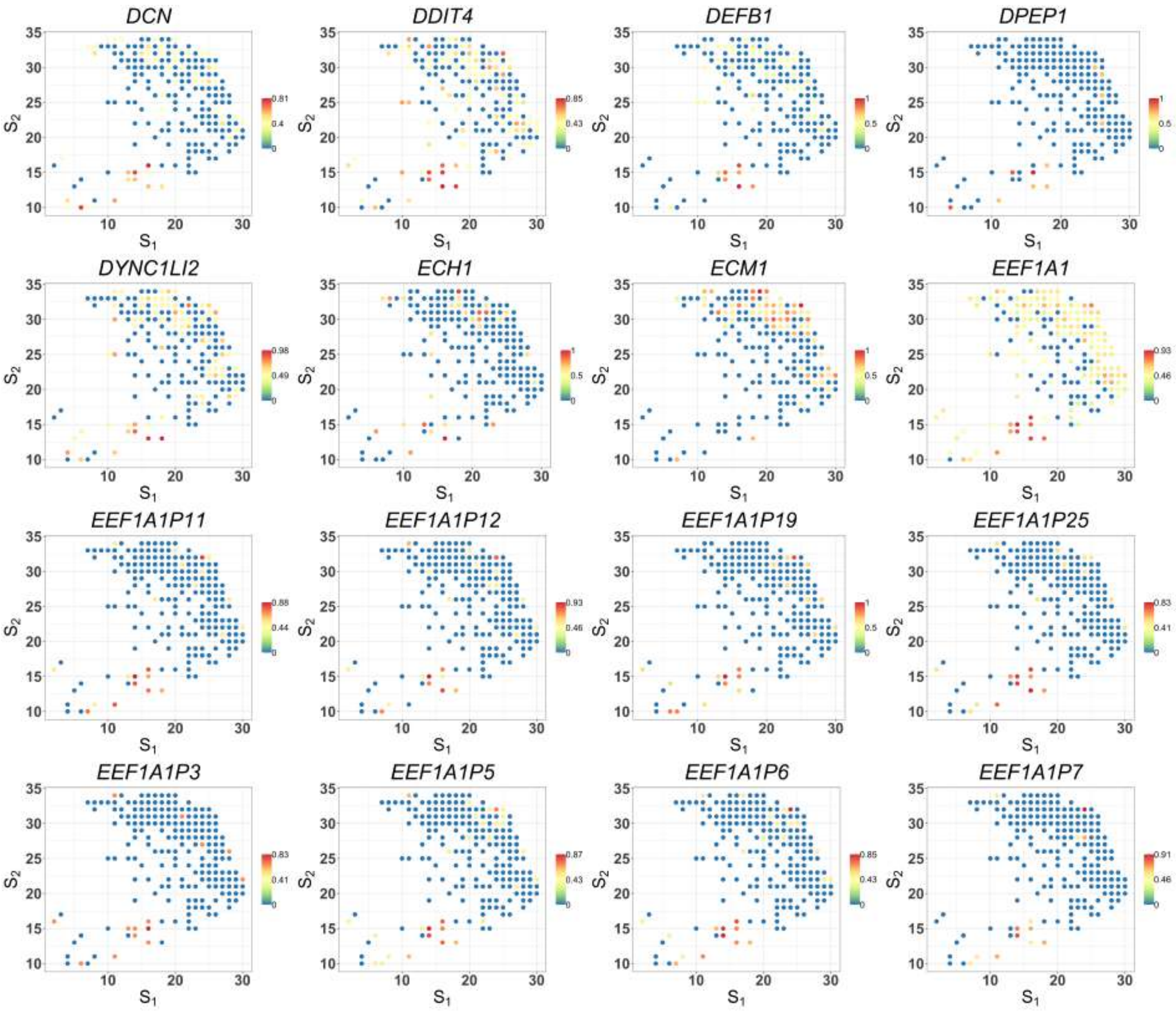


Figure S8: Spatial expression patterns of cancer-cell-specific SV genes detected by SPARK-X in the cancer region of PDAC data (part 28). Values are relative expressions.

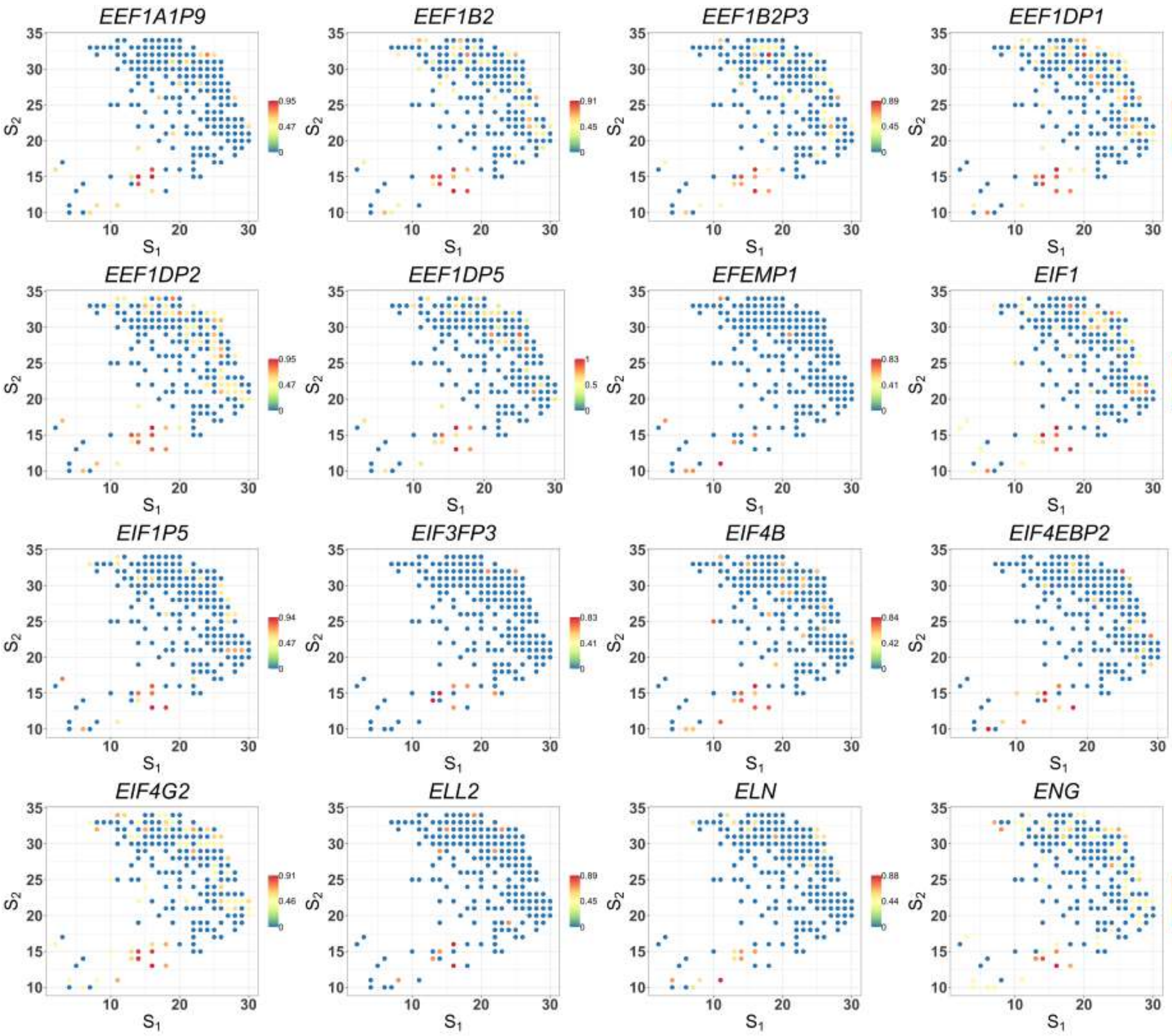


Figure S8: Spatial expression patterns of cancer-cell-specific SV genes detected by SPARK-X in the cancer region of PDAC data (part 29). Values are relative expressions.

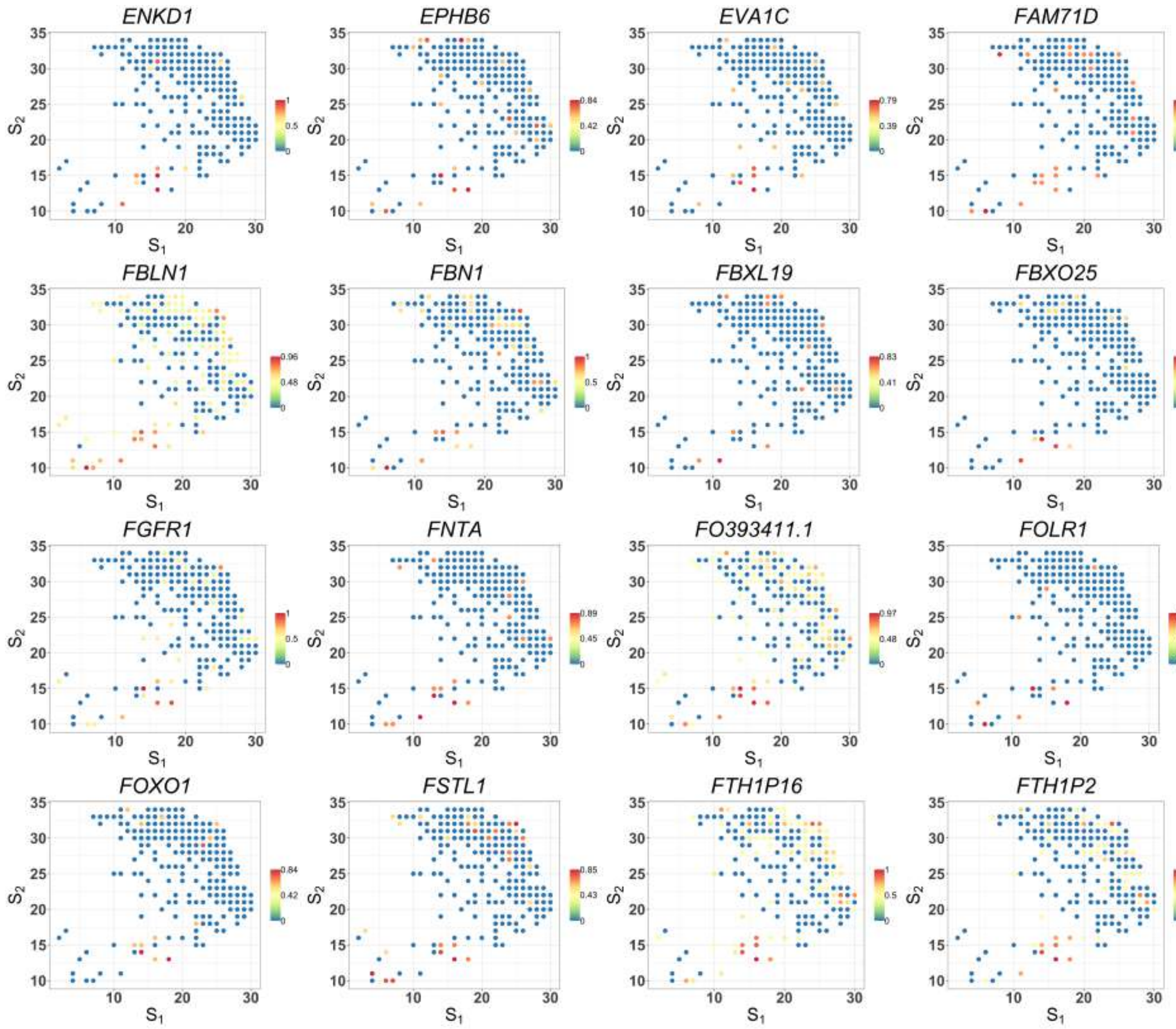


Figure S8: Spatial expression patterns of cancer-cell-specific SV genes detected by SPARK-X in the cancer region of PDAC data (part 30). Values are relative expressions.

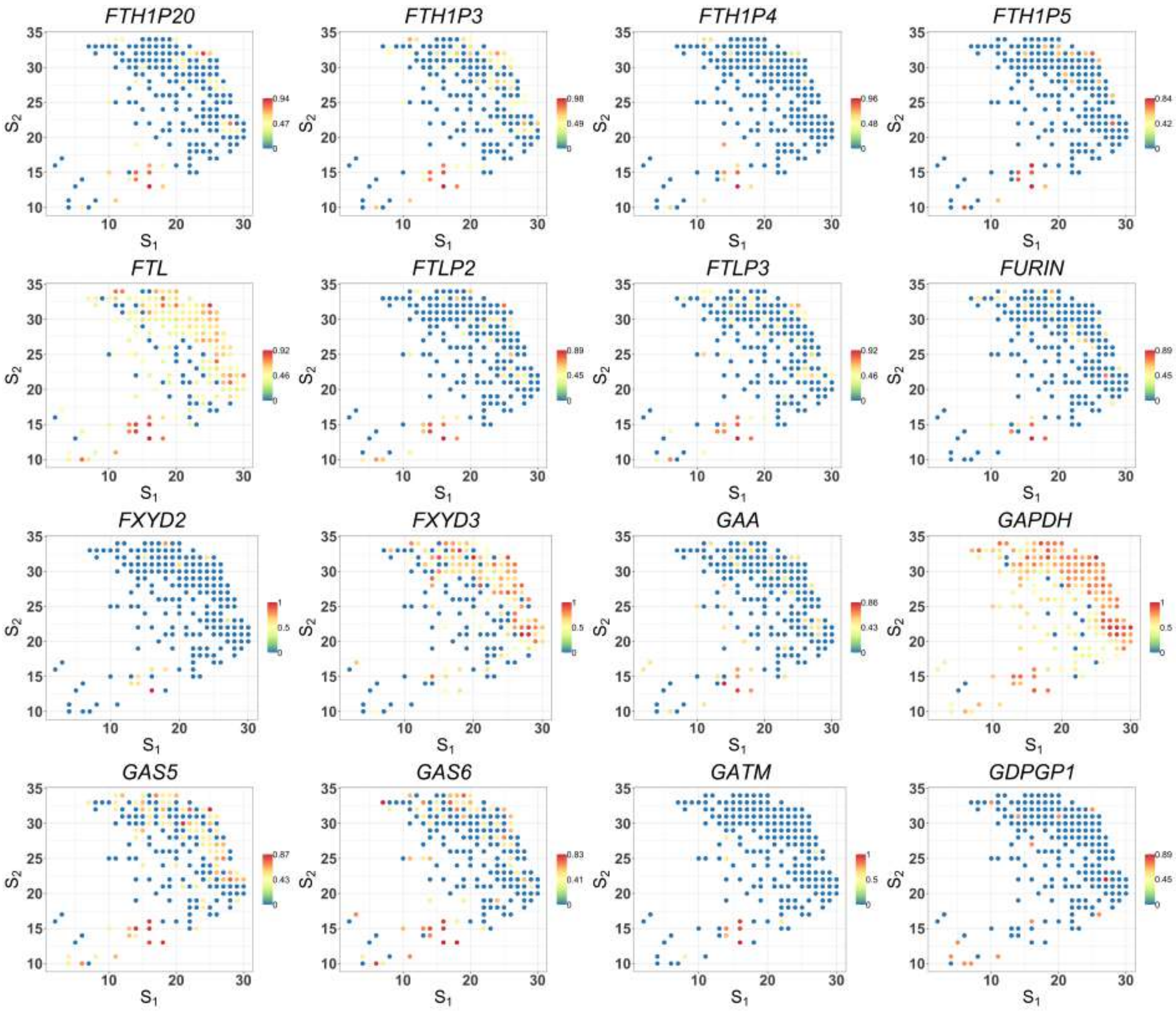


Figure S8: Spatial expression patterns of cancer-cell-specific SV genes detected by SPARK-X in the cancer region of PDAC data (part 31). Values are relative expressions.

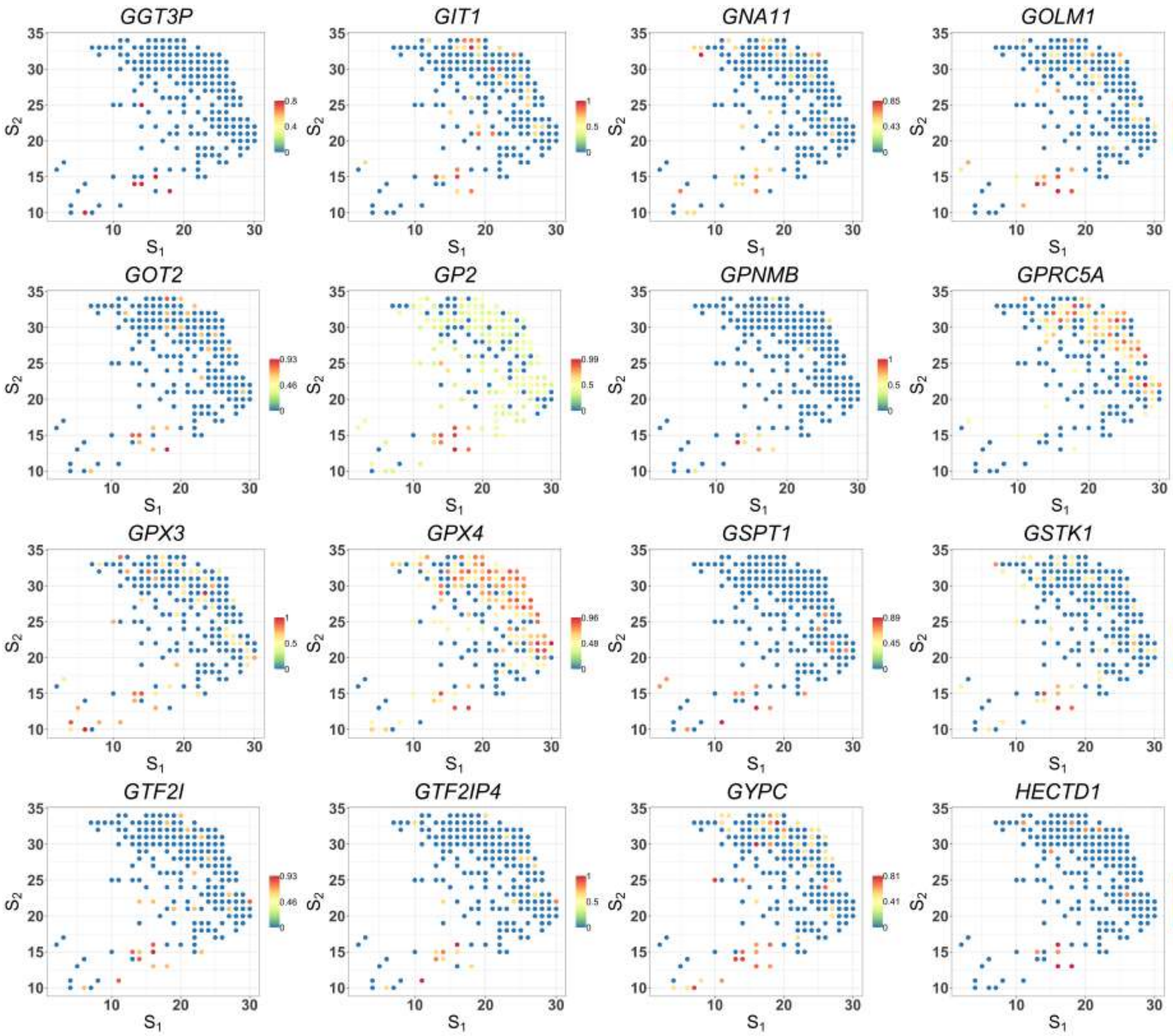


Figure S8: Spatial expression patterns of cancer-cell-specific SV genes detected by SPARK-X in the cancer region of PDAC data (part 32). Values are relative expressions.

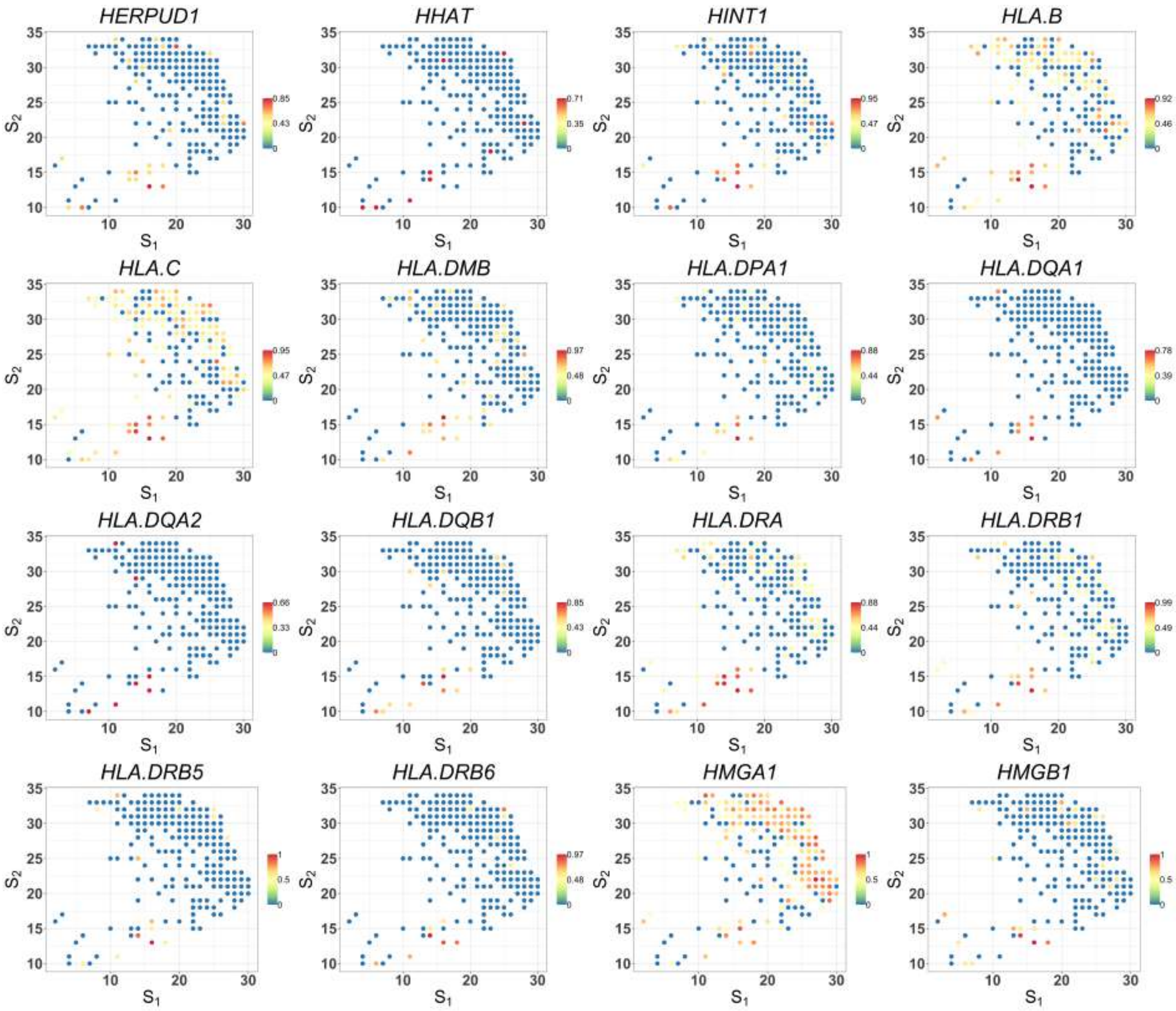


Figure S8: Spatial expression patterns of cancer-cell-specific SV genes detected by SPARK-X in the cancer region of PDAC data (part 33). Values are relative expressions.

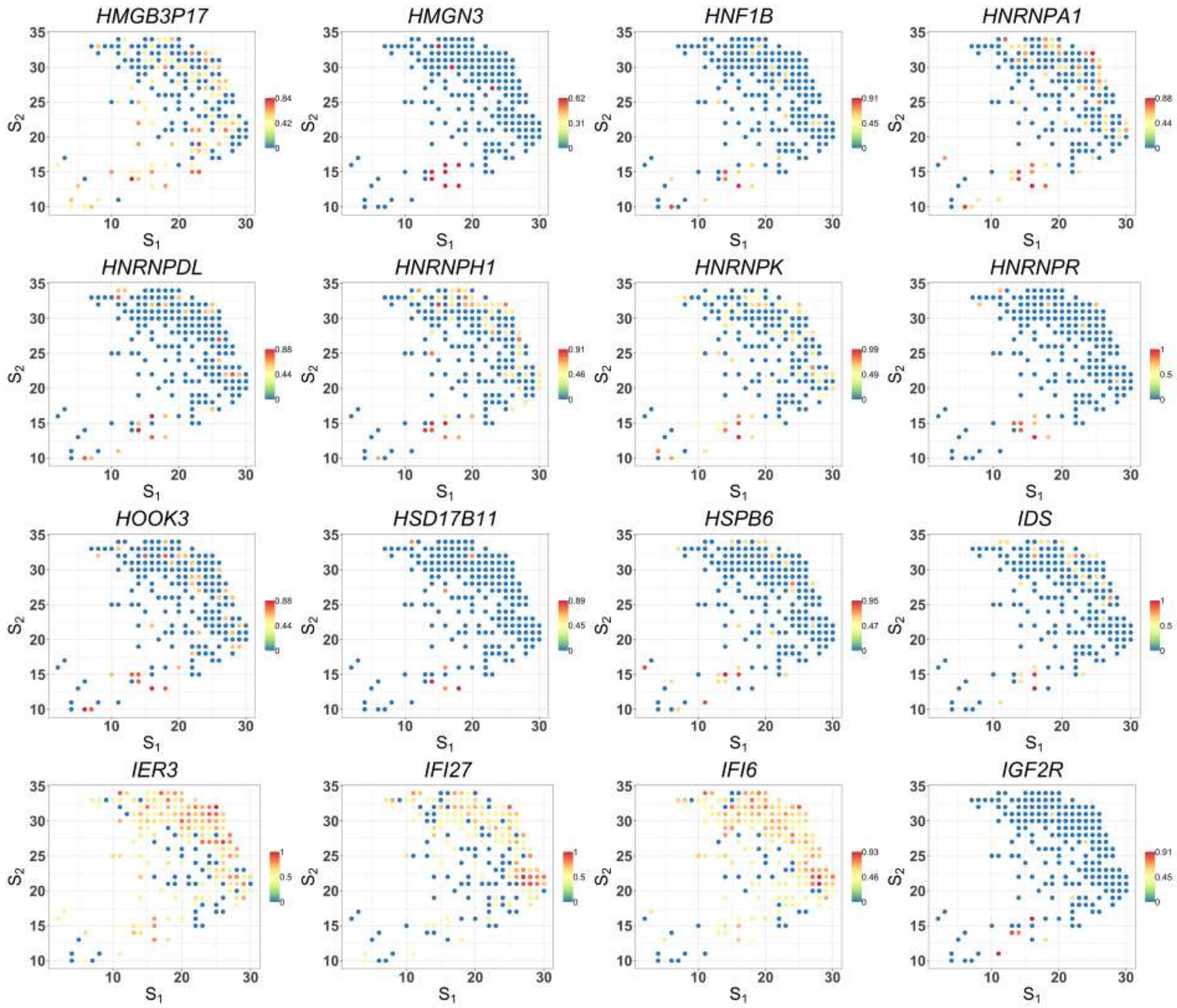


Figure S8: Spatial expression patterns of cancer-cell-specific SV genes detected by SPARK-X in the cancer region of PDAC data (part 34). Values are relative expressions.

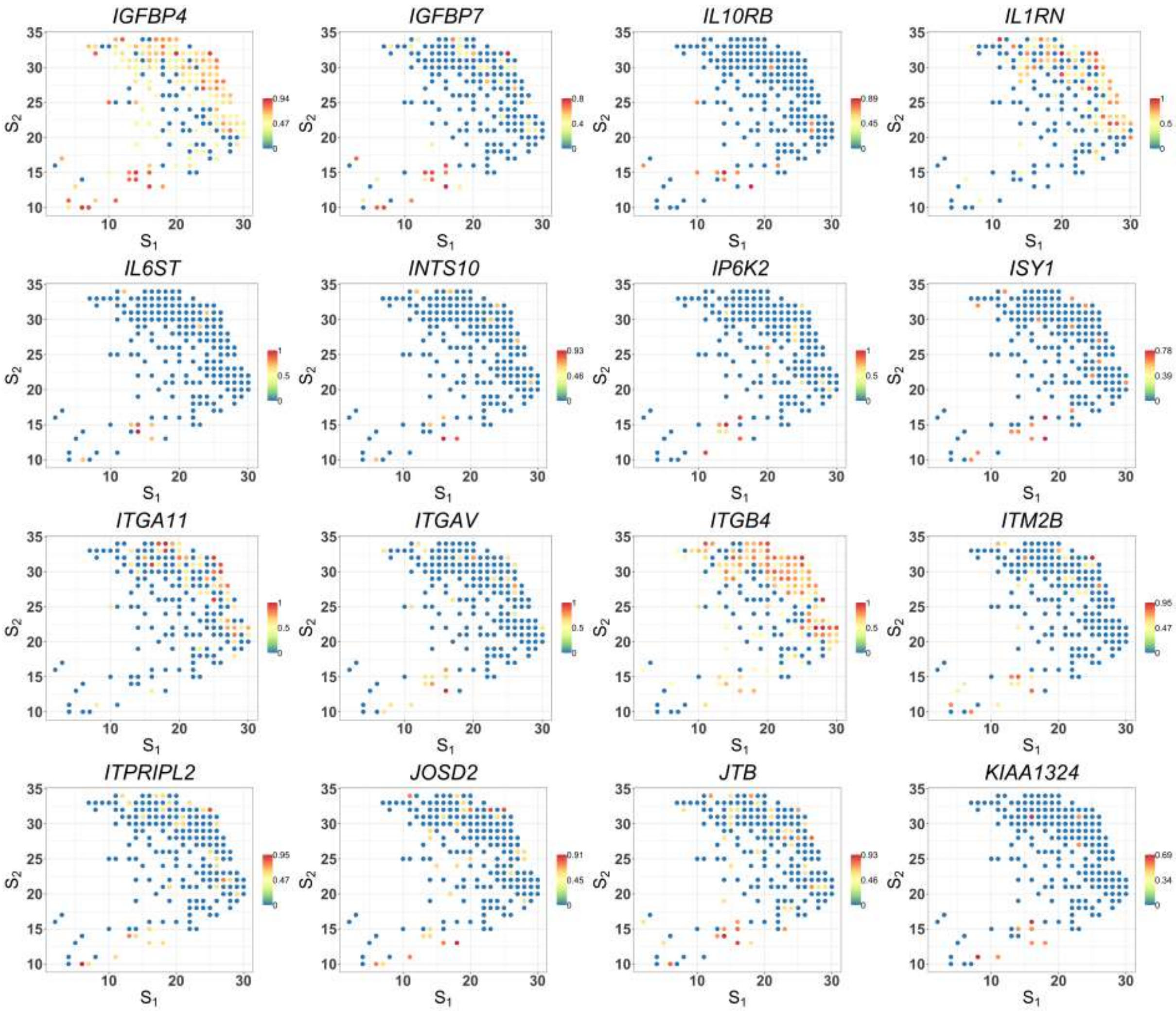


Figure S8: Spatial expression patterns of cancer-cell-specific SV genes detected by SPARK-X in the cancer region of PDAC data (part 35). Values are relative expressions.

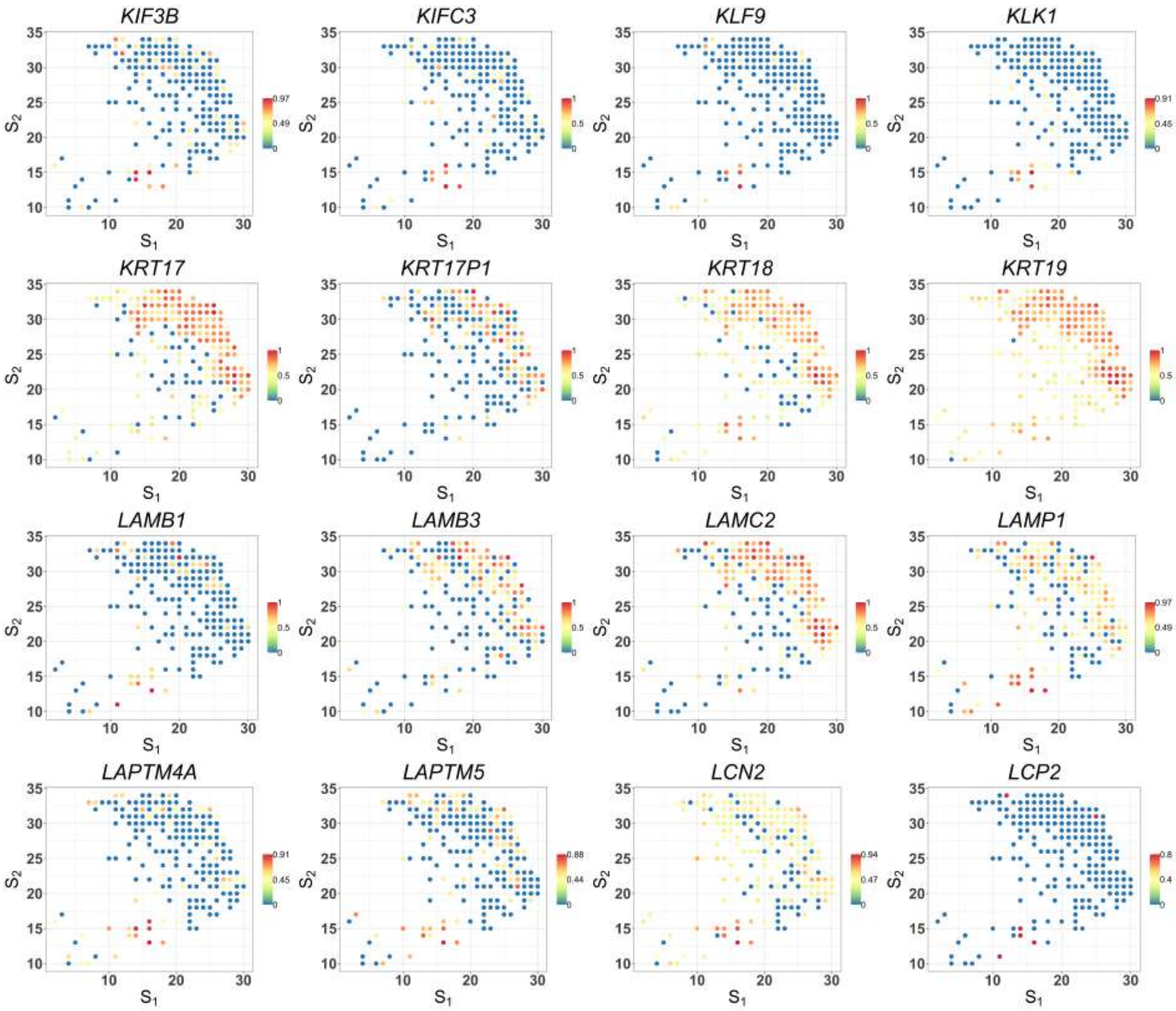


Figure S8: Spatial expression patterns of cancer-cell-specific SV genes detected by SPARK-X in the cancer region of PDAC data (part 36). Values are relative expressions.

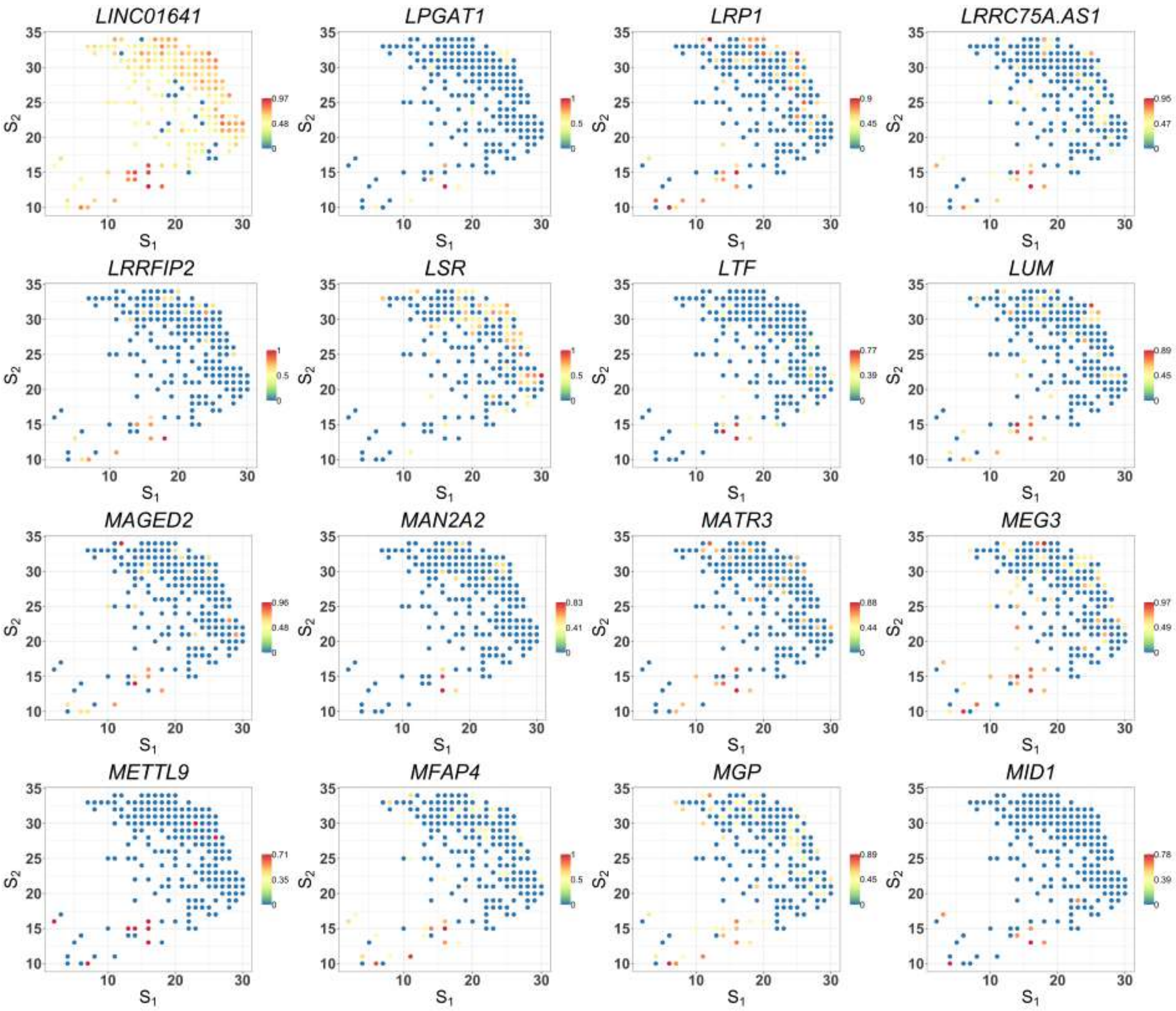


Figure S8: Spatial expression patterns of cancer-cell-specific SV genes detected by SPARK-X in the cancer region of PDAC data (part 37). Values are relative expressions.

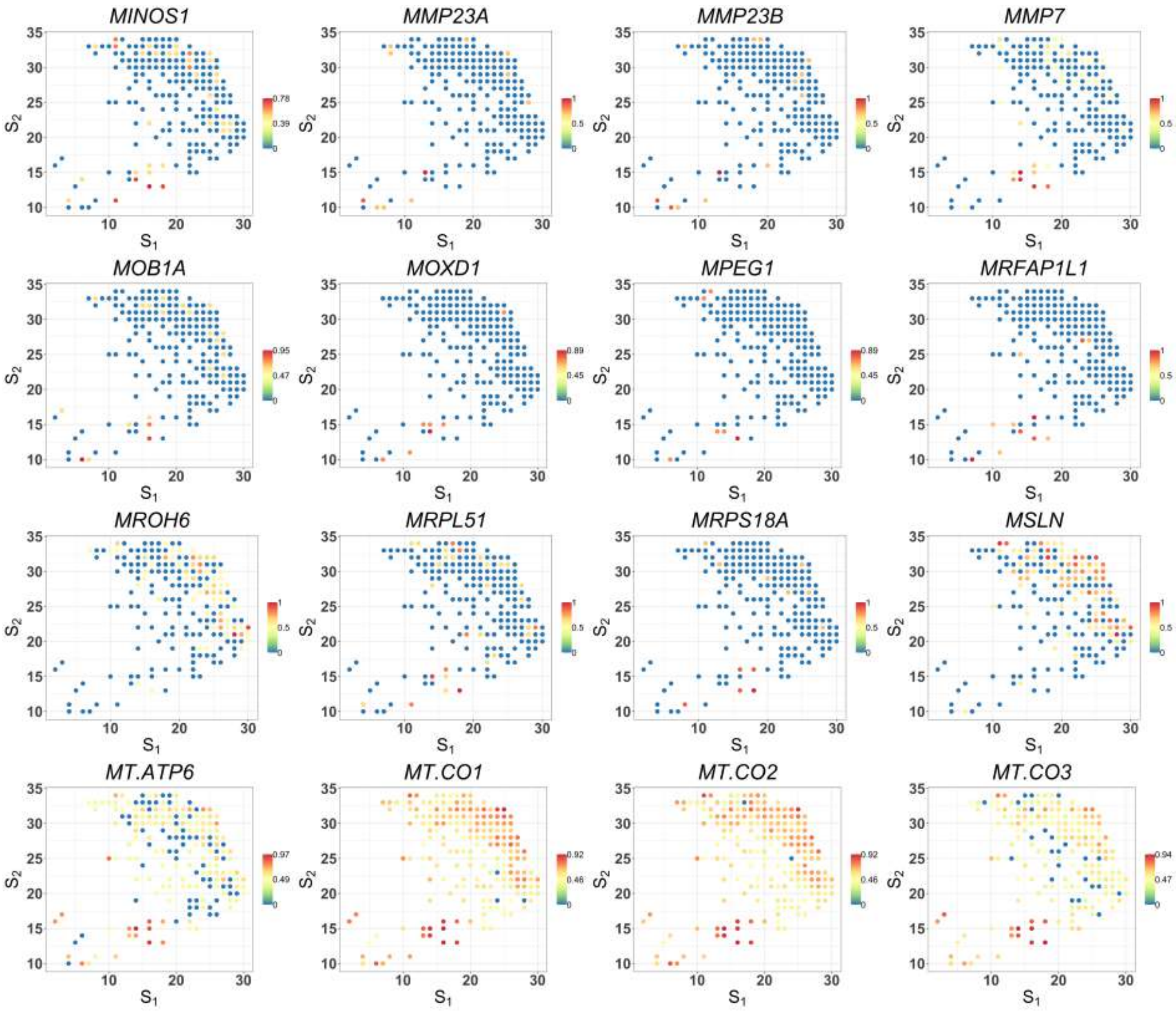


Figure S8: Spatial expression patterns of cancer-cell-specific SV genes detected by SPARK-X in the cancer region of PDAC data (part 38). Values are relative expressions.

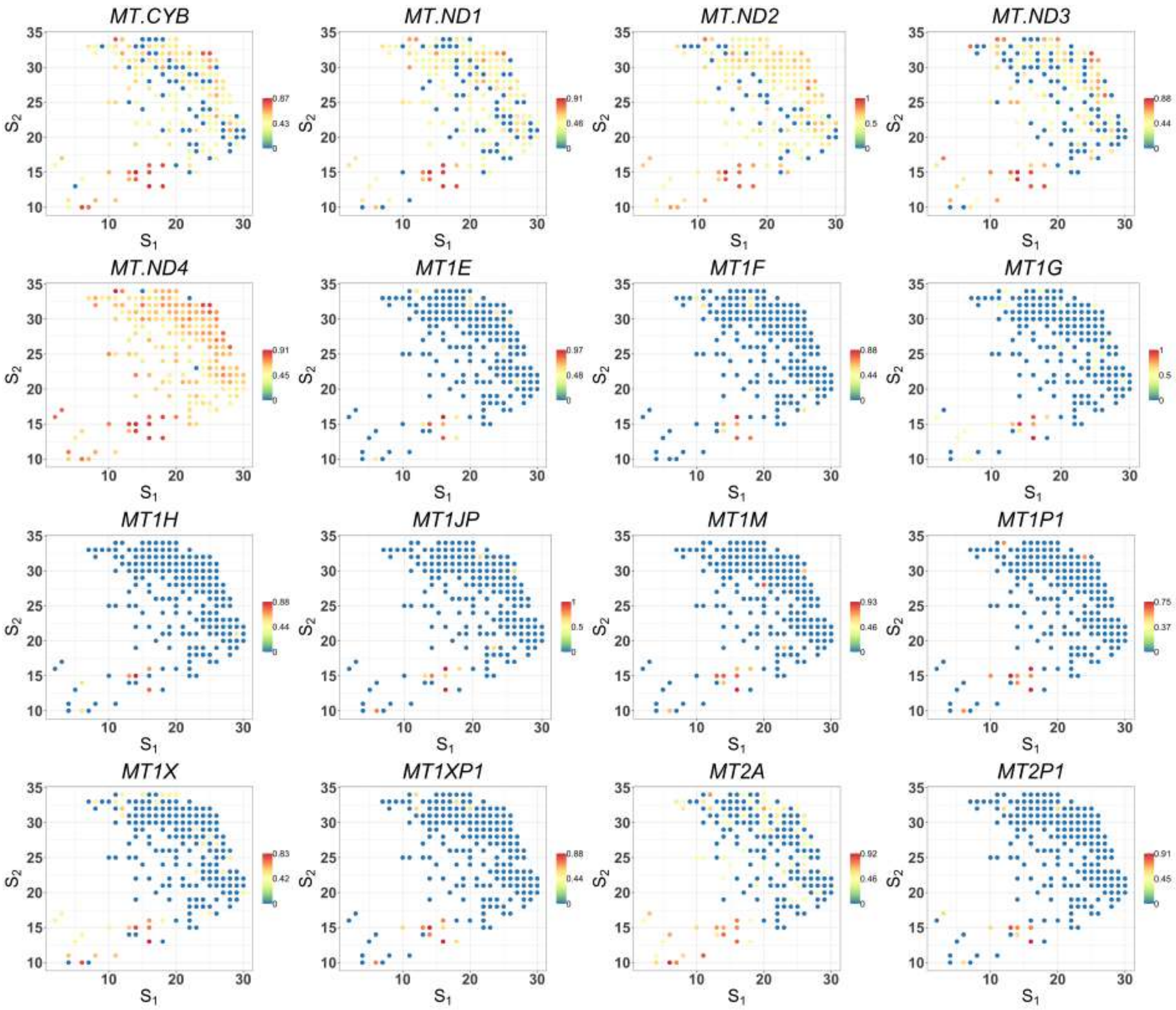


Figure S8: Spatial expression patterns of cancer-cell-specific SV genes detected by SPARK-X in the cancer region of PDAC data (part 39). Values are relative expressions.

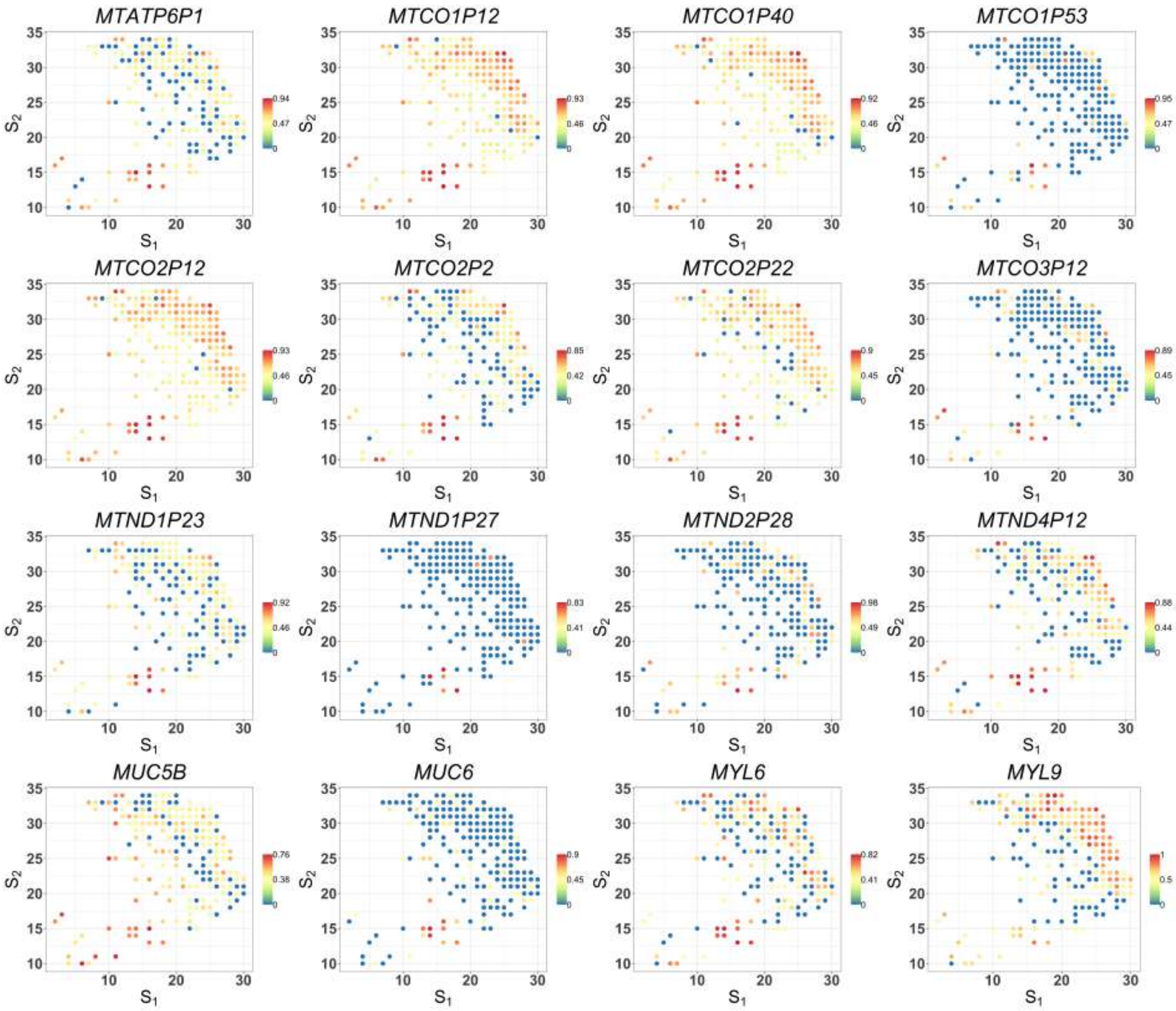


Figure S8: Spatial expression patterns of cancer-cell-specific SV genes detected by SPARK-X in the cancer region of PDAC data (part 40). Values are relative expressions.

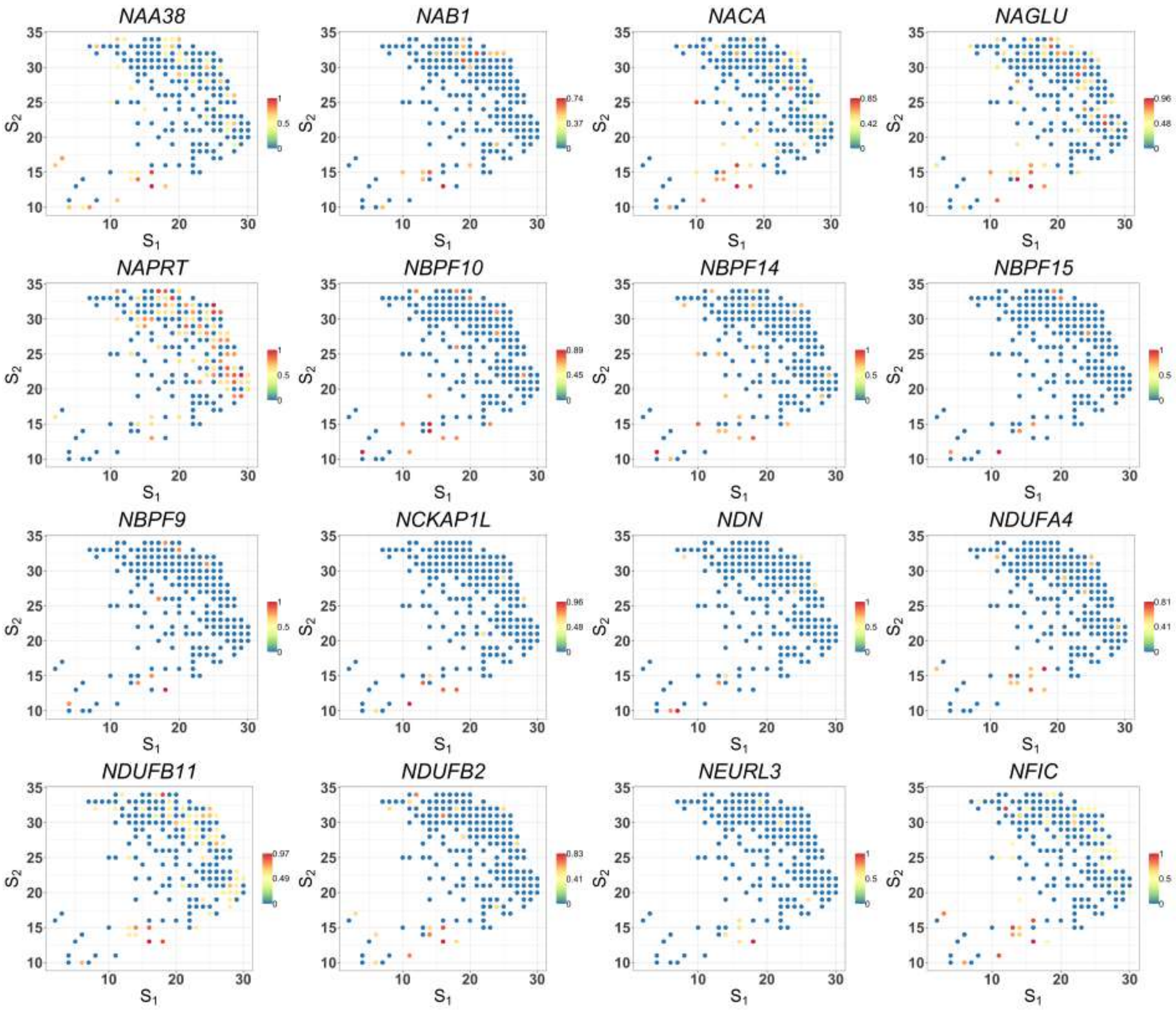


Figure S8: Spatial expression patterns of cancer-cell-specific SV genes detected by SPARK-X in the cancer region of PDAC data (part 41). Values are relative expressions.

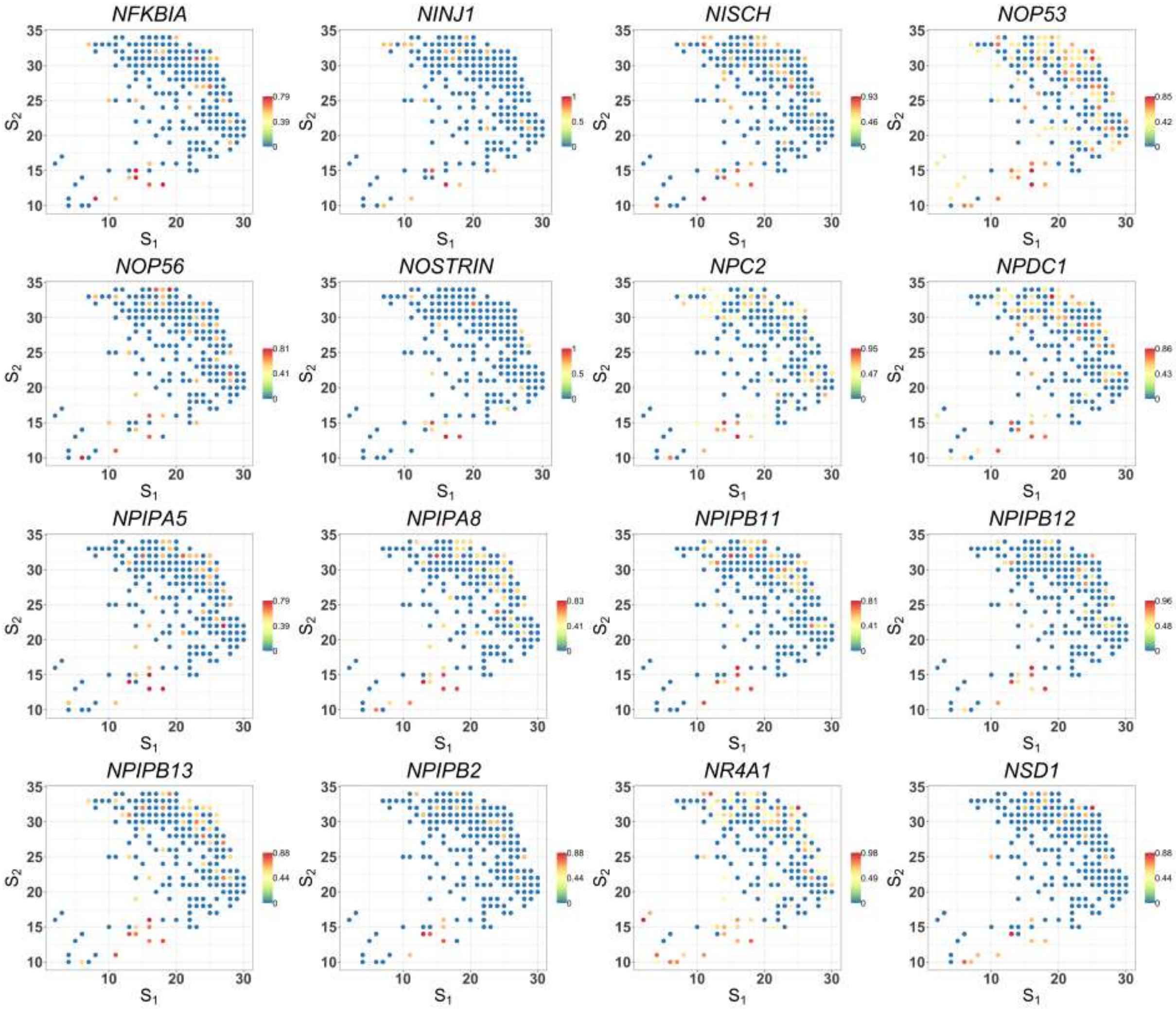


Figure S8: Spatial expression patterns of cancer-cell-specific SV genes detected by SPARK-X in the cancer region of PDAC data (part 42). Values are relative expressions.

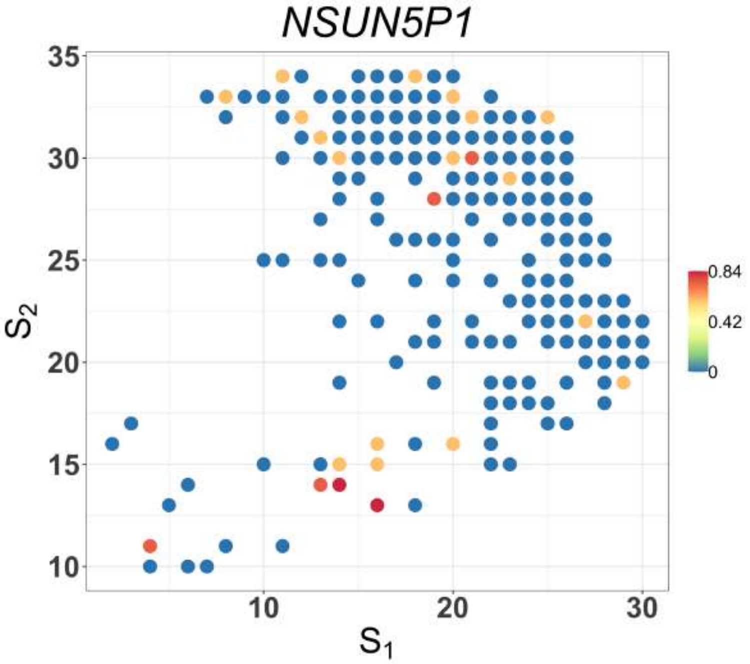


Figure S8: Spatial expression patterns of cancer-cell-specific SV genes detected by SPARK-X in the cancer region of PDAC data (part 43). Values are relative expressions.

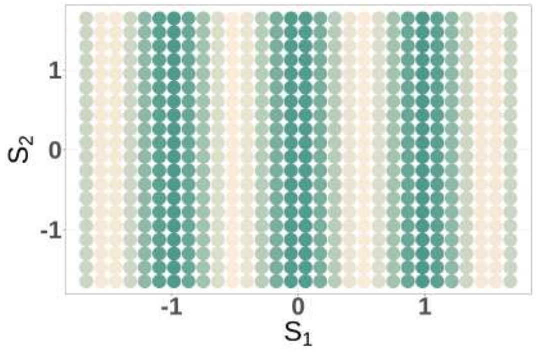


Figure S9: An example of the SV gene expression pattern where only $s_{i1}$ is associated with expressions.
